# Mitochondrial defects result in decreased susceptibility to echinocandins via the transcriptional regulator Pdr1 in *Candida glabrata*

**DOI:** 10.1101/2025.03.25.645289

**Authors:** Natalia Klimova, Hetaben Patel, Frédéric Devaux, Bernard Turcotte

## Abstract

In the human fungal pathogen *Candida glabrata*, the transcription factor Pdr1 controls the expression of genes encoding drug efflux pumps such as Cdr1. Pdr1 also controls its own expression by binding to pleiotropic drug responsive elements (PDREs) located in its promoter. Increased resistance to the antifungal drugs azoles (e.g. fluconazole) is often due to gain-of-function mutations in *PDR1* that render the factor hyperactive. Mitochondrial defects also result in increased resistance to azoles via Pdr1. Resistance to another class of antifungals, the echinocandins (e.g. micafungin), is generally due to mutations in the *FKS1* and *FKS2* genes that encode the catalytic subunit of 1,3-beta-D-glucan synthase, the echinocandin target. We have observed that mitochondrial defects also result in increased resistance to echinocandins and we have shown that this process is mediated by Pdr1. Mitochondrial defects result in increased levels of *PDR1* mRNA. Overexpression of *PDR1* by replacing its native promoter by the strong *ADH1* promoter resulted in increased resistance to micafungin. However, mutations in *PDR1* that decrease susceptibility to azoles generally have modest effects on echinocandin resistance. We randomly mutagenized the *PDR1* gene and screened for mutants with altered resistance to micafungin. Single amino acid changes downstream of the DNA binding domain result in preferential resistance to micafungin as compared to fluconazole. We also show that auto-regulation of *PDR1* expression is necessary for resistance to micafungin. One *PDR1* mutant showed over 15-fold increased promoter activity in a PDRE-dependent manner. In summary, we have identified a novel role for the transcriptional regulator Pdr1.

## INTRODUCTION

In humans, fungi are responsible for approximately 1.4 million deaths each year (Brown *et al*., 2012). Infections by pathogenic fungi are becoming more widespread, especially in immunosuppressed patients. *Candida glabrata* now ranks as the second most important human fungal pathogen after *C. albicans* (reviewed in (Whaley and Rogers, 2016; Galocha *et al*., 2019)). Despite its formal name, *C. glabrata* is more closely related to the non-pathogenic baker’s yeast *Saccharomyces cerevisiae* (Dujon *et al*., 2004). There are various classes of antifungal drugs but the most currently used are azoles and echinocandins that interfere with ergosterol and β-glucan synthesis, respectively. Ergosterol is related to cholesterol and is an important component of the fungal plasma membrane. Azoles (e.g. fluconazole, ketoconazole) can be used orally or intravenously to treat infections but these drugs are only fungistatic. Azoles target lanosterol 14α-demethylase involved in the synthesis of ergosterol and this enzyme is encoded by the *ERG11* gene. Antifungal activity is due to decreased ergosterol levels and increased production of toxic ergosterol derivatives (Lupetti *et al*., 2002).

Resistance to azoles is often due to mutations in the *PDR1* gene that belongs to a large family of transcriptional regulators, the zinc cluster proteins (reviewed in (MacPherson *et al*., 2006; Whaley and Rogers, 2016; Moye-Rowley, 2019)). *C. glabrata* Pdr1, the homologue of *S. cerevisiae* Pdr1/Pdr3, has been shown to confer drug resistance by positively controlling the expression of various genes including the ABC transporters *CDR1*, *PDH1*, and *SNQ2* that act as drug efflux pumps (Vermitsky and Edlind, 2004; Vermitsky *et al*., 2006). Direct binding of various compounds (including azoles) to Pdr1 activates this transcription factor (Thakur *et al*., 2008). Pdr1 binds to pleiotropic drug response elements (PDREs) present in promoters of target genes for regulation of gene expression (Paul *et al*., 2011). The *PDR1* promoter also contains two PDREs involved in Pdr1 positive autoregulation which is essential for resistance to azoles (Paul *et al*., 2011). Overexpression of *PDR1* by replacing its promoter by a stronger one increases resistance to azoles (Paul *et al*., 2011). Moreover, mutations in the *PDR1* gene result in hyperactivation of the transcription factor causing increased resistance to various drugs such as azoles and, unexpectedly, increased virulence (Tsai *et al*., 2006; Vermitsky *et al*., 2006; Ferrari *et al*., 2009; Berila *et al*., 2009). Although Pdr3 is absent in *C. glabrata*, recent observations suggest that *C. glabrata* Pdr1 is a hybrid molecule between *S. cerevisiae* Pdr1 and Pdr3 (Khakhina *et al*., 2018). In *C. glabrata*, mitochondrial dysfunction results in the activation of Pdr1 and increased resistance to azoles while in *S. cerevisiae*, activation of Pdr3, but not Pdr1, is observed with a mitochondrial defect (Sanglard *et al*., 2001; Vermitsky and Edlind, 2004; Tsai *et al*., 2006). Thus, *PDR1* plays a pivotal role in modulating resistance to azoles.

Echinocandins (caspofungin, micafungin and anidulafungin) are the latest class of antifungal drugs and, in contrast to azoles, show fungicidal activity (reviewed in (Chen *et al*., 2011; Healey and Perlin, 2018)). They are the drugs of choice to treat *C*. *glabrata* infections, but they must be administered intravenously. Echinocandins inhibit the activity of a two-subunit enzyme involved in the synthesis of the polysaccharide 1,3-ß-D-glucan, a major and essential component of the cell wall. *FKS1* and *FKS2* are two redundant genes that encode the subunit with ß-glucan synthase activity (Katiyar *et al*., 2012:1) while *RHO1* encode the second subunit. Rho1 is a GTP-binding protein that controls the activity of the complex (Mazur and Baginsky, 1996). Resistance to echinocandins is due to mutations in hot-spots of the *FKS1* and *FKS2* genes (Perlin, 2015). Overexpression of drug efflux pumps in *S. cerevisiae* does not significantly affect susceptibility to echinocandins (Niimi *et al*., 2006). However, this study in *C. glabrata* shows a link between echinocandins and *PDR1*. We show that mitochondrial defects result in increased resistance to echinocandin via the transcription factor Pdr1.

## MATERIAL AND METHODS

### Strains

Strains used in this study are listed in Table 1. Rho^-^ strains were generated according to Fox *et al*. (1991) except that after the first overnight culture, cells were diluted approximately 100-fold and grown again overnight in minimal medium containing 25 µg/ ml ethidium bromide. Single colonies were grown in YPD and spotted on YEP supplemented with 2% ethanol. Strains unable to grow on ethanol plates were selected. The rho^-^ phenotype was confirmed by isolating mitochondrial DNA and performing qPCR for the mitochondrial genes *ATP9* (oligos CgATP9-1 and CgATP9-2) and *COX1* (oligos CgCOX1-1 and CgCOX1-2) where gene alteration was observed for at least one of these genes (data not shown). Strains CG292 and CG293, overexpressing the *PDR1* coding region, were obtained by replacing the native *PDR1* promoter with the strong *S. cerevisiae ADH1* promoter. Oligos Tc-CgZCF1-1 and Tc-CgZCF1-2 were used to amplify the ‘*URA3*-*ADH1*=promoter-3X aptamers’ region of a plasmid derived from pADH1-tc3-3xHA and carrying a *URA3* marker (Klimova *et al*., 2014). This was followed by further extension of sequences homologous to *PDR1* by a second amplification of the initial PCR product with oligos Tc-CgZCF1-3 and Tc-CgZCF1-4. The final PCR product was transformed into strain 66032 followed by selection for the *URA3* marker. Strains CG197 and CG198: the MYC-URA3-MYC region of plasmid pMPY-3XMYC (Schneider *et al*., 1995) was amplified using oligos PET-

**Table 1.** Strains used in this study.

| Strain | Parent strain | Gene | Tag/ gene | Reference |
| --- | --- | --- | --- | --- |
| 66032 | N/A | *ura3* | - | (Vermitsky *et al.*, 2006) |
| CBS138 | N/A | - | G418<sup>R</sup> | ATCC2001 |
| CG197 | 66032 | $\Delta pdr1$ | 3XMyC-URA3-3XMyC | This study |
| CG198 | 66032 | $\Delta pdr1$ | 3XMyC-URA3-3XMyC | This study |
| CG252 | 66032 | *S. cerevisiae*<br>*URA3* | *C. glabrata ura3* replaced<br>by *S. cerevisiae URA3* | This study |
| CG253 | 66032 | *URA3* | *C. glabrata ura3* replaced<br>by *S. cerevisiae URA3* | This study |
| CG266 | CG198 | $\Delta pdr1$ rho<sup>-</sup> | treated with ethidium<br>bromide to induce loss of<br>Mito DNA | This study |
| CG267 | CG198 | $\Delta pdr1$ rho<sup>-</sup> | treated with ethidium<br>bromide to induce loss of<br>Mito DNA | This study |
| CG268 | CBS138 | rho<sup>-</sup> | treated with ethidium<br>bromide to induce loss of<br>Mito DNA | This study |
| CG269 | CBS138 | rho<sup>-</sup> | treated with ethidium<br>bromide to induce loss of<br>Mito DNA | This study |
| CG291 | 66032 | ADH-3TC-3HA-PDR1 | *S. cerevisiae ADH1*<br>promoter controlling<br>expression *PDR1*<br>(translation *PDR1*<br>repressible by tetracycline | This study |
| CG292 | 66032 | ADH-3TC-3HA-PDR1 | *S. cerevisiae ADH1*<br>promoter controlling<br>expression *PDR1*<br>(translation *PDR1*<br>repressible by tetracycline | This study |
| CG295 | CG252 | rho<sup>-</sup> | treated with ethidium<br>bromide to induce loss of<br>mito DNA | This study |
| CG296 | CG252 | rho<sup>-</sup> | treated with ethidium<br>bromide to induce loss of<br>mito DNA | This study |
| CG298 | CG253 | Sc*URA3* rho<sup>-</sup> | treated with ethidium<br>bromide to induce loss of<br>mito DNA | This study |
| CG299 | CG253 | ScURA3 rho<sup>-</sup> | treated with ethidium bromide to induce loss of mito DNA | This study |
| CG301 | CG198 | *PDR1*- L328F | Mutation integrated at the natural *PDR1* | This study |
| CG302 | CG198 | *PDR1*- L328F | Mutation integrated at the natural *PDR1* | This study |
| CG304 | CG198 | *PDR1*- P822L | Mutation integrated at the natural *PDR1* | This study |
| CG305 | CG198 | *PDR1*- P822L | Mutation integrated at the natural *PDR1* | This study |
| CG306 | CG198 | *PDR1*-T588A | Mutation integrated at the natural *PDR1* | This study |
| CG307 | CG198 | *PDR1*-T588A | Mutation integrated at the natural *PDR1* | This study |
| CG350 | CG253 | *PDR1*-Y256R | Mutation integrated at the natural *PDR1* | This study |
| CG351 | CG253 | *PDR1*-Y256R | Mutation integrated at the natural *PDR1* | This study |
| CG354 | CG253 | *PDR1*-D261G | Mutation integrated at the natural *PDR1* | This study |
| CG355 | CG253 | *PDR1*-D261G | Mutation integrated at the natural *PDR1* | This study |
| CG360 | CG253 | *PDR1*-Y208C | Mutation integrated at the natural *PDR1* | This study |
| CG361 | CG253 | *PDR1*-Y208C | Mutation integrated at the natural *PDR1* | This study |

**Table 2.**
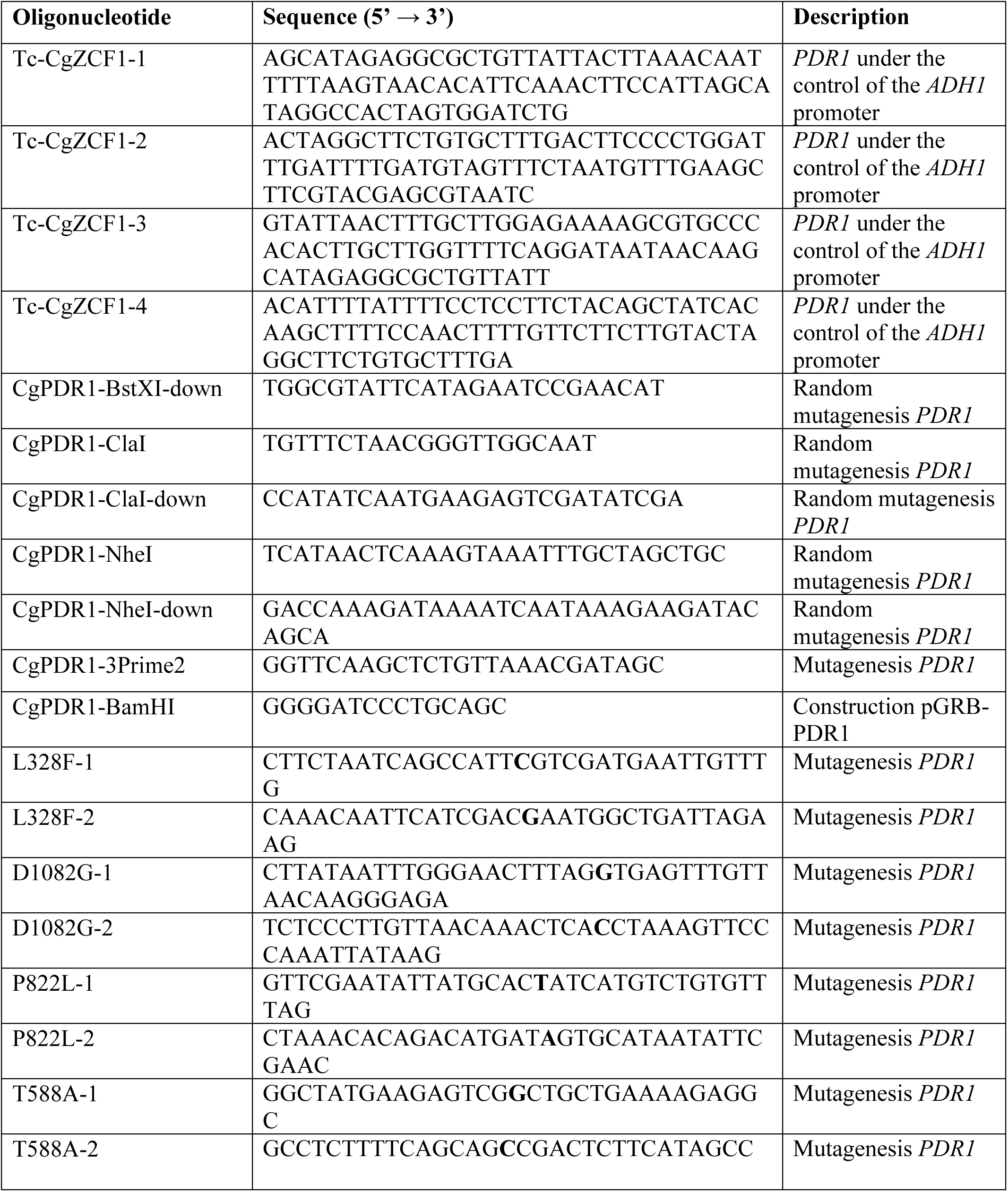
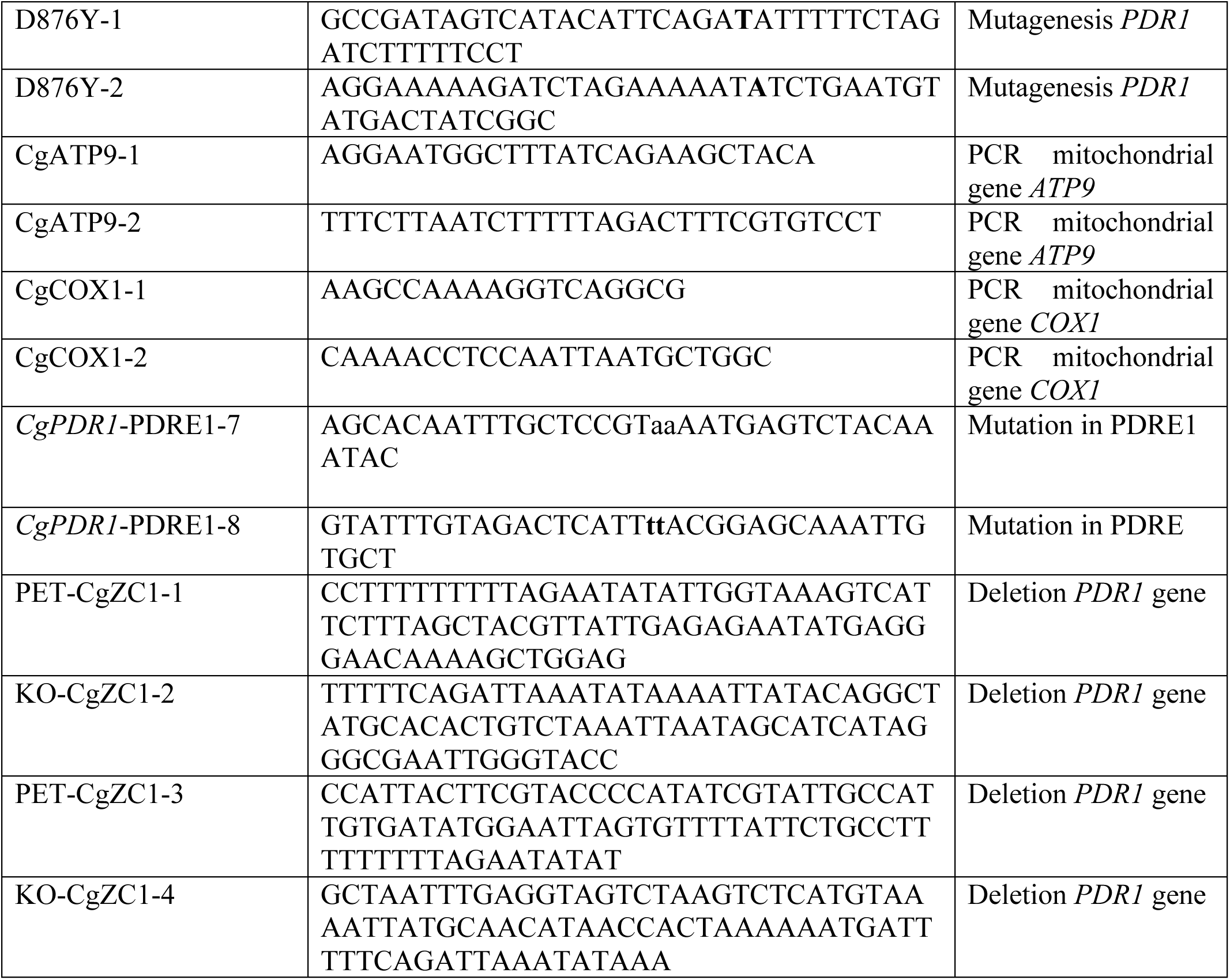
Oligonucleotides used in this study.

CgZC1 + KO-CgZC1-2 followed by extension of the *PDR1* region of homology by amplification of the first PCR product using oligos PET-CgZC1-3 and KO-CgZC1-4. The PCR product was transformed into strain 66032 followed by selection for *URA3*. The URA3 marker was removed by selection on FOA plates as described (Schneider *et al*., 1995). For strains CG350, CG351, CG360, CG361, CG354, and CG355, plasmids pGRB-PDR1-Y208C, pGRB-PDR1-Y256R and pGRB-PDR1-D261G were cut with SacI and SmaI and transformed into strain CG252 followed by selection on 5-fluoroootic acid (FOA) for introduction of the desired mutations at the native *PDR1* locus.

### Plasmids and random mutagenesis

Plasmids pPDR1-C, pGRB-PDR1, pPDR1-Y208C and pPDR1-Y208C-mPDRE were constructed in multiple steps using pGRB2.1 as a starting plasmid and genomic DNA from strain 66032 as template for PCR of the *PDR1* gene. Similarly, pPDR1-lacZ and pPDR1-lacZmPDRE were constructed in multiple steps. Sequencing of the final constructs was performed by Plasmidsaurus using Oxford Nanopore Technology with custom analysis and annotation. The plasmid DNA sequences and maps generated using the software Snapgene are provided in Supplemental Fig. S1. Indels and SNPs were found in the *PDR1* promoter derived from the 66032 strain as compared to the reference genome CBS138 (Dujon *et al*., 2004). Plasmid pPDR1-C pGRB-PDR1 have almost identical sequences with differences lying in the plasmid backbone. Both plasmids contain the *PDR1* promoter, its coding region and the 3’UTR and carry an *URA3* marker and a chimeric CEN/ARS sequence. pPDR1-Y208C is identical to pGRB-PDR1 except that it contains a mutation changing tyrosine to cysteine at amino acid 208. Plasmid pPDR1-Y208C-mPDRE is identical to pPDR1-Y208C except that both PDREs have been mutated (PDRE1 ‘TCCACGGA’ to ‘TaaaaaaA’; PDRE2 ‘TCCGTGGA’ to ‘TaaaaaaA’).

For random mutagenesis, 3 different fragments parts of the *PDR1* ORF were amplified using pPDR1-C as a template. Thirty-five cycles for amplification cycles were performed using Taq DNA polymerase. Oligos CgPDR1-BstXI-down and CgPDR1-ClaI were used to amplify the 5’ part of the *PDR1* ORF and the PCR product was cut with BstXI and ClaI and subcloned in pPDR1-C cut with the same enzymes. For the middle part, oligos CgPDR1-ClaI down and CgPDR1-NheI were used. The PCR product was cut with ClaI and NheI and subcloned into pPDR1-C cut with the same enzymes. Finally, random mutagenesis of the 3’ part of the *PDR1* ORF was performed similarly using oligos Cg-NheI-down and M13 reverse with enzymes NheI and EcoRI for subcloning. For PDR1 mutants Y208C, Y256R and D261G, two PCR products encompassing the site to be mutated were introduced into pGRB-PDR1 cut with PflMI and AvrII by Gibson assembly to yield pGRB-PDR1-Y208C, pGRB-PDR1-Y256R and pGRB-PDR1-D261G.

### Βeta-galactosidase assays

Reporters pPDR1-lacZ and pPDR1-lacZmPDRE were transformed into strains 66032 and CG360. Βeta-galactosidase were performed with permeabilized cells using ortho-nitrophenyl-D-galactopyranoside (CPRG) as a substrate at a final concentration of 1 mM.

## RESULTS

Spotting assays with a wild-type *C. glabrata* strain on plates containing micafungin typically revealed the appearance of multiple resistant colonies (Fig. 1A). Since resistant colonies were obtained at a relatively high frequency (approximately 0.1%), we reasoned that this phenotype is unlikely be due to mutations in the *FKS1* or *FKS2* genes but may be due to mitochondrial dysfunction, in analogy to azoles where mitochondrial defects result in resistance to these drugs (Sanglard *et al*., 2001; Vermitsky and Edlind, 2004; Tsai *et al*., 2006). Indeed, our results show that resistant strains are respiratory deficient since no growth was observed when spotting resistant strains on plates containing ethanol as the sole carbon source (Fig. 1A, right panel), an observation consistent with a mitochondrial defect. To directly test this hypothesis, we generated rho^-^ strains (i.e. cells lacking in part mitochondrial DNA) via treatment with ethidium bromide. These strains were resistant to micafungin (Fig. 1B). We determined if these observations also apply to another echinocandin, caspofungin, by measuring minimal inhibitory concentration (MIC) with a wild-type strain and 2 independent rho^-^ strains. Results show increased MIC with both rho^-^ stains as compared to the wild-type strain (Fig. 2, top panel). We also tested a strain with a different background. Results show that resistance to micafungin is increased when rendering strain CBS138 rho^-^ (Fig. 2, bottom panel) as well as 2 other strains. Thus, our observations strongly suggest that increased resistance to echinocandins due to mitochondrial defects is not a strain-specific phenomenon.

**Fig. 1.**
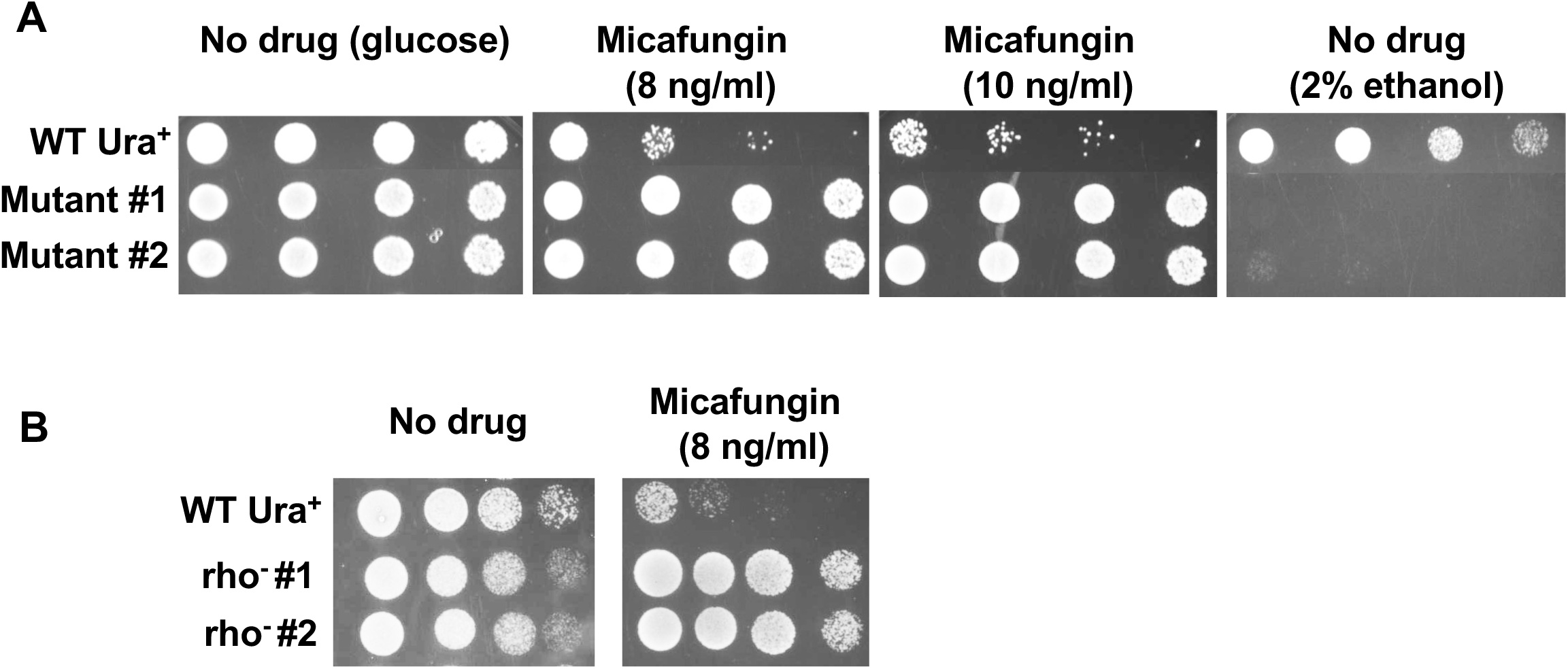
Spontaneous mutants showing resistance to micafungin are respiratory-deficient. A) Cultures of wild-type *C. glabrata* strain (CG252) and two spontaneous mutants showing decreased susceptibility to micafungin were serially diluted and spotted on plates containing 2% glucose and 0, 8, and 10 ng/ml micafungin or without micafungin and 2% ethanol as the sole carbon source as indicated on top of the Fig. Absence of growth on plates containing ethanol as a carbon source indicates deficiency in respiration. B) Two independent rho^-^ clones (CGG295 and CG296) were generated from wild-type strain CG252 and tested for sensitivity to micafungin.

**Fig. 2.**
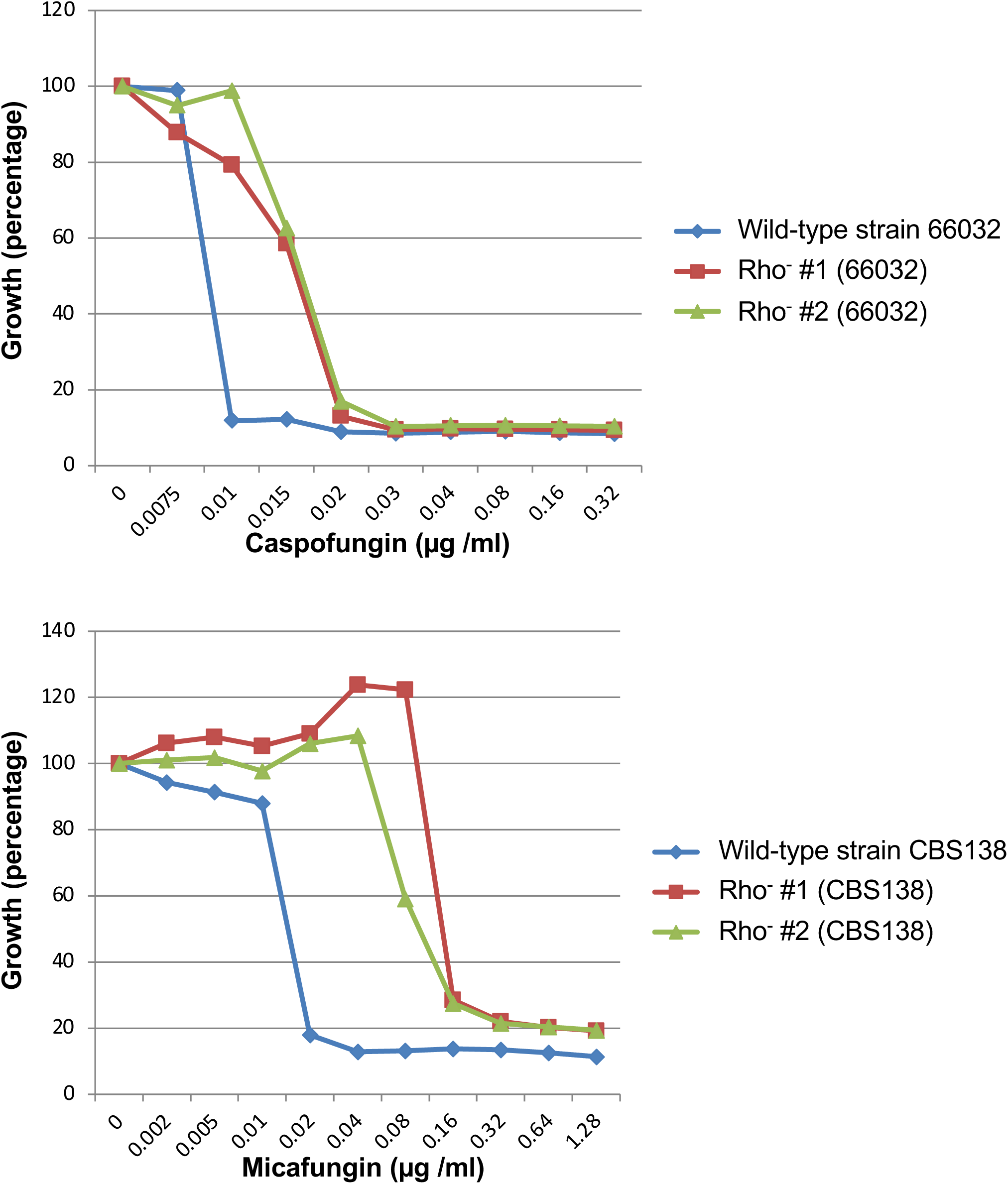
Rho^-^ strains (background 66032) are less susceptible to caspofungin (top panel) while rho^-^ strains (background CBS138) are less susceptible to micafungin (bottom panel). MIC were determined as described in Material and Methods. Top panel: caspofungin MIC for strain CG252 and rho^-^ (CG295, CG296) derivatives. Bottom panel: micafungin MIC for strain CBS138 and rho^-^ derivatives (CG268 and CG269).

Since *PDR1* was shown to be required to confer azole resistance in strains with mitochondrial defects, we hypothesize that it could also be a mediator of echinocandin resistance. As described above, rho^-^ strains are more resistant to micafungin (Fig. 3A). However, deletion of *PDR1* in a rho^-^ strain drastically decreased resistance (Fig 3A). Next, we wanted to test if it is possible to bypass the requirement for a mitochondrial defect to obtain resistance to micafungin by overexpressing *PDR1*. To this end, we constructed strains in which the native *PDR1* promoter was replaced by the strong *ADH1* promoter from *S. cerevisiae* by homologous recombination. As expected from a previous study (Khakhina *et al*., 2018), overexpression of *PDR1* results in increased resistance to fluconazole while deletion of this gene greatly increased sensitivity to fluconazole as compared to a wild-type strain (Fig. 3B, bottom panel). Regarding micafungin, overexpression of *PDR1* in a rho^+^ strain resulted in increased resistance to micafungin (Fig. 3B, top panel). Thus, these results show that overexpression of *PDR1* bypasses the requirement for mitochondrial defect for enhanced resistance to micafungin. Results also suggest that mitochondrial defects somehow activate Pdr1 for regulation of gene(s) involved in modulating resistance to echinocandins.

**Fig. 3.**
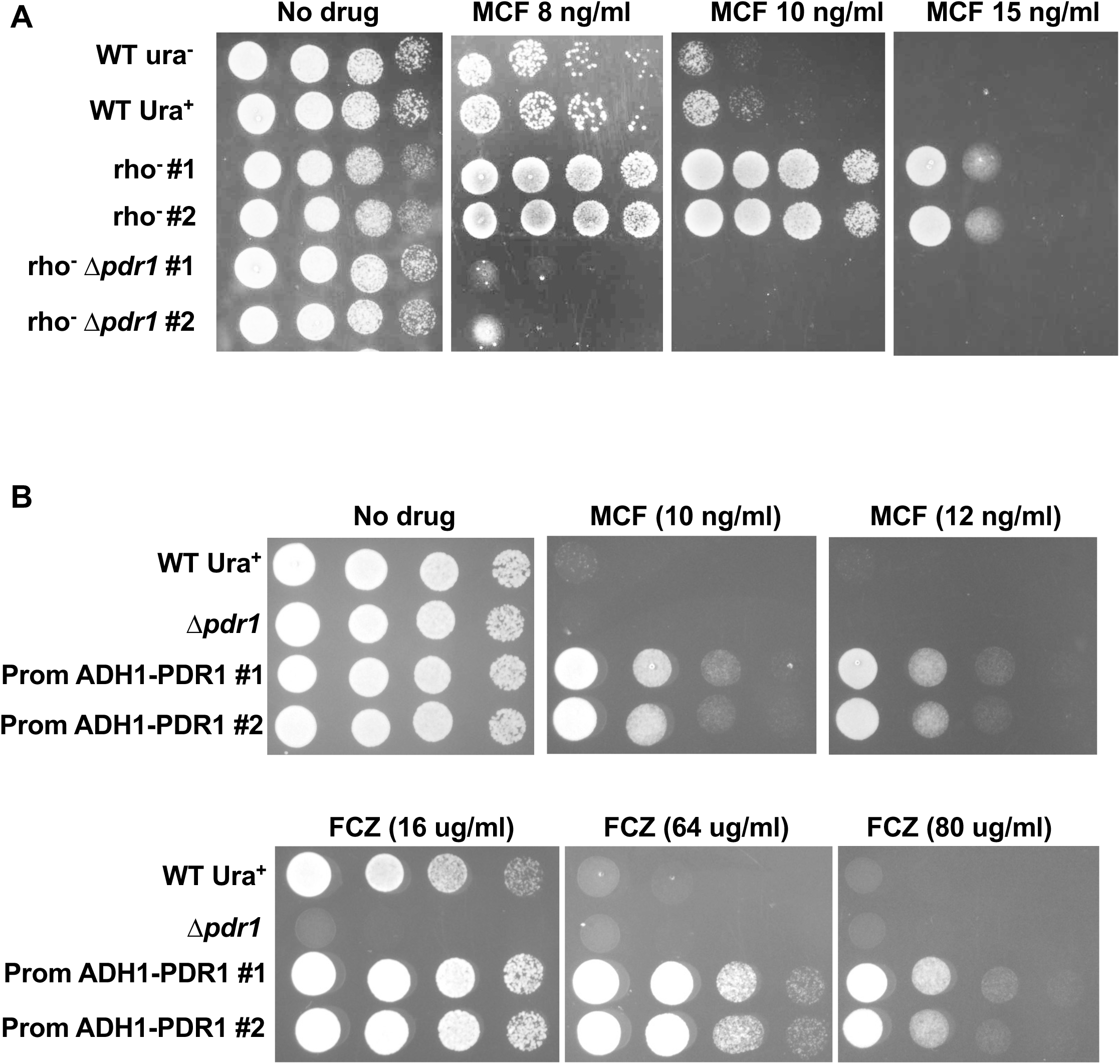
Pdr1 mediates resistance to micafungin and its overexpression bypasses the requirement for a mitochondrial defect. Cells were grown overnight in rich medium, serially diluted and spotted on plates containing micafungin (MCF) at the indicated concentration. A) Deletion of the *PDR1*gene in a rho^-^ background abolishes resistance to MCF. Spotting assays were performed with wild-type strain CG252, rho^-^ derivatives (strains CG295 and CG296), and rho^-^ strains carrying a deletion of the *PDR1* gene (strains CG266 and CG267). Concentration of MCF is indicated at the top of the panels. B) Overexpression of *PDR1*with the strong *ADH1* promoter confers resistance to both MCF and fluconazole (FCZ) in the wild-type strain 66032. Strains (from top to bottom of panels) CG253, CG198, CG291, and CG292 were used for spotting assays diluted and spotted on plates containing 0, 10 and 12 ng/ ml micafungin (‘MCF’) Top panel) or 16, 64 and 80 µg/ ml fluconazole (‘FCZ’) (bottom panel).

As stated above, some mutations in the coding region of the *PDR1* gene result in a hyperactive transcription factor leading to increased expression of target genes including those encoding drug efflux pumps resulting in increased resistance to azoles. We were interested in determining if mutations in the *PDR1* that increase resistance to azoles also modulate resistance to micafungin. Using clinical isolates showing resistance to azoles, the Sanglard’s group has identified a number of mutations in the *PDR1* gene that are responsible for this phenotype (Ferrari *et al*., 2009). We selected three gain-of-function mutations (L328F, P822L and T558A) that were introduced at the native *PDR1* locus by homologous recombination. As expected (Ferrari *et al*., 2009), these mutants showed greatly increased resistance to fluconazole as compared to a wild-type strain (Fig. 4, top panel). Greatly reduced growth was observed in a strain lacking the *PDR1* gene while a Δ*pdr1* rho^-^ strain was sensitive to fluconazole. Regarding micafungin, the mutants showed some increased resistance to this drug (Fig. 4, bottom panel). However, they were less resistant than a rho^-^ strain. For example, with 12 ng micafungin per ml, greatly reduced growth was observed for mutants P822L and T558A as compared to a rho^-^ strain. Similar results were obtained with *PDR1* mutants D1082G and D876Y (data not shown). These results may be explained by the fact that these mutants derived from clinical isolates were ‘selected’ for resistance to azoles but not micafungin. In these mutants, changes in gene expression may not result in altered expression of some Pdr1 targets involved in conferring resistance to echinocandins. In agreement with this hypothesis, a study showed that *PDR1* mutants alter expression of a common set of genes as well as genes only regulated by a specific mutation (Caudle *et al*., 2011).

**Fig. 4.**
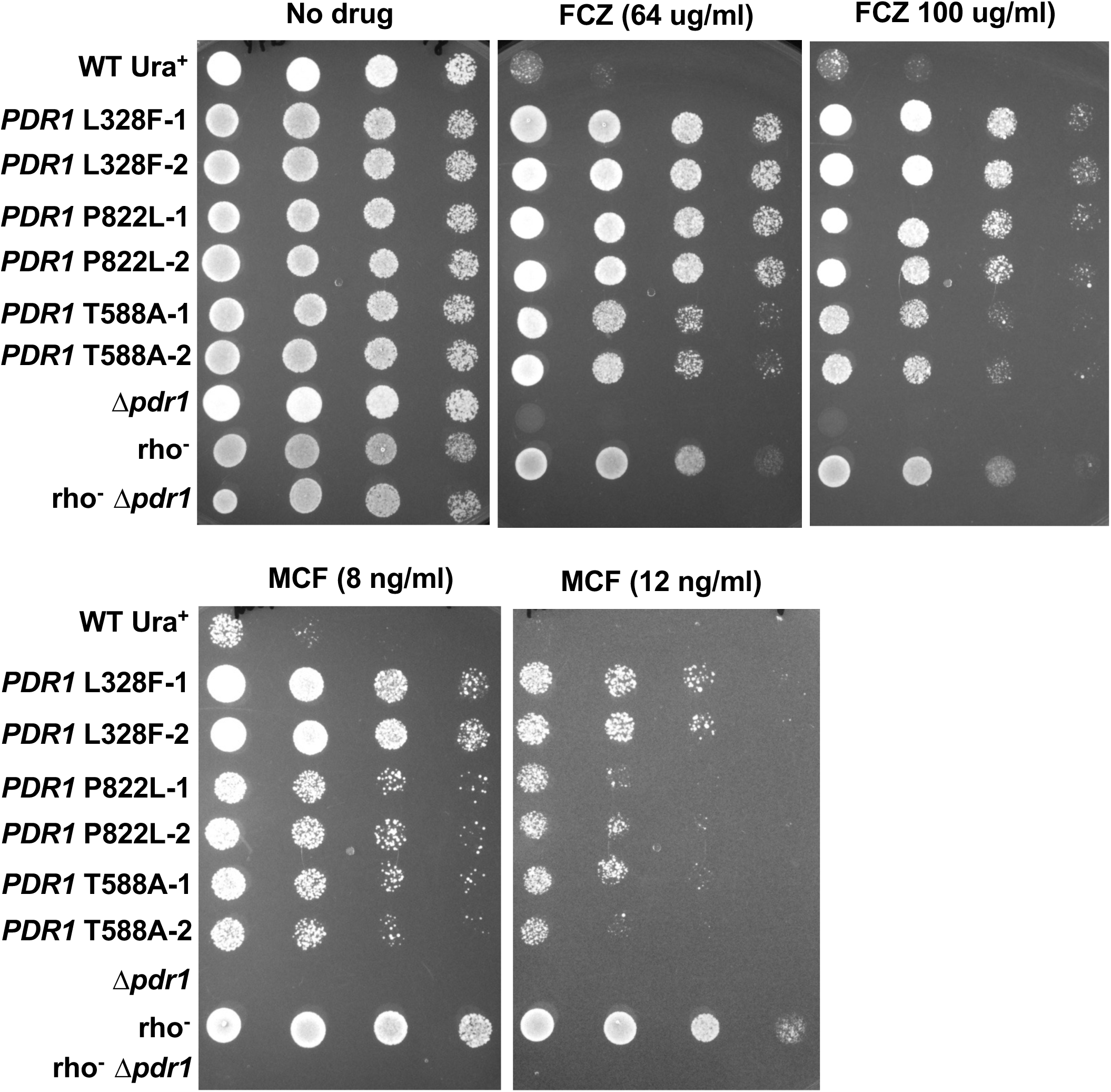
*PDR1* mutants derived from clinical isolates resistant to micafungin (MCF) show less resistance to micafungin (MCF) as compared to a rho^-^ strain. Spotting assays were performed with 2 independent integrants (‘-1’ and ‘-2’) for the various *PDR1* mutants tested using strains (from top to bottom of the panels) CG253, CG301, CG302, CG304, CG305, CG306, CG307. CG198, CG298, and CG267.

We were interested to determine if some *PDR1* mutants would preferentially confer increased resistance to micafungin as compared to the ones tested above. To this end, we randomly mutagenized the *PDR1* gene. Three PCR products corresponding to the 5’, middle and 3’ parts of the coding region were generated and subcloned in a plasmid containing the entire *PDR1* gene as well as a *C. glabrata* origin of replication and the *URA3* selective marker. Plasmids were transformed into a Δ*pdr1* strain and selected for resistance to micafungin. A total of approximately 25 clones were obtained and they were all rho**^+^**. For all *PDR1* mutants, the mutagenized region was from the 5’ part of the coding region. The phenotypes were confirmed by transferring the plasmids to *E. coli* and retransformed into *C. glabrata*. We focused on 3 mutants: RM2, RM7 and RM8. DNA sequencing showed that the mutants carry multiple non-silent and silent mutations. In order to map the mutation(s) responsible for increased resistance, we made hybrid molecules using wild-type *PDR1* sequences and mutants. Phenotypes could be assigned to changes Y256R, Y208C and D261G for mutants RM1, RM2, and RM7, respectively. Fig. 5A shows that these mutations were located next to the DNA binding domain in the middle homology region (MHR) involved in controlling activity of the transcription factor (Khakhina *et al*., 2018). Results were confirmed by introducing these mutations at the *PDR1* locus by homologous recombination (Fig. 5B). For example, with mutants Y256R and D261G, solid growth was observed at 18 ng/ml micafungin while greatly reduced growth was seen with 64 µg/ml fluconazole (Fig. 5, bottom panel). This contrasts with, for example, mutant P822L where strong growth was seen with 100 µg fluconazole while there was barely growth with 12 ng/ml micafungin (Fig. 4). As stated above, these results can be explained by the fact that different *PDR1* mutations have different effects on the expression of the targets of this transcription factor (Caudle *et al*., 2011).

**Fig. 5.**
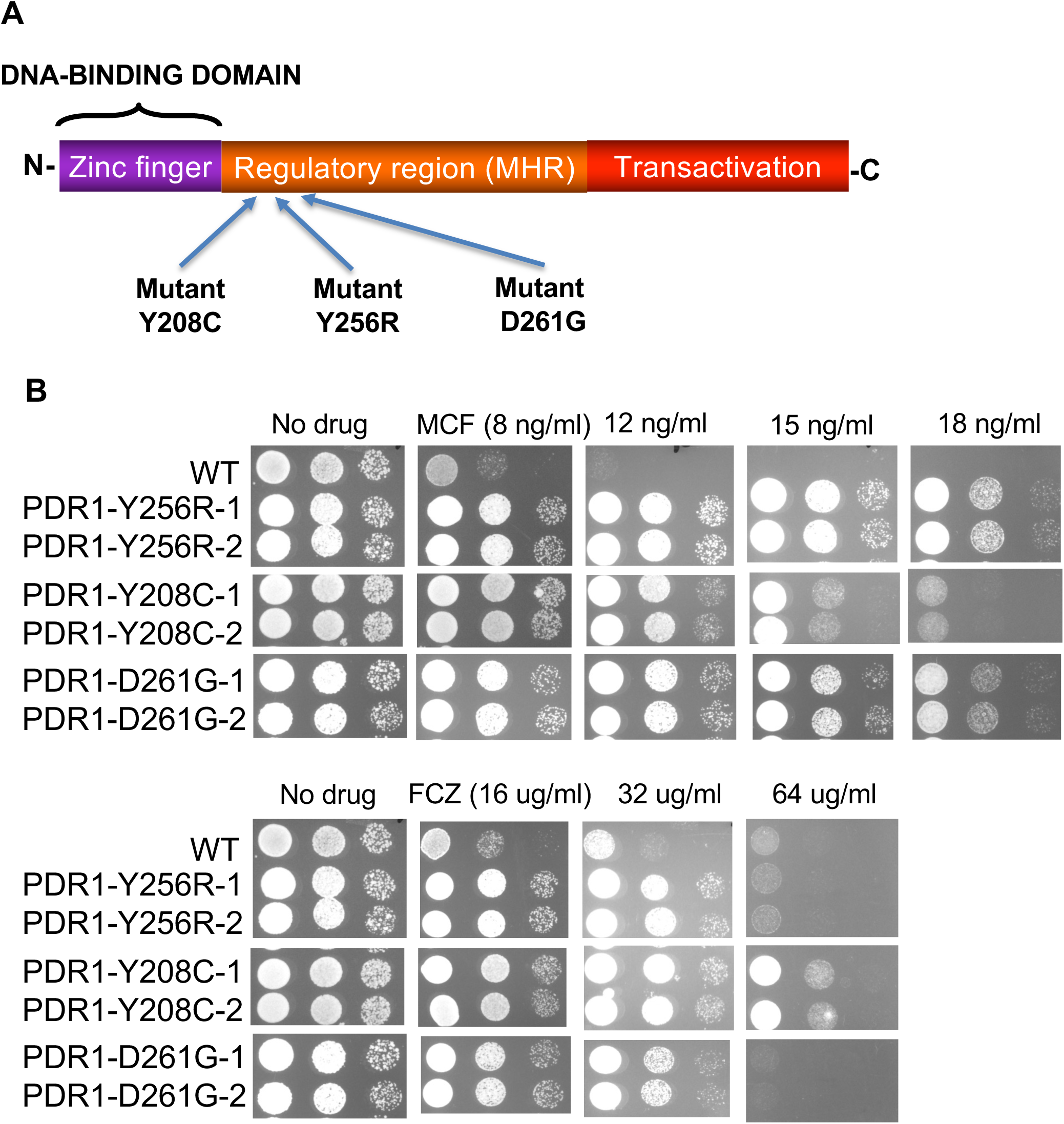
*PDR1* mutants show preferential resistance to micafungin as compared to fluconazole (FCZ). A) Location of the mutations obtained by random mutagenesis. Functional domains of *PDR1* are also indicated. B) Spotting assays were performed with wild-type strain 66032 and mutant strains (with 2 independent integrants ‘-1’ and ‘-2’) CG350, CG351, CG360, CG361, CG354, and CG355.

We were interested to determine if the autoregulatory loop of *PDR1* involved in conferring resistance to azoles is also implicated in micafungin resistance. To this end, we introduced into a Δ*pdr1* strain various expression vectors with mutations in the coding region of *PDR1* and/or the PDREs (Fig. 6A). In the absence of a drug, similar growth was observed for all expression vectors tested (Fig. 6A, left panel). At a high concentration of fluconazole (128 µg /ml), growth was observed with the *PDR1* mutant Y206C but not when PDREs were mutated or with a wild-type strain (Fig. 6A, middle panel). Similarly, mutant Y208C showed resistance to micafungin only with intact PDREs (Fig. 6A, right panel). We tested if the increased resistance to micafungin is due to increased promoter activity of *PDR1*-Y208C. To this end a PDR1-lacZ reporter was introduced into a wild-type strain of a strain where the native *PDR1* gene carries a Y208C mutation (Fig. 6B). Βeta-galactosidase was increased approximately 19-fold in strain *PDR1*-Y208C as compared to a wild-type strain. Moreover, mutating the *PDR1* PDREs drastically decreased Βeta-galactosidase activity in agreement with spotting assays (Fig. 6A). Thus, the *PDR1* auto-regulatory loop is necessary to confer resistance to micafungin via increased promoter activity.

**Fig. 6.**
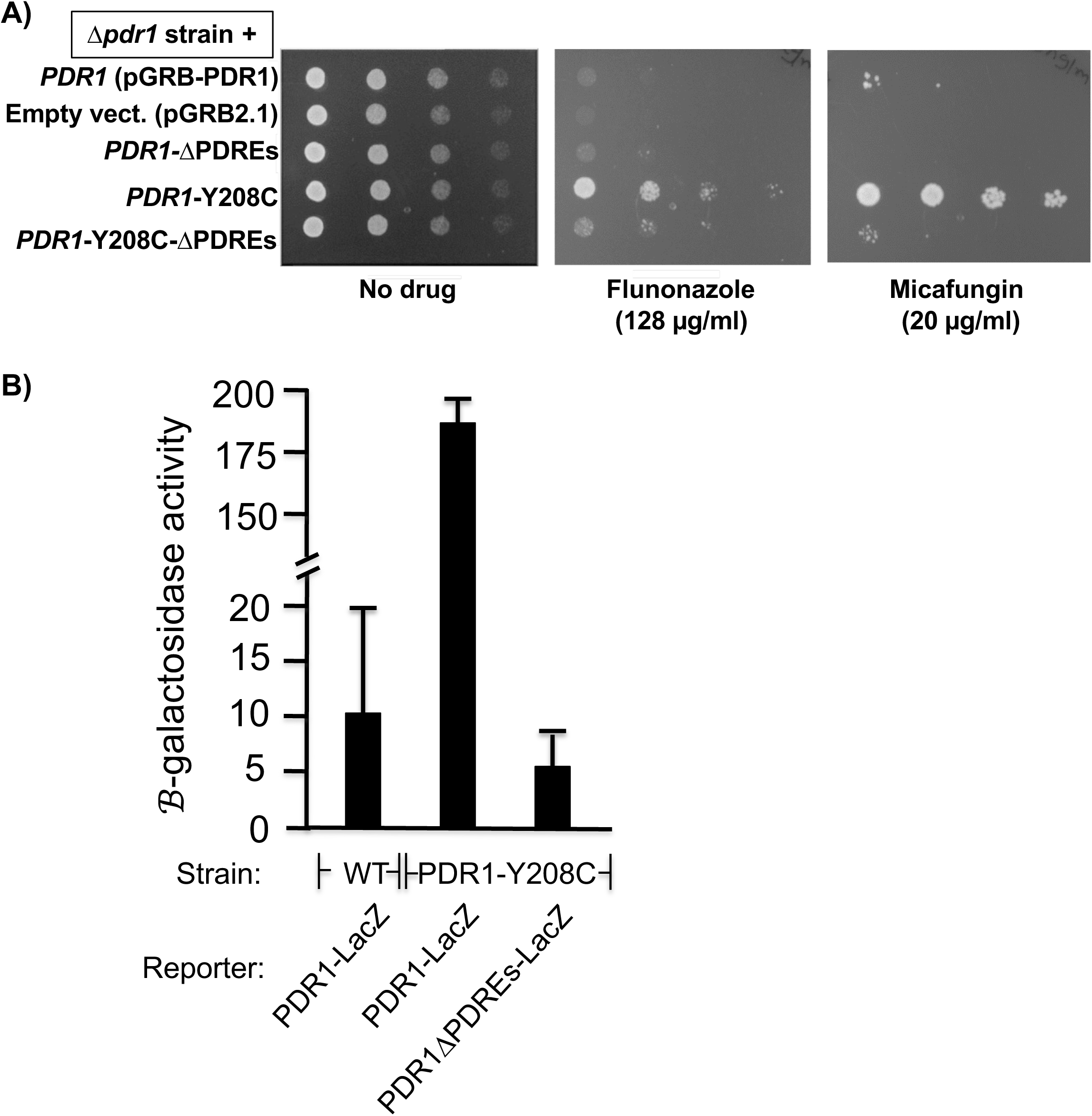
PDREs are required to confer micafungin resistance of mutant *PDR1*-Y208C. A) The Δ*pdr1* strain CG197 was transformed with the plasmids indicated on the left. Colonies were then grown overnight under selective conditions for the plasmids, serially diluted and spotted on plates lacking uracil and containing or not antifungal drugs, as indicated at the bottom of the panels. B) A wild-type strain (‘WT’, 66032) and strain CG260 (‘*PDR1*-Y208C) were transformed with plasmids indicated at the bottom of the panel. Colonies were grown overnight on minimal media lacking uracil. Cells were diluted to an OD_600_ of 0.2 in selective media and grown for 4h. Β-galactosidase assays were performed as described in Material and Methods.

## DISCUSSION

Previous studies have shown that mitochondrial defects in *C. glabrata* results in resistance to azoles, a process mediated by the transcription factor Pdr1 (Vermitsky and Edlind, 2004; Tsai *et al*., 2006). Interestingly, our results show that mitochondrial defects also increase resistance to another class of antifungals, the echinocandins. This process is mediated by Pdr1 since deletion of its gene in a rho^-^ background abolishes resistance to micafungin (Fig. 3A). Moreover, overexpression of *PDR1* by replacing its native promoter by the strong promoter *ADH1* also confers resistance to micafungin (Fig. 3B), as observed for fluconazole (Paul *et al*., 2011). In line with these results, an in vivo study has shown that rho^-^ cells outcompete wild-type cells upon echinocandin treatment (Arastehfar *et al*., 2023).

As stated above, Pdr1 mutants acts by increasing the expression of drug efflux pumps, thereby decreasing the intracellular concentration of azoles and, as a result, conferring resistance to azoles. However, it is not clear how mitochondrial defects result in increased resistance to azoles. This is also true for echinocandins. What is the Pdr1 target(s) responsible for conferring resistance to echinocandins? It is unlikely that Pdr1 acts via increased expression of drug efflux pumps since echinocandins targets Fks1 and Fks2, the catalytic subunits of 1,3-beta-D-glucan synthase, that are located at the plasma membrane. In *C. albicans*, overexpression of drug efflux pumps such as *CDR1* and *CDR2* does not alter susceptibility to echinocandins (Niimi *et al*., 2006). One possibility would be that Pdr1 controls expression of *FKS1* and *FKS2*. However, when performing expression profiling of strains with a rho^-^ strain (or a strain carrying the *PDR1* mutation Y208C) as compared to a wild-type strain treated or not with micafungin, results show no altered expression of *FKS1* or *FKS2* (data not shown). Similar results were obtained in a different study where expression profiling was performed with a rho^-^ strain as compared to a wild-type strain (Ferrari *et al*., 2011).

It is not clear why with the random mutagenesis of *PDR1*, we identified mutations in *PDR1* that confer resistance to micafungin are located in a region located just downstream of the DNA binding domain. This contrasts with azoles where gain-of-function mutations of *PDR1* are located at various regions of the *PDR1* ORF (Ferrari *et al*., 2009). When comparing MIC of rho^-^ strains with wild-type strains, differences in MICs are approximatively 2 to 4-fold (Fig. 2). This is in contrast with gain-of-function of Pdr1 that greatly increase resistance to azoles (e.g. (Ferrari *et al*., 2009)). However, these mutants were generally obtained from resistant clinical isolates that were ‘selected’ *in vivo* while the Pdr1 mutants described in this study were obtained from the analysis of a relatively small number of mutants generated in vitro by random mutagenesis. While resistance to echinocandins has been attributed to mutations in the *FKS1* or the *FKS2* genes (Perlin, 2015), it is possible that strong resistance to this class of antifungal drugs due to mutation in *PDR1* has been overlooked.

## Acknowledgments

We are grateful to Dr. B.P. Cormack for the gift of plasmid pGRB2.1. Plasmid pJH2972 was a gift from James Haber (Addgene plasmid # 100956). We are grateful to Dr. F. Robert (Université de Montréal) for critical reading of the manuscript. We also thank Dr. Marcel Behr for support. This work was supported by grants to BT from the Natural Sciences and Engineering Research Council of Canada, the Fonds de recherche du Québec-Nature et Technologies and the Research Institute of the McGill University Health Centre and to FD from by the Agence Nationale pour la Recherche (CANDIHUB project, grant number ANR-14-CE14-0018-02).

