## Supplementary Figs. S1 for "Mitochondrial defects result in decreased susceptibility to echinocandins via the transcriptional regulator Pdr1 in *Candida glabrata*"

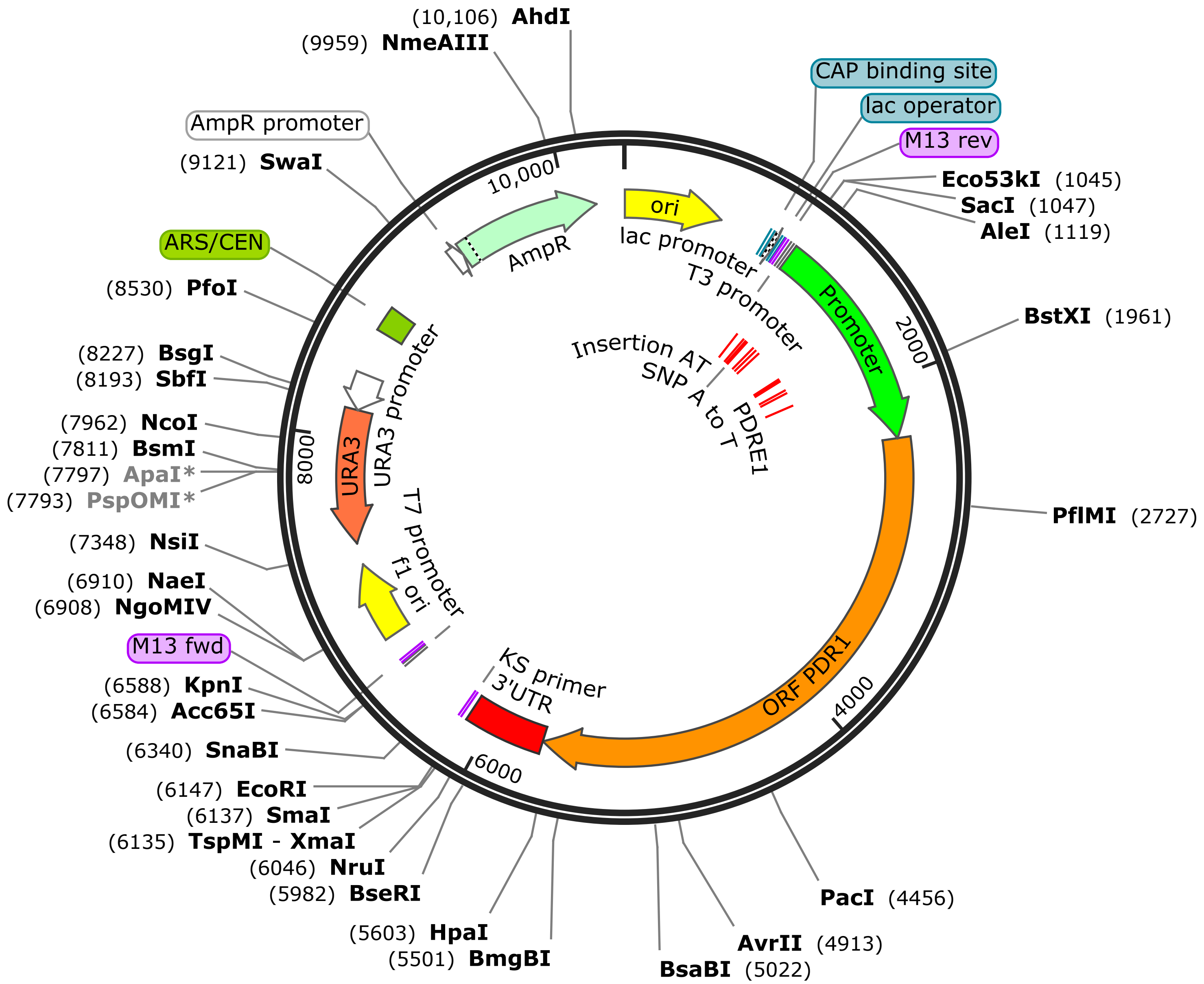


**pGRB-PDR1**
10,349 bp

...

TTGAGATCCTTTTTTTCTGCGCGTAATCTGCTGCTTGCAAACAAA 45
AAAACCACCGCTACCAGCGGTGGTTTGTTTGCCGGATCAAGAGCT 90
ACCAACTCTTTTTCCGAAGGTAACTGGCTTCAGCAGAGCGCAGAT 135
ACCAAATACTGTTCTTCTAGTGTAGCCGTAGTTAGGCCACCACTT 180
CAAGAACTCTGTAGCACCGCCTACATACCTCGCTCTGCTAATCCT 225
GTTACCAGTGGCTGCTGCCAGTGGCGATAAGTCGTGTCTTACCGG 270
GTTGGACTCAAGACGATAGTTACCGGATAAGGCGCAGCGGTCGGG 315
CTGAACGGGGGGTTCGTGCACACAGCCCAGCTTGGAGCGAACGAC 360
CTACACCGAACTGAGATACCTACAGCGTGAGCTATGAGAAAGCGC 405
CACGCTTCCCGAAGGGAGAAAGGCGGACAGGTATCCGGTAAGCGG 450
CAGGGTCGGAACAGGAGAGCGCACGAGGGAGCTTCCAGGGGGAAA 495
CGCCTGGTATCTTTATAGTCCTGTCGGGTTTCGCCACCTCTGACT 540
TGAGCGTCGATTTTTGTGATGCTCGTCAGGGGGGCGGAGCCTATG 585
GAAAAACGCCAGCAACGCGGCCTTTTTACGGTTCCTGGCCTTTTG 630
CTGGCCTTTTGCTCACATGTTCTTTCCTGCGTTATCCCCTGATTC 675
TGTGGATAACCGTATTACCGCCTTTGAGTGAGCTGATACCGCTCG 720
CCGCAGCCGAACGACCGAGCGCAGCGAGTCAGTGAGCGAGGAAGC 765
GGAAGAGCGCCCAATACGCAAACCGCCTCTCCCCGCGCGTTGGCC 810
GATTCATTAATGCAGCTGGCACGACAGGTTTCCCGACTGGAAAGC 855
GGGCAGTGAGCGCAACGCAATTAATGTGAGTTAGCTCACTCATTA 900
GGCACCCCAGGCTTTACACTTTATGCTTCCGGCTCGTATGTTGTG 945
TGGAATTGTGAGCGGATAACAATTTCACACAGGAAACAGCTATGA 990
CCATGATTACGCCAAGCGCGCAATTAACCCTCACTAAAGGGAACA 1035
AAAGCTGGAGCTCGCATTATCTAGGTGGCTGGGTGCCGGTAATCT 1080
GAGTTTGGATATATATATATGCGTTTGATGAAGGCACAATGGTGG 1125
GGGTCCGATTTTAAAGATATTTCCTTTTTAGTTTTGTTGTTAT 1170
TTTTTGATATAAGCCGAATGATCATCTGAATCTAAATTTGAAATG 1215
AAAGACTTCGTTAGTTCTTTTCCTTTTTCAAGTGGGTATCTTTAC 1260
ACTACTGAGATGAAAGCAATAACTGTTGATTGTACCCATACAGAA 1305
GAAAACTTAGAAAAATGAAATACTAGTACATTAAGAAAAAAACAG 1350
CAGAAAAAAAAAAGAAAAGCTGAAGCATTATTATACATCGTAACA 1395
AACATTTCCTCATAGATCTAATCTTTTACTTCAGCAATTTCTAGT 1440
CTCTGCAACAAATTACACTAAAAAAATTACAAATCTTGCACGAGT 1485
ATCATAACGATGCAGTGTCAAGGTCTGAGCATACTGAGGTTATTT 1530
TGTTGGCTTGTATTATCAGTTTCTTTTCTTTCTTCCTTTTTTGTT 1575
TTTTTTTTGTATTTCTTTTGGAGTTGTCCGTTTCCCGCAACCAGA 1620
CAAAGACCCCAGTATTTGTAGACTCATTCCACGGAGCAAATTGTG 1665
CTATATGGATTATTTCTCTGTGCTTCATTTTCTACCTCGTAGATT 1710
AGGTTACGTTCAAATTTTAAGTGGCAAAACCATATCCTGATCGTC 1755
AGTACTAAACCTGAGTTGTCCCGATGAGCCTCCTATTCCGTGGAA 1800
AGATGGCTTTTCCTATTGATCTAAGTGGATAAGCAGGTATTGCGT 1845
TGATCATTATAATTGTGGGTAAAACTAGGTATTAGGTCCTTCTAA 1890
TAGTCATCTTTATTGTTGTAAAAATTGTTTCGATGGCGTATTCAT 1935
AGAATCCGAACATATATACCAACCAAATGGATTTTAAAGGAAAAA 1980
GCAAATACATTAAGGTCTGTTGTAGGAAAAGACCATATTGATTTA 2025
TCTATTTAGTTGAGTAACAGAAATCACATATAGTTTTAAATTTTT 2070
GGTAATCAAAGACACTTGAAACTATAAAACATTTGGTTGAAGAAG 2115
AAGTATTAACTTTGCTTGGAGAAAAGCGTGCCCACACTTGCTTGG 2160
TTTTCAGGATAATAACAAGCATAGAGGCGCTGTTATTACTTAAAC 2205
AATTTTTAAGTAACACATTCAAACTTCCATTACTTCGTACCCCAT 2250
ATCGTATTGCCATTGTGATATGGAATTAGTGTTTTATTCTGCCTT 2295

```
TTTTTTTAGAATATATTGGTAAAGTCATTCTTTAGCTACGTTATT    2340
GAGAGAATATGCAAACATTAGAAACTACATCAAAATCAAATCCAG    2385
GGGAAGTCAAAGCACAGAAGCCTAGTACAAGAAGAACAAAAGTTG    2430
GAAAAGCTTGTGATAGCTGTAGAAGGAGGAAAATAAAATGTAATG    2475
GGCTAAAACCTTGTCCATCTTGTACAATCTATGGCTGTGAATGTA    2520
CATATACTGATGCAAAATCGACAAAAAATCTCAAATCAAATGATG    2565
CAGGTAAATCAAAACCAACAGGGAGAGTATCAAAGAATAAAGAAA    2610
CTACTAGAGTCGACAAAGATATTAGGAAATTAGAGCAGCAGTATG    2655
TCCCTATTAATGCTAATATTCATGTTGGTCCCAGGTTCCCCTCCG    2700
AGAATATATTGAATGGATATCCACAATGTGGAGCACCACAGAACA    2745
ATGTTGTGGGTAATCCACTAGCGGTTAATACTCAATGCCATAGAG    2790
GTCTTTCTGAAACTCCTATGTCCTCAACATTCAAAGAATCTAACT    2835
TAAGAGATGATCGGCTACTACAGTCATCAGATACAGATGATATGA    2880
GGAATGGTGACTCGGAAGAAAGGGACTTGAAAGGGAGTGACAGCG    2925
AGAATGTCAAAAGTAAAGACAATAAAAGTGATCCTTTGATTATAT    2970
ACAAAGATGATACACATATTGAAAGCACGGTTAATAAACTAACAC    3015
AGGCAGTTAATGAACTCAAATCACTTCAAAATGCACCCAGTTCGA    3060
TAAAATCATCCATTGACGCCATTGAGTTACAACTTAGAAACATTT    3105
TAGACAACTGGAAACCAGAGGTAGATTTCGAGAAAGCAAAGATTA    3150
ATGAAAGTGCCACCACTAAGTCACTTGAAACAAACTTGCTGAGGA    3195
ATAAATACACTAATCACGTTCATTTAACAAGATTTAGGATATGGA    3240
TAGATTATAAAAATGCGAACAAAAACAATCATTTTATGGGAGAGT    3285
GTGGATTTAGTCTTGCAGAATCTTTTTTTGCTTCTAATCAGCCAT    3330
TGGTCGATGAATTGTTTGGGTTGTATTCCCAGGTAGAGGCCTTTT    3375
CTTTGCAAGGTCTTGGTTACTGTGTTCACCTTTATGAGCCATATA    3420
TGAAAACTGAGGAAGCGATAAAACTGATGAAAGAGACCTTATATA    3465
TTATACTACGGTTTATTGATATATGTGTTCACCATATCAATGAAG    3510
AGTCGATATCGATTGCCAACCCGTTAGAAACATATTTACGAAAAA    3555
AACATCTAATGCCTATGACTCCTACACCAAGGTCGTCCTATGGAA    3600
GTCCACAAAGTGCTAGTACAAAGAGCTTGGTAAGTAAGATAATAG    3645
AGAGAATACCGCAACCGTTTATTGAGAGTGTAACTAATGTGTCGA    3690
GTCTTCAACTATTAGATCTTCGAGATGACGAGTCAAAAATGTTTG    3735
GAACATTGCTGAACATGTGTAAGTCTATAAGGCGAAAATTTGACT    3780
CTGTTATGAGCGATTACGATTCCATTGTCACAGAAAAATCCGAAG    3825
GCGAACAAAATGATGGTAAAGTAACTGTAGCTGAGTTCACATCTT    3870
TGTGTGAAGCGGAAGAAATGCTCTTAGCATTATGCTATAACTATT    3915
ATAATCTGACGTTATACAGTTTCTTTGAATTTGGGACTAATATTG    3960
AATACATGGAACATCTGTTGCTTCTTCTTGAAGAACAGCTTGCTC    4005
TCGACGAATACTATGGTTTTGAAAAGGTCTTGAATGTAGCTGTTG    4050
CAAATGCTAAAAAAATGGGTTTCCACCGTTGGGAGTTTTACGTCG    4095
GCTATGAAGAGTCGACTGCTGAAAAGAGGCGGCTACTATGGTGGA    4140
AGTTATACAATTATGAAAAAGCCAGTACTATGAAGAAGGGTTTTT    4185
TTTCTGTGATTGATGATGCTACTGTCAACTGTTTATTACCTAAGA    4230
TTTTTAGAAACTTTGGCTATCTGGATAGGGTGGAGTTTCTAGAAA    4275
ATATTCAAAAGCCAATGGATCTTAGTGTGTTTTCCGATGTTCCAA    4320
TTTCTGTCCTTTGTAAATACGGTGAGTTGGCCCTTACAATAGTTA    4365
CCAGTGAGTTTCATGAAAAATTTTTATATGCTGATAGATACACTT    4410
CTATTCGAAATTCCGCGAAACCGCCGACATTAAAAAACCAATTAA    4455
TTAAGGAAATTGTGGATGGTATAGCTTATACAGAGACATCATATG    4500
AGGCAATCAGAAAGCAAACTGCAAAACTATGGGATATTGCATTAG    4545
GTAAGGTGACCAAAGATAAAATCAATAAAGAAGATACAGCAGCAG    4590
```

CTAGCAAATTTACTTTGAGTTATGAATATCACAGATTCAGGCTAA 4635
TCAATATGGCAGACAATTTAATTGCTAGACTAATGGTGAAACCAA 4680
AATCAGATTGGCTAATATCAGTCATGAAGGGGCATCTTAACAGAC 4725
TATATGAGCACTGGAAAGTAATGAATGAAATTATCCTAAGTATGG 4770
ACAACGATTATTCAATTGCAACAACGTTCGAATATTATGCACCAT 4815
CATGTCTGTGTTTAGCTACGCAGACTTTCCTTATTGTGAGGAATA 4860
TGGAAATGGATGATGTCAAGATGATGGTTGCAGTATATAAAAGAT 4905
TTCTTAACCTAGGAATGTTTTTGCAGAGTGCCAAAGTATGCAGCC 4950
TTGCCGATAGTCATACATTCAGAGATTTTTCTAGATCTTTTTCCT 4995
TTATTACGATAATTTCAAGATTGATGATAATCGAATTTATGCAAA 5040
TTAAAGAATTGACGAAGGTAGAGTTTATTGAGAAGTTTTCTGAAG 5085
TATGCCCTGACCTTGCAGATCTACCTCCGATGCTTCTAGATCCAA 5130
ACTCTTGCTTATATTTTTCATTGTTACAGCAGATTAAGAAATCTG 5175
GTTTTACGTTGTCATTCAAAAAAATTCTTGAAGACGCTAGAATGA 5220
TGGACTTCAATTACGACCGCAATTTGGACTCAGAGGCCATTAAAA 5265
AGTGCAATGGTGAATTTAGCAAGTCAATGCCTTCCTGTACCAATG 5310
TCTCAGATACCACCACCGCTGTTTCTGACAATAGTGCTAAGAAGA 5355
AAGCTTCAATGGGGTCGGCGAGGGTAAATTCAACTGATACACTAA 5400
CTGCATCTCCCTTATCGGGCTTAAGGAATCAAACGCAGTTGGATT 5445
CTAAAGACAGTGTTCCATCTCTCGAGGCTTATACACCAATTGATT 5490
CTGTCTCTGACGTGCCCACTGGGGAGATCAACGTTCCATTCCCTC 5535
CTGTTTATAATCAAAATGGATTGGATCAGCAAACCACTTATAATT 5580
TGGGAACTTTAGATGAGTTTGTTAACAAGGGAGATTTGAATGAAC 5625
TCTATAATAGCCTATGGGGTGACCTATTTTCTGATGTTTACTTGT 5670
GATGCTATTAATTTAGACAGTGTGCATAGCCTGTATAATTTTATA 5715
TTTAATCTGAAAAATCATTTTTTAGTGGTTATGTTGCATAATTTT 5760
ACATGAGACTTAGACTACCTCAAATTAGCGATAACACTACAATAT 5805
CTCTCATTAAAAAATTGTAGTTTAATATATGATGATTATTAAAAC 5850
GTATTGTACAAATTATAACTTATAAAAGAAAAACCCTTCGATTCT 5895
GAATGAAGATTCTAACAGCAAAACTGTTTGGGTTTATTCAATATT 5940
AAGGATACCCATTATCCTTTTTCCGTGAGGAGTGTTTCTAATATT 5985
ACCAACTAGTAGTGGTGTAACCAAGTCAACCAATTTCTTATACAT 6030
ATAAGGATTTGTGTCGCGACTAATATCCAATGCAGTCTGTAGCAC 6075
ATAGTTACCGAAGCTATCGTTTAACAGAGCTTGAACCTCAGCACT 6120
ACCTTCGTTGATCCCCCGGGCTGCAGGAATTCGATATCAAGCTTA 6165
TCGATACCGTCGACCTCGAGAACACAACCCACAGCTACCACCATC 6210
AACAATATTTATATATATAACGTACACATAGAAATCACACAAACA 6255
GAGTATTTATTCTTAACTACATGAACTACCATCAGACCGTCTGGG 6300
CCACTATATAATGTGCCATTCATAAACGTGATCACTTTACGTAGC 6345
AGGCAACCCCAGGTGAAAATTTTTCAGCGAGCTGCCAGATTGTCA 6390
GGTGAAAAACTGAAAAAAACTTCTGGGCGATGAGCTTGTGGCGGG 6435
AAATTAAGTATATAAAGCAGTTAGTTTCTTCTGCTTCTTGTGGTT 6480
CTGGGTATTCTTGTAATATTTGGAAGAATAGAAAATCAAGACTAC 6525
TGCCTTTCTTTTCATATTATAGAGGAAGAAATACGCACGAACACG 6570
ATATAGAGGTAAAGGTACCCAATTCGCCCTATAGTGAGTCGTATT 6615
ACGCGCGCTCACTGGCCGTCGTTTTACAACGTCGTGACTGGGAAA 6660
ACCCTGGCGTTACCCAACTTAATCGCCTTGCAGCACATCCCCCTT 6705
TCGCCAGCTGGCGTAATAGCGAAGAGGCCCGCACCGATCGCCCTT 6750
CCCAACAGTTGCGCAGCCTGAATGGCGAATGGACGCGCCCTGTAG 6795
CGGCGCATTAAGCGCGGCGGGTGTGGTGGTTACGCGCAGCGTGAC 6840
CGCTACACTTGCCAGCGCCCTAGCGCCCGCTCCTTTCGCTTTCTT 6885

```
CCCTTCCTTTCTCGCCACGTTCGCCGGCTTTCCCCGTCAAGCTCT 6930
AAATCGGGGGCTCCCTTTAGGGTTCCGATTTAGTGCTTTACGGCA 6975
CCTCGACCCCAAAAAACTTGATTAGGGTGATGGTTCACGTAGTGG 7020
GCCATCGCCCTGATAGACGGTTTTTCGCCCTTTGACGTTGGAGTC 7065
CACGTTCTTTAATAGTGGACTCTTGTTCCAAACTGGAACAACACT 7110
CAACCCTATCTCGGTCTATTCTTTTGATTTATAAGGGATTTTGCC 7155
GATTTCGGCCTATTGGTTAAAAAATGAGCTGATTTAACAAAAATT 7200
TAACGCGAATTTTAACAAAATATTAACGCTTACAATTTCCTGATG 7245
CGGTATTTTCTCCTTACGCATCTGTGCGGTATTTCACACCGCATA 7290
GGGTAATAACTGATATAATTAAATTGAAGCTCTAATTTGTGAGTT 7335
TAGTATACATGCATTTACTTATAATACAGTTTTTTAGTTTTGCTG 7380
GCCGCATCTTCTCAAATATGCTTCCCAGCCTGCTTTTCTGTAACG 7425
TTCACCCTCTACCTTAGCATCCCTTCCCTTTGCAAATAGTCCTCT 7470
TCCAACAATAATAATGTCAGATCCTGTAGAGACCACATCATCCAC 7515
GGTTCTATACTGTTGACCCAATGCGTCTCCCTTGTCATCTAAACC 7560
CACACCGGGTGTCATAATCAACCAATCGTAACCTTCATCTCTTCC 7605
ACCCATGTCTCTTTGAGCAATAAAGCCGATAACAAAATCTTTGTC 7650
GCTCTTCGCAATGTCAACAGTACCCTTAGTATATTCTCCAGTAGA 7695
TAGGGAGCCCTTGCATGACAATTCTGCTAACATCAAAAGGCCTCT 7740
AGGTTCCTTTGTTACTTCTTCTGCCGCCTGCTTCAAACCGCTAAC 7785
AATACCTGGGCCCACCACACCGTGTGCATTCGTAATGTCTGCCCA 7830
TTCTGCTATTCTGTATACACCCGCAGAGTACTGCAATTTGACTGT 7875
ATTACCAATGTCAGCAAATTTTCTGTCTTCGAAGAGTAAAAAATT 7920
GTACTTGGCGGATAATGCCTTTAGCGGCTTAACTGTGCCCTCCAT 7965
GGAAAAATCAGTCAAGATATCCACATGTGTTTTTAGTAAACAAAT 8010
TTTGGGACCTAATGCTTCAACTAACTCCAGTAATTCCTTGGTGGT 8055
ACGAACATCCAATGAAGCACACAAGTTTGTTTGCTTTTCGTGCAT 8100
GATATTAAATAGCTTGGCAGCAACAGGACTAGGATGAGTAGCAGC 8145
ACGTTCCTTATATGTAGCTTTCGACATGATTTATCTTCGTTTCCT 8190
GCAGGTTTTTGTTCTGTGCAGTTGGGTTAAGAATACTGGGCAATT 8235
TCATGTTTCTTCAACACTACATATGCGTATATATACCAATCTAAG 8280
TCTGTGCTCCTTCCTTCGTTCTTCCTTCTGTTCGGAGATTACCGA 8325
ATCAAAAAAATTTCAAGGAAACCGAAATCAAAAAAAAGAATAAAA 8370
AAAAAATGATGAATTGAAAAGGTGGTATGGTGCACTCTCAGTACA 8415
ATCTGCTCTGATGCCGCATAGTTAAGCCAGCCCCGACACCCGCCA 8460
ACACCCGCTGACGCGCCCTGACGGGCTTGTCTGCTCCCGGCATCC 8505
GCTTACAGACAAGCTGTGACCGTCTCCGGGAGCTGCATGTGTCAG 8550
AGGTTTTCACCGTCATCACCGAAACGCGCGAGACGAAAGGGCCTC 8595
GTGATACGCCTATTTTTATAGGTTAATGTCATGATAATAATGGTT 8640
TCTTAGCAGGTTAGACTAACATGCAAAAAGTTATATTCATTTTT 8685
GTAGTTATTCTTGTTTTCATTAGGATGATTACACACTGTGGATGT 8730
GGATTAATCGATCATTGTTAATTTAGTGTAGCTACTACCATTTCA 8775
AACAAAACGTAAATTCTTGCAGTCGTACTGGATCTGTGAATCTAT 8820
TAGTATATATGAATTAAAGTAGCTTGTACATATTATTCTGTTGAA 8865
TCATATCGCAGAGCATTGGTTGAAATCCAAAATATAAAAATGTAA 8910
TATCACAAAAAATAATACTAATTCTAACATTAATGGGTCAGATTT 8955
TTAGTGAATACTTAAATTTATAATATCGTCTATTTAAGCTAGCAA 9000
ATGGAACAACATTTAAAGTAAGAACATCATATCTACATGAAAATG 9045
TATATTTCAATCTGACTAATAACGCAGAGCACATCTTTCAGTGAT 9090
GTCTGTCACATGATCAAAAAGAATTGTATTTAAATAATTCATAAT 9135
AAAAGCTTAAAAAATTACAATAATGAAAATAAAGTAAATAATGAC 9180
```

ATGGGTAAGAGTTCCGAAATAGAATCTTAGTGTACAAAACAAAAA 9225
TTGCATCATTAGAGATCCCCACATTCAAATATGTATCCGCTCATG 9270
AGACAATAACCCTGATAAATGCTTCAATAATATTGAAAAAGGAAG 9315
AGTATGAGTATTCAACATTTCCGTGTCGCCCTTATTCCCTTTTTT 9360
GCGGCATTTTGCCTTCCTGTTTTTGCTCACCCAGAAACGCTGGTG 9405
AAAGTAAAAGATGCTGAAGATCAGTTGGGTGCACGAGTGGGTTAC 9450
ATCGAACTGGATCTCAACAGCGGTAAGATCCTTGAGAGTTTTCGC 9495
CCCGAAGAACGTTTTCCAATGATGAGCACTTTTAAAGTTCTGCTA 9540
TGTGGCGCGGTATTATCCCGTATTGACGCCGGGCAAGAGCAACTC 9585
GGTCGCCGCATACACTATTCTCAGAATGACTTGGTTGAGTACTCA 9630
CCAGTCACAGAAAAGCATCTTACGGATGGCATGACAGTAAGAGAA 9675
TTATGCAGTGCTGCCATAACCATGAGTGATAACACTGCGGCCAAC 9720
TTACTTCTGACAACGATCGGAGGACCGAAGGAGCTAACCGCTTTT 9765
TTGCACAACATGGGGGATCATGTAACTCGCCTTGATCGTTGGGAA 9810
CCGGAGCTGAATGAAGCCATACCAAACGACGAGCGTGACACCACG 9855
ATGCCTGTAGCAATGGCAACAACGTTGCGCAAACTATTAACTGGC 9900
GAACTACTTACTCTAGCTTCCCGGCAACAATTAATAGACTGGATG 9945
GAGGCGGATAAAGTTGCAGGACCACTTCTGCGCTCGGCCCTTCCG 9990
GCTGGCTGGTTTATTGCTGATAAATCTGGAGCCGGTGAGCGTGGG 10,035
TCTCGCGGTATCATTGCAGCACTGGGGCCAGATGGTAAGCCCTCC 10,080
CGTATCGTAGTTATCTACACGACGGGGAGTCAGGCAACTATGGAT 10,125
GAACGAAATAGACAGATCGCTGAGATAGGTGCCTCACTGATTAAG 10,170
CATTGGTAACTGTCAGACCAAGTTTACTCATATATACTTTAGATT 10,215
GATTTAAAACTTCATTTTTAATTTAAAAGGATCTAGGTGAAGATC 10,260
CTTTTTGATAATCTCATGACCAAAATCCCTTAACGTGAGTTTTCG 10,305
TTCCACTGAGCGTCAGACCCCGTAGAAAAGATCAAAGGATCTTC ••• 10,349

Sequence: pPDR1-C.dna (Circular / 9892 bp)
Enzymes: Unique 6+ Cutters (36 of 653 total)
Features: 30 total

Unique Cutters **Bold**

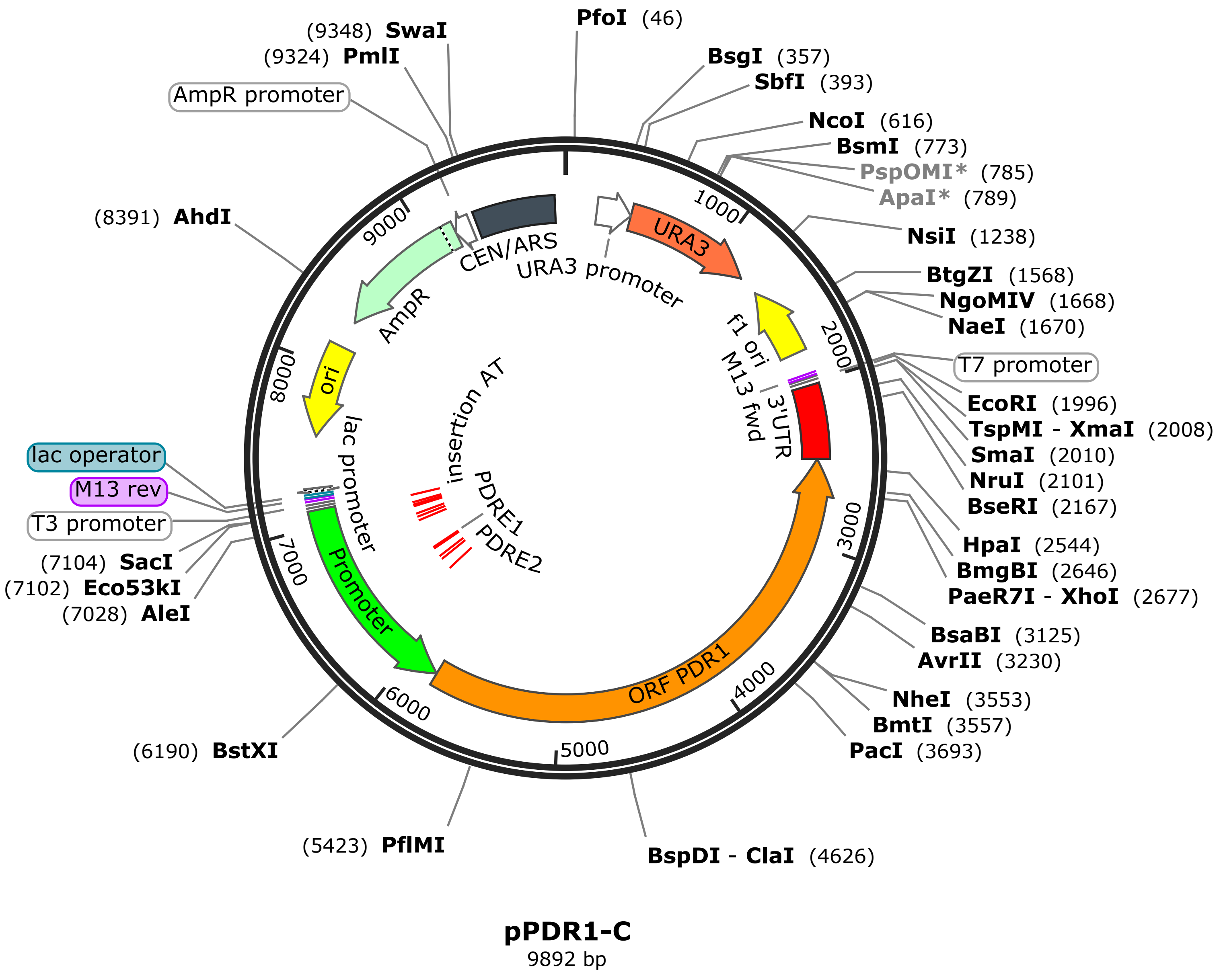

```
TCGCGCGTTTCGGTGATGACGGTGAAAACCTCTGACACATGCAGCTCCCG    50
GAGACGGTCACAGCTTGTCTGTAAGCGGATGCCGGGAGCAGACAAGCCCG   100
TCAGGGCGCGTCAGCGGGTGTTGGCGGGTGTCGGGGCTGGCTTAACTATG   150
CGGCATCAGAGCAGATTGTACTGAGAGTGCACCACGCTTTTCAATTCAAT   200
TCATCATTTTTTTTTTATTCTTTTTTTTGATTTCGGTTTCTTTGAAATTT   250
TTTTGATTCGGTAATCTCCGAACAGAAGGAAGAACGAAGGAAGGAGCACA   300
GACTTAGATTGGTATATATACGCATATGTAGTGTTGAAGAAACATGAAAT   350
TGCCCAGTATTCTTAACCCAACTGCACAGAACAAAAACCTGCAGGAAACG   400
AAGATAAATCATGTCGAAAGCTACATATAAGGAACGTGCTGCTACTCATC   450
CTAGTCCTGTTGCTGCCAAGCTATTTAATATCATGCACGAAAAGCAAACA   500
AACTTGTGTGCTTCATTGGATGTTCGTACCACCAAGGAATTACTGGAGTT   550
AGTTGAAGCATTAGGTCCCAAAATTTGTTTACTAAAAACACATGTGGATA   600
TCTTGACTGATTTTTCCATGGAGGGCACAGTTAAGCCGCTAAAGGCATTA   650
TCCGCCAAGTACAATTTTTTACTCTTCGAAGACAGAAAATTTGCTGACAT   700
TGGTAATACAGTCAAATTGCAGTACTCTGCGGGTGTATACAGAATAGCAG   750
AATGGGCAGACATTACGAATGCACACGGTGTGGTGGGCCCAGGTATTGTT   800
AGCGGTTTGAAGCAGGCGGCAGAAGAAGTAACAAAGGAACCTAGAGGCCT   850
TTTGATGTTAGCAGAATTGTCATGCAAGGGCTCCCTATCTACTGGAGAAT   900
ATACTAAGGGTACTGTTGACATTGCGAAGAGCGACAAAGATTTTGTTATC   950
GGCTTTATTGCTCAAAGAGACATGGGTGGAAGAGATGAAGGTTACGATTG  1000
GTTGATTATGACACCCGGTGTGGGTTTAGATGACAAGGGAGACGCATTGG  1050
GTCAACAGTATAGAACCGTGGATGATGTGGTCTCTACAGGATCTGACATT  1100
ATTATTGTTGGAAGAGGACTATTTGCAAAGGGAAGGGATGCTAAGGTAGA  1150
GGGTGAACGTTACAGAAAAGCAGGCTGGGAAGCATATTTGAGAAGATGCG  1200
GCCAGCAAAACTAAAAAACTGTATTATAAGTAAATGCATGTATACTAAAC  1250
TCACAAATTAGAGCTTCAATTTAATTATATCAGTTATTACCCTGCGGTGT  1300
GAAATACCGCACAGATGCGTAAGGAGAAAATACCGCATCAGGAAATTGTA  1350
AACGTTAATATTTTGTTAAAATTCGCGTTAAATTTTTGTTAAATCAGCTC  1400
ATTTTTTAACCAATAGGCCGAAATCGGCAAAATCCCTTATAAATCAAAAG  1450
AATAGACCGAGATAGGGTTGAGTGTTGTTCCAGTTTGGAACAAGAGTCCA  1500
CTATTAAAGAACGTGGACTCCAACGTCAAAGGGCGAAAAACCGTCTATCA  1550
GGGCGATGGCCCACTACGTGAACCATCACCCTAATCAAGTTTTTTGGGGT  1600
CGAGGTGCCGTAAAGCACTAAATCGGAACCCTAAAGGGAGCCCCCGATTT  1650
AGAGCTTGACGGGGAAAGCCGGCGAACGTGGCGAGAAAGGAAGGGAAGAA  1700
AGCGAAAGGAGCGGGCGCTAGGGCGCTGGCAAGTGTAGCGGTCACGCTGC  1750
GCGTAACCACCACACCCGCCGCGCTTAATGCGCCGCTACAGGGCGCGTCG  1800
CGCCATTCGCCATTCAGGCTGCGCAACTGTTGGGAAGGGCGATCGGTGCG  1850
GGCCTCTTCGCTATTACGCCAGCTGGCGAAGGGGGATGTGCTGCAAGGC   1900
GATTAAGTTGGGTAACGCCAGGGTTTTCCCAGTCACGACGTTGTAAAACG  1950
ACGGCCAGTgagcgcgctaatacgactcactatagggcgaattggaatt  2000
cCTGCAGCCCGGGGGATCAACGAAGGTAGTGCTGAGGTTCAAGCTCTGTT  2050
AAACGATAGCTTCGGTAACTATGTGCTACAGACTGCATTGGATATTAGTC  2100
GCGACACAAATCCTTATATGTATAAGAAATTGGTTGACTTGGTTACACCA  2150
CTACTAGTTGGTAATATTAGAAACACTCCTCACGGAAAAAGGATAATGGG  2200
TATCCTTAATATTGAATAAACCCAAACAGTTTTGCTGTTAGAATCTTCAT  2250
TCAGAATCGAAGGGTTTTTCTTTTATAAGTTATAATTTGTACAATACGTT  2300
TTAATAATCATCATATATTAAACTACAATTTTTTAATGAGAGATATTGTA  2350
GTGTTATCGCTAATTTGAGGTAGTCTAAGTCTCATGTAAAATTATGCAAC  2400
ATAACCACTAAAAAATGATTTTTCAGATTAAATATAAAATTATACAGGCT  2450
ATGCACACTGTCTAAATTAATAGCATCACAAGTAAACATCAGAAAATAGG  2500
TCACCCCATAGGCTATTATAGAGTTCATTCAAATCTCCCTTGTTAACAAA  2550
```

CTCATCTAAAGTTCCCAAATTATAAGTGGTTTGCTGATCCAATCCATTTT 2600
GATTATAAACAGGAGGGAATGGAACGTTGATCTCCCCAGTGGGCACGTCA 2650
GAGACAGAATCAATTGGTGTATAAGCCTCGAGAGATGGAACACTGTCTTT 2700
AGAATCCAACTGCGTTTGATTCCTTAAGCCCGATAAGGGAGATGCAGTTA 2750
GTGTATCAGTTGAATTTACCCTCGCCGACCCCATTGAAGCTTTCTTCTTA 2800
GCACTATTGTCAGAAACAGCGGTGGTGGTATCTGAGACATTGGTACAGGA 2850
AGGCATTGACTTGCTAAATTCACCATTGCACTTTTTAATGGCCTCTGAGT 2900
CCAAATTGCGGTCGTAATTGAAGTCCATCATTCTAGCGTCTTCAAGAATT 2950
TTTTTGAATGACAACGTAAAACCAGATTTCTTAATCTGCTGTAACAATGA 3000
AAAATATAAGCAAGAGTTTGGATCTAGAAGCATCGGAGGTAGATCTGCAA 3050
GGTCAGGGCATACTTCAGAAAACTTCTCAATAAACTCTACCTTCGTCAAT 3100
TCTTTAATTTGCATAAATTCGATTATCATCAATCTTGAAATTATCGTAAT 3150
AAAGGAAAAAGATCTAGAAAAATCTCTGAATGTATGACTATCGGCAAGGC 3200
TGCATACTTTGGCACTCTGCAAAAACATTCCTAGGTTAAGAAATCTTTTA 3250
TATACTGCAACCATCATCTTGACATCATCCATTTCCATATTCCTCACAAT 3300
AAGGAAAGTCTGCGTAGCTAAACACAGACATGATGGTGCATAATATTCGA 3350
ACGTTGTTGCAATTGAATAATCGTTGTCCATACTTAGGATAATTTCATTC 3400
ATTACTTTCCAGTGCTCATATAGTCTGTTAAGATGCCCCTTCATGACTGA 3450
TATTAGCCAATCTGATTTTGGTTTCACCATTAGTCTAGCAATTAAATTGT 3500
CTGCCATATTGATTAGCCTGAATCTGTGATATTCATAACTCAAAGTAAAT 3550
TTGCTAGCTGCTGCTGTATCTTCTTTATTGATTTTATCTTTGGTCACCTT 3600
ACCTAATGCAATATCCCATAGTTTTGCAGTTTGCTTTCTGATTGCCTCAT 3650
ATGATGTCTCTGTATAAGCTATACCATCCACAATTTCCTTAATTAATTGG 3700
TTTTTTAATGTCGGCGGTTTCGCGGAATTTCGAATAGAAGTGTATCTATC 3750
AGCATATAAAAATTTTTCATGAAACTCACTGGTAACTATTGTAAGGGCCA 3800
ACTCACCGTATTTACAAAGGACAGAAATTGGAACATCGGAAAACACACTA 3850
AGATCCATTGGCTTTTGAATATTTTCTAGAAACTCCACCCTATCCAGATA 3900
GCCAAAGTTTCTAAAAATCTTAGGTAATAAACAGTTGACAGTAGCATCAT 3950
CAATCACAGAAAAAAAACCCTTCTTCATAGTACTGGCTTTTTCATAATTG 4000
TATAACTTCCACCATAGTAGCCGCCTCTTTTCAGCAGTCGACTCTTCATA 4050
GCCGACGTAAAACTCCCAACGGTGGAAACCCATTTTTTTAGCATTTGCAA 4100
CAGCTACATTCAAGACCTTTTCAAAACCATAGTATTCGTCGAGAGCAAGC 4150
TGTTCTTCAAGAAGAAGCAACAGATGTTCCATGTATTCAATATTAGTCCC 4200
AAATTCAAAGAAACTGTATAACGTCAGATTATAATAGTTATAGCATAATG 4250
CTAAGAGCATTTCTTCCGCTTCACACAAAGATGTGAACTCAGCTACAGTT 4300
ACTTTACCATCATTTTGTTCGCCTTCGGATTTTTCTGTGACAATGGAATC 4350
GTAATCGCTCATAACAGAGTCAAATTTTCGCCTTATAGACTTACACATGT 4400
TCAGCAATGTTCCAAACATTTTTGACTCGTCATCTCGAAGATCTAATAGT 4450
TGAAGACTCGACACATTAGTTACACTCTCAATAAACGGTTGCGGTATTCT 4500
CTCTATTATCTTACTTACCAAGCTCTTTGTACTAGCACTTTGTGGACTTC 4550
CATAGGACGACCTTGGTGTAGGAGTCATAGGCATTAGATGTTTTTTTCGT 4600
AAATATGTTTCTAACGGGTTGGCAATCGATATCGACTCTTCATTGATATG 4650
GTGAACACATATATCAATAAACCGTAGTATAATATATAAGGTCTCTTTCA 4700
TCAGTTTTATCGCTTCCTCAGTTTTCATATATGGCTCATAAAGGTGAACA 4750
CAGTAACCAAGACCTTGCAAAGAAAAGGCCTCTACCTGGGAATACAACCC 4800
AAACAATTCATCGACCAATGGCTGATTAGAAGCAAAAAAAGATTCTGCAA 4850
GACTAAATCCACACTCTCCCATAAAATGATTGTTTTTGTTCGCATTTTTA 4900
TAATCTATCCATATCCTAAATCTTGTTAAATGAACGTGATTAGTGTATTT 4950
ATTCCTCAGCAAGTTTGTTTCAAGTGACTTAGTGGTGGCACTTTCATTAA 5000
TCTTTGCTTTCTCGAAATCTACCTCTGGTTTCCAGTTGTCTAAAATGTTT 5050
CTAAGTTGTAACTCAATGGCGTCAATGGATGATTTTATCGAACTGGGTGC 5100

ATTTTGAAGTGATTTGAGTTCATTAACTGCCTGTGTTAGTTTATTAACCG 5150
TGCTTTCAATATGTGTATCATCTTTGTATATAATCAAAGGATCACTTTTA 5200
TTGTCTTTACTTTTGACATTCTCGCTGTCACTCCCTTTCAAGTCCCTTTC 5250
TTCCGAGTCACCATTCCTCATATCATCTGTATCTGATGACTGTAGTAGCC 5300
GATCATCTCTTAAGTTAGATTCTTTGAATGTTGAGGACATAGGAGTTTCA 5350
GAAAGACCTCTATGGCATTGAGTATTAACCGCTAGTGGATTACCCACAAC 5400
ATTGTTCTGTGGTGCTCCACATTGTGGATATCCATTCAATATATTCTCGG 5450
AGGGGAACCTGGGACCAACATGAATATTAGCATTAATAGGGACATACTGC 5500
TGCTCTAATTTCCTAATATCTTTGTCGACTCTAGTAGTTTCTTTATTCTT 5550
TGATACTCTCCCTGTTGGTTTTGATTTACCTGCATCATTTGATTTGAGAT 5600
TTTTTGTCGATTTTGCATCAGTATATGTACATTCACAGCCATAGATTGTA 5650
CAAGATGGACAAGGTTTTAGCCCATTACATTTTATTTTCCTCCTTCTACA 5700
GCTATCACAAGCTTTTCCAACTTTTGTTCTTCTTGTACTAGGCTTCTGTG 5750
CTTTGACTTCCCCTGGATTTGATTTTGATGTAGTTTCTAATGTTTGCATA 5800
TTCTCTCAATAACGTAGCTAAAGAATGACTTTACCAATATATTCTAAAAA 5850
AAAAGGCAGAATAAAACACTAATTCCATATCACAATGGCAATACGATATG 5900
GGGTACGAAGTAATGGAAGTTTGAATGTGTTACTTAAAAATTGTTTAAGT 5950
AATAACAGCGCCTCTATGCTTGTTATTATCCTGAAAACCAAGCAAGTGTG 6000
GGCACGCTTTTCTCCAAGCAAAGTTAATACTTCTTCTTCAACCAAATGTT 6050
TTATAGTTTCAAGTGTCTTTGATTACCAAAAATTTAAAACTATATGTGAT 6100
TTCTGTTACTCAACTAAATAGATAAATCAATATGGTCTTTTCCTACAACA 6150
GACCTTAATGTATTTGCTTTTTCCTTTAAAATCCATTTGGTTGGTATATA 6200
TGTTCGGATTCTATGAATACGCCATCGAAACAATTTTTACAACAATAAAG 6250
ATGACTATTAGAAGGACCTAATACCTAGTTTTACCCACAATTATAATGAT 6300
CAACGCAATACCTGCTTATCCACTTAGATCAATAGGAAAAGCCATCTTTC 6350
CACGGAATAGGAGGCTCATCGGGACAACTCAGGTTTAGTACTGACGATCA 6400
GGATATGGTTTTGCCACTTAAAATTTGAACGTAACCTAATCTACGAGGTA 6450
GAAAATGAAGCACAGAGAAATAATCCATATAGCACAATTTGCTCCGTGGA 6500
ATGAGTCTACAAATACTGGGGTCTTTGTCTGGTTGCGGGAAACGGACAAC 6550
TCCAAAAGAAATACAAAAAAAAAACAAAAAAGGAAGAAAGAAAAGAAACT 6600
GATAATACAAGCCAACAAAATAACCTCAGTATGCTCAGACCTTGACACTG 6650
CATCGTTATGATACTCGTGCAAGATTTGTAATTTTTTTAGTGTAATTTGT 6700
TGCAGAGACTAGAAATTGCTGAAGTAAAAGATTAGATCTATGAGGAAATG 6750
TTTGTTACGATGTATAATAATGCTTCAGCTTTTCTTTTTTTTTTCTGCTG 6800
TTTTTTTCTTAATGTACTAGTATTTCATTTTTCTAAGTTTTCTTCTGTAT 6850
GGGTACAATCAACAGTTATTGCTTTCATCTCAGTAGTGTAAAGATACCCA 6900
CTTGAAAAAGGAAAAGAACTAACGAAGTCTTTCATTTCAAATTTAGATTC 6950
AGATGATCATTCGGCTTATATCAAAAAATAACAACAAAACTAAAAAAGGA 7000
AAATATCTTTAAAATCGGACCCCCACCATTGTGCCTTCATCAAACGCATA 7050
TATATATATCCAAACTCAGATTACCGGCACCCAGCCACCTAGATAATGCg 7100
agctccagcttttgttccctttagtgagggttaattgcgcgcttggcgta 7150
atcatgGTCATAGCTGTTTCCTGTGTGAAATTGTTATCCGCTCACAATTC 7200
CACACAACATAGGAGCCGGAAGCATAAAGTGTAAAGCCTGGGGTGCCTAA 7250
TGAGTGAGGTAACTCACATTAATTGCGTTGCGCTCACTGCCCGCTTTCCA 7300
GTCGGGAAACCTGTCGTGCCAGCTGCATTAATGAATCGGCCAACGCGCGG 7350
GGAGAGGCGGTTTGCGTATTGGGCGCTCTTCCGCTTCCTCGCTCACTGAC 7400
TCGCTGCGCTCGGTCGTTCGGCTGCGGCGAGCGGTATCAGCTCACTCAAA 7450
GGCGGTAATACGGTTATCCACAGAATCAGGGGATAACGCAGGAAAGAACA 7500
TGTGAGCAAAAGGCCAGCAAAAGGCCAGGAACCGTAAAAAGGCCGCGTTG 7550
CTGGCGTTTTTCCATAGGCTCCGCCCCCCTGACGAGCATCACAAAAATCG 7600
ACGCTCAAGTCAGAGGTGGCGAAACCCGACAGGACTATAAAGATACCAGG 7650

```
CGTTCCCCCCTGGAAGCTCCCTCGTGCGCTCTCCTGTTCCGACCCTGCCG 7700
CTTACCGGATACCTGTCCGCCTTTCTCCCTTCGGGAAGCGTGGCGCTTTC 7750
TCAATGCTCACGCTGTAGGTATCTCAGTTCGGTGTAGGTCGTTCGCTCCA 7800
AGCTGGGCTGTGTGCACGAACCCCCCGTTCAGCCCGACCGCTGCGCCTTA 7850
TCCGGTAACTATCGTCTTGAGTCCAACCCGGTAAGACACGACTTATCGCC 7900
ACTGGCAGCAGCCACTGGTAACAGGATTAGCAGAGCGAGGTATGTAGGCG 7950
GTGCTACAGAGTTCTTGAAGTGGTGGCCTAACTACGGCTACACTAGAAGG 8000
ACAGTATTTGGTATCTGCGCTCTGCTGAAGCCAGTTACCTTCGGAAAAAG 8050
AGTTGGTAGCTCTTGATCCGGCAAACAAACCACCGCTGGTAGCGGTGGTT 8100
TTTTTGTTTGCAAGCAGCAGATTACGCGCAGAAAAAAAGGATCTCAAGAA 8150
GATCCTTTGATCTTTTCTACGGGGTCTGACGCTCAGTGGAACGAAAACTC 8200
ACGTTAAGGGATTTTGGTCATGAGATTATCAAAAAGGATCTTCACCTAGA 8250
TCCTTTTAAATTAAAAATGAAGTTTTAAATCAATCTAAAGTATATATGAG 8300
TAAACTTGGTCTGACAGTTACCAATGCTTAATCAGTGAGGCACCTATCTC 8350
AGCGATCTGTCTATTTCGTTCATCCATAGTTGCCTGACTGCCCGTCGTGT 8400
AGATAACTACGATACGGGAGGGCTTACCATCTGGCCCCAGTGCTGCAATG 8450
ATACCGCGAGACCCACGCTCACCGGCTCCAGATTTATCAGCAATAAACCA 8500
GCCAGCCGGAAGGGCCGAGCGCAGAAGTGGTCCTGCAACTTTATCCGCCT 8550
CCATCCAGTCTATTAATTGTTGCCGGGAAGCTAGAGTAAGTAGTTCGCCA 8600
GTTAATAGTTTGCGCAACGTTGTTGCCATTGCTACAGGCATCGTGGTGTC 8650
ACGCTCGTCGTTTGGTATGGCTTCATTCAGCTCCGGTTCCCAACGATCAA 8700
GGCGAGTTACATGATCCCCCATGTTGTGAAAAAAGCGGTTAGCTCCTTC 8750
GGTCCTCCGATCGTTGTCAGAAGTAAGTTGGCCGCAGTGTTATCACTCAT 8800
GGTTATGGCAGCACTGCATAATTCTCTTACTGTCATGCCATCCGTAAGAT 8850
GCTTTTCTGTGACTGGTGAGTACTCAACCAAGTCATTCTGAGAATAGTGT 8900
ATGCGGCGACCGAGTTGCTCTTGCCCGGCGTCAATACGGGATAATACCGC 8950
GCCACATAGCAGAACTTTAAAAGTGCTCATCATTGGAAAACGTTCTTCGG 9000
GGCGAAAACTCTCAAGGATCTTACCGCTGTTGAGATCCAGTTCGATGTAA 9050
CCCACTCGTGCACCCAACTGATCTTCAGCATCTTTTACTTTCACCAGCGT 9100
TTCTGGGTGAGCAAAAACAGGAAGGCAAAATGCCGCAAAAAAGGGAATAA 9150
GGGCGACACGGAAATGTTGAATACTCATACTCTTCCTTTTTCAATATTAT 9200
TGAAGCATTTATCAGGGTTATTGTCTCATGAGCGGATACATATTTGAATG 9250
TATTTAGAAAAATAAACAAATAGGGGTTCCGCGCACATTTCCCCGAAAAG 9300
TGCCACCTGGGTCCTTTTCATCACGTGCTATAAAAATAATTATAATTTAA 9350
ATTTTTTAATATAAATATATAAATTAAAAATAGAAAGTAAAAAAAGAAAT 9400
TAAAGAAAAAATAGTTTTTGTTTTCCGAAGATGTAAAAGACTCTAGGGGG 9450
ATCGCCAACAAATACTACCTTTTATCTTGCTCTTCCTGCTCTCAGGTATT 9500
AATGCCGAATTGTTTCATCTTGTCTGTGTAGAAGACCACACACGAAAATC 9550
CTGTGATTTTACATTTTACTTATCGTTAATCGAATGTATATCTATTTAAT 9600
CTGCTTTTCTTGTCTAATAAATATATATGTAAAGTACGCTTTTTGTTGAA 9650
ATTTTTTAAACCTTTGTTTATTTTTTTTTCTTCATTCCGTAACTCTTCTA 9700
CCTTCTTTATTTACTTTCTAAAATCCAAATACAAAACATAAAAATAAATA 9750
AACACAGAGTAAATTCCCAAATTATTCCATCATTAAAAGATACGAGGCGC 9800
GTGTAAGTTACAGGCAAGCGATCCGTCCTAAGAAACCATTATTATCATGA 9850
CATTAACCTATAAAAATAGGCGTATCACGAGGCCCTTTCGTC ••• 9892
```

Sequence: pPDR1-Y208C.dna (Circular / 10,349 bp)
Enzymes: Unique 6+ Cutters (32 of 653 total)
Features: 32 total

Unique Cutters **Bold**

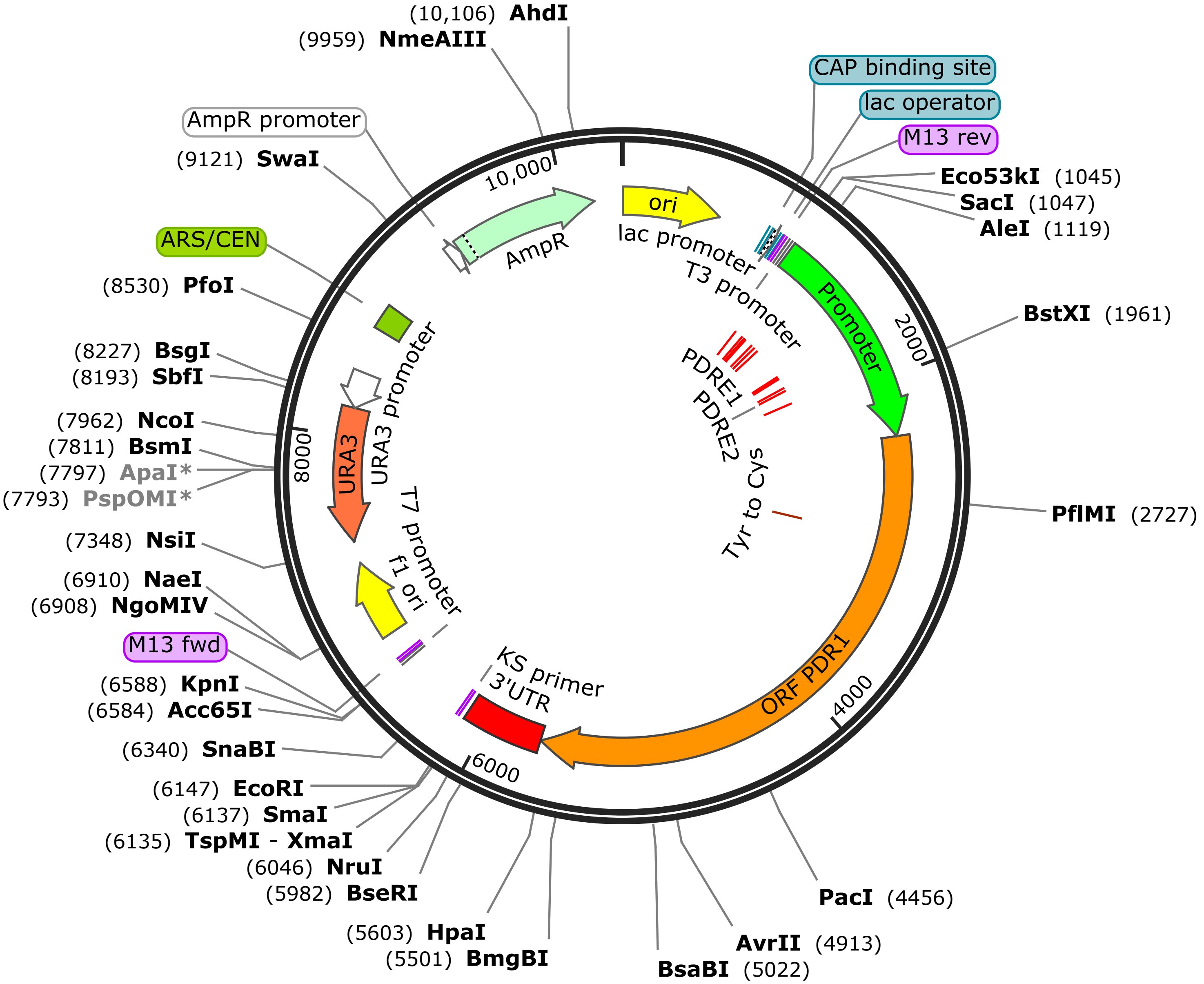


**pPDR1-Y208C**
10,349 bp

```
•••  ttgagatcctttttttctgcgcgtaatctgctgcttgcaaacaaa    45
     aaaaccaccgctaccagcggtggtttgtttgccggatcaagagct    90
     accaactctttttccgaaggtaactggcttcagcagagcgcagat   135
     accaaatactgttcttctagtgtagccgtagttaggccaccactt   180
     caagaactctgtagcaccgcctacatacctcgctctgctaatcct   225
     gttaccagtggctgctgccagtggcgataagtcgtgtcttaccgg   270
     gttggactcaagacgatagttaccggataaggcgcagcggtcggg   315
     ctgaacggggggttcgtgcacacagcccagcttggagcgaacgac   360
     ctacaccgaactgagatacctacagcgtgagctatgagaaagcgc   405
     cacgcttcccgaagggagaaaggcggacaggtatccggtaagcgg   450
     cagggtcggaacaggagagcgcacgagggagcttccagggggaaa   495
     cgcctggtatctttatagtcctgtcgggtttcgccacctctgact   540
     tgagcgtcgatttttgtgatgctcgtcaggggggcggagcctatg   585
     gaaaaacgccagcaacgcggcctttttacggttcctggccttttg   630
     ctggccttttgctcacatgttctttcctgcgttatcccctgattc   675
     tgtggataaccgtattaccgcctttgagtgagctgataccgctcg   720
     ccgcagccgaacgaccgagcgcagcgagtcagtgagcgaggaagc   765
     ggaagagcgcccaatacgcaaaccgcctctccccgcgcgttggcc   810
     gattcattaatgcagctggcacgacaggtttcccgactggaaagc   855
     gggcagtgagcgcaacgcaattaatgtgagttagctcactcatta   900
     ggcaccccaggctttacactttatgcttccggctcgtatgttgtg   945
     tggaattgtgagcggataacaatttcacacaggaaacagctatga   990
     ccatgattacgccaagcgcgcaattaaccctcactaaagggaaca  1035
     aaagctggagctcGCATTATCTAGGTGGCTGGGTGCCGGTAATCT  1080
     GAGTTTGGATATATATATATGCGTTTGATGAAGGCACAATGGTGG  1125
     GGGTCCGATTTTAAAGATATTTTCCTTTTTTAGTTTTGTTGTTAT  1170
     TTTTTGATATAAGCCGAATGATCATCTGAATCTAAATTTGAAATG  1215
     AAAGACTTCGTTAGTTCTTTTCCTTTTTCAAGTGGGTATCTTTAC  1260
     ACTACTGAGATGAAAGCAATAACTGTTGATTGTACCCATACAGAA  1305
     GAAAACTTAGAAAAATGAAATACTAGTACATTAAGAAAAAAACAG  1350
     CAGAAAAAAAAAAGAAAAGCTGAAGCATTATTATACATCGTAACA  1395
     AACATTTCCTCATAGATCTAATCTTTTACTTCAGCAATTTCTAGT  1440
     CTCTGCAACAAATTACACTAAAAAAATTACAAATCTTGCACGAGT  1485
     ATCATAACGATGCAGTGTCAAGGTCTGAGCATACTGAGGTTATTT  1530
     TGTTGGCTTGTATTATCAGTTTCTTTTCTTTCTTCCTTTTTTGTT  1575
     TTTTTTTTGTATTTCTTTTGGAGTTGTCCGTTTCCCGCAACCAGA  1620
     CAAAGACCCCAGTATTTGTAGACTCATTCCACGGAGCAAATTGTG  1665
     CTATATGGATTATTTCTCTGTGCTTCATTTTCTACCTCGTAGATT  1710
     AGGTTACGTTCAAATTTTAAGTGGCAAAACCATATCCTGATCGTC  1755
     AGTACTAAACCTGAGTTGTCCCGATGAGCCTCCTATTCCGTGGAA  1800
     AGATGGCTTTTCCTATTGATCTAAGTGGATAAGCAGGTATTGCGT  1845
     TGATCATTATAATTGTGGGTAAAACTAGGTATTAGGTCCTTCTAA  1890
     TAGTCATCTTTATTGTTGTAAAAATTGTTTCGATGGCGTATTCAT  1935
     AGAATCCGAACATATATACCAACCAAATGGATTTTAAAGGAAAAA  1980
     GCAAATACATTAAGGTCTGTTGTAGGAAAAGACCATATTGATTTA  2025
     TCTATTTAGTTGAGTAACAGAAATCACATATAGTTTTAAATTTTT  2070
     GGTAATCAAAGACACTTGAAACTATAAAACATTTGGTTGAAGAAG  2115
     AAGTATTAACTTTGCTTGGAGAAAAGCGTGCCCACACTTGCTTGG  2160
     TTTTCAGGATAATAACAAGCATAGAGGCGCTGTTATTACTTAAAC  2205
     AATTTTTAAGTAACACATTCAAACTTCCATTACTTCGTACCCCAT  2250
     ATCGTATTGCCATTGTGATATGGAATTAGTGTTTTATTCTGCCTT  2295
```

TTTTTTTAGAATATATTGGTAAAGTCATTCTTTAGCTACGTTATT 2340
GAGAGAATATGCAAACATTAGAAACTACATCAAAATCAAATCCAG 2385
GGGAAGTCAAAGCACAGAAGCCTAGTACAAGAAGAACAAAAGTTG 2430
GAAAAGCTTGTGATAGCTGTAGAAGGAGGAAAATAAAATGTAATG 2475
GGCTAAAACCTTGTCCATCTTGTACAATCTATGGCTGTGAATGTA 2520
CATATACTGATGCAAAATCGACAAAAAATCTCAAATCAAATGATG 2565
CAGGTAAATCAAAACCAACAGGGAGAGTATCAAAGAATAAAGAAA 2610
CTACTAGAGTCGACAAAGATATTAGGAAATTAGAGCAGCAGTATG 2655
TCCCTATTAATGCTAATATTCATGTTGGTCCCAGGTTCCCCTCCG 2700
AGAATATATTGAATGGATATCCACAATGTGGAGCACCACAGAACA 2745
ATGTTGTGGGTAATCCACTAGCGGTTAATACTCAATGCCATAGAG 2790
GTCTTTCTGAAACTCCTATGTCCTCAACATTCAAAGAATCTAACT 2835
TAAGAGATGATCGGCTACTACAGTCATCAGATACAGATGATATGA 2880
GGAATGGTGACTCGGAAGAAAGGGACTTGAAAGGGAGTGACAGCG 2925
AGAATGTCAAAAGTAAAGACAATAAAAGTGATCCTTTGATTATAT 2970
GCAAAGATGATACACATATTGAAAGCACGGTTAATAAACTAACAC 3015
AGGCAGTTAATGAACTCAAATCACTTCAAAATGCACCCAGTTCGA 3060
TAAAATCATCCATTGACGCCATTGAGTTACAACTTAGAAACATTT 3105
TAGACAACTGGAAACCAGAGGTAGATTTCGAGAAAGCAAAGATTA 3150
ATGAAAGTGCCACCACTAAGTCACTTGAAACAAACTTGCTGAGGA 3195
ATAAATACACTAATCACGTTCATTTAACAAGATTTAGGATATGGA 3240
TAGATTATAAAAATGCGAACAAAAACAATCATTTTATGGGAGAGT 3285
GTGGATTTAGTCTTGCAGAATCTTTTTTTGCTTCTAATCAGCCAT 3330
TGGTCGATGAATTGTTTGGGTTGTATTCCCAGGTAGAGGCCTTTT 3375
CTTTGCAAGGTCTTGGTTACTGTGTTCACCTTTATGAGCCATATA 3420
TGAAAACTGAGGAAGCGATAAAACTGATGAAAGAGACCTTATATA 3465
TTATACTACGGTTTATTGATATATGTGTTCACCATATCAATGAAG 3510
AGTCGATATCGATTGCCAACCCGTTAGAAACATATTTACGAAAAA 3555
AACATCTAATGCCTATGACTCCTACACCAAGGTCGTCCTATGGAA 3600
GTCCACAAAGTGCTAGTACAAAGAGCTTGGTAAGTAAGATAATAG 3645
AGAGAATACCGCAACCGTTTATTGAGAGTGTAACTAATGTGTCGA 3690
GTCTTCAACTATTAGATCTTCGAGATGACGAGTCAAAAATGTTTG 3735
GAACATTGCTGAACATGTGTAAGTCTATAAGGCGAAAATTTGACT 3780
CTGTTATGAGCGATTACGATTCCATTGTCACAGAAAAATCCGAAG 3825
GCGAACAAAATGATGGTAAAGTAACTGTAGCTGAGTTCACATCTT 3870
TGTGTGAAGCGGAAGAAATGCTCTTAGCATTATGCTATAACTATT 3915
ATAATCTGACGTTATACAGTTTCTTTGAATTTGGGACTAATATTG 3960
AATACATGGAACATCTGTTGCTTCTTCTTGAAGAACAGCTTGCTC 4005
TCGACGAATACTATGGTTTTGAAAAGGTCTTGAATGTAGCTGTTG 4050
CAAATGCTAAAAAAATGGGTTTCCACCGTTGGGAGTTTTACGTCG 4095
GCTATGAAGAGTCGACTGCTGAAAAGAGGCGGCTACTATGGTGGA 4140
AGTTATACAATTATGAAAAAGCCAGTACTATGAAGAAGGGTTTTT 4185
TTTCTGTGATTGATGATGCTACTGTCAACTGTTTATTACCTAAGA 4230
TTTTTAGAAACTTTGGCTATCTGGATAGGGTGGAGTTTCTAGAAA 4275
ATATTCAAAAGCCAATGGATCTTAGTGTGTTTTCCGATGTTCCAA 4320
TTTCTGTCCTTTGTAAATACGGTGAGTTGGCCCTTACAATAGTTA 4365
CCAGTGAGTTTCATGAAAAATTTTTATATGCTGATAGATACACTT 4410
CTATTCGAAATTCCGCGAAACCGCCGACATTAAAAAACCAATTAA 4455
TTAAGGAAATTGTGGATGGTATAGCTTATACAGAGACATCATATG 4500
AGGCAATCAGAAAGCAAACTGCAAAACTATGGGATATTGCATTAG 4545
GTAAGGTGACCAAAGATAAAATCAATAAAGAAGATACAGCAGCAG 4590

CTAGCAAATTTACTTTGAGTTATGAATATCACAGATTCAGGCTAA 4635
TCAATATGGCAGACAATTTAATTGCTAGACTAATGGTGAAACCAA 4680
AATCAGATTGGCTAATATCAGTCATGAAGGGGCATCTTAACAGAC 4725
TATATGAGCACTGGAAAGTAATGAATGAAATTATCCTAAGTATGG 4770
ACAACGATTATTCAATTGCAACAACGTTCGAATATTATGCACCAT 4815
CATGTCTGTGTTTAGCTACGCAGACTTTCCTTATTGTGAGGAATA 4860
TGGAAATGGATGATGTCAAGATGATGGTTGCAGTATATAAAAGAT 4905
TTCTTAACCTAGGAATGTTTTTGCAGAGTGCCAAAGTATGCAGCC 4950
TTGCCGATAGTCATACATTCAGAGATTTTTCTAGATCTTTTTCCT 4995
TTATTACGATAATTTCAAGATTGATGATAATCGAATTTATGCAAA 5040
TTAAAGAATTGACGAAGGTAGAGTTTATTGAGAAGTTTTCTGAAG 5085
TATGCCCTGACCTTGCAGATCTACCTCCGATGCTTCTAGATCCAA 5130
ACTCTTGCTTATATTTTTCATTGTTACAGCAGATTAAGAAATCTG 5175
GTTTTACGTTGTCATTCAAAAAAATTCTTGAAGACGCTAGAATGA 5220
TGGACTTCAATTACGACCGCAATTTGGACTCAGAGGCCATTAAAA 5265
AGTGCAATGGTGAATTTAGCAAGTCAATGCCTTCCTGTACCAATG 5310
TCTCAGATACCACCACCGCTGTTTCTGACAATAGTGCTAAGAAGA 5355
AAGCTTCAATGGGGTCGGCGAGGGTAAATTCAACTGATACACTAA 5400
CTGCATCTCCCTTATCGGGCTTAAGGAATCAAACGCAGTTGGATT 5445
CTAAAGACAGTGTTCCATCTCTCGAGGCTTATACACCAATTGATT 5490
CTGTCTCTGACGTGCCCACTGGGGAGATCAACGTTCCATTCCCTC 5535
CTGTTTATAATCAAAATGGATTGGATCAGCAAACCACTTATAATT 5580
TGGGAACTTTAGATGAGTTTGTTAACAAGGGAGATTTGAATGAAC 5625
TCTATAATAGCCTATGGGGTGACCTATTTTCTGATGTTTACTTGT 5670
GATGCTATTAATTTAGACAGTGTGCATAGCCTGTATAATTTTATA 5715
TTTAATCTGAAAAATCATTTTTTAGTGGTTATGTTGCATAATTTT 5760
ACATGAGACTTAGACTACCTCAAATTAGCGATAACACTACAATAT 5805
CTCTCATTAAAAAATTGTAGTTTAATATATGATGATTATTAAAAC 5850
GTATTGTACAAATTATAACTTATAAAAGAAAAACCCTTCGATTCT 5895
GAATGAAGATTCTAACAGCAAAACTGTTTGGGTTTATTCAATATT 5940
AAGGATACCCATTATCCTTTTTCCGTGAGGAGTGTTTCTAATATT 5985
ACCAACTAGTAGTGGTGTAACCAAGTCAACCAATTTCTTATACAT 6030
ATAAGGATTTGTGTCGCGACTAATATCCAATGCAGTCTGTAGCAC 6075
ATAGTTACCGAAGCTATCGTTTAACAGAGCTTGAACCTCAGCACT 6120
ACCTTCGTTGATCCCCCGGGCTGCAGgaattcgatatcaagctta 6165
tcgataccgtcgacctcgagaacacaacccacagctaccaccatc 6210
aacaatatttatatatataacgtacacatagaaatcacacaaaca 6255
gagtatttattcttaactacatgaactaccatcagaccgtctggg 6300
ccactatataatgtgccattcataaacgtgatcactttacgtagc 6345
aggcaaccccaggtgaaaatttttcagcgagctgccagattgtca 6390
ggtgaaaaactgaaaaaaacttctgggcgatgagcttgtggcggg 6435
aaattaagtatataaagcagttagtttcttctgcttcttgtggtt 6480
ctgggtattcttgtaatatttggaagaatagaaaatcaagactac 6525
tgcctttcttttcatattatagaggaagaaatacgcacgaacacg 6570
atatagaggtaaaggtacccaattcgccctatagtgagtcgtatt 6615
acgcgcgctcactggccgtcgttttacaacgtcgtgactgggaaa 6660
accctggcgttacccaacttaatcgccttgcagcacatccccctt 6705
tcgccagctggcgtaatagcgaagaggcccgcaccgatcgccctt 6750
cccaacagttgcgcagcctgaatggcgaatggacgcgccctgtag 6795
cggcgcattaagcgcggcgggtgtggtggttacgcgcagcgtgac 6840
cgctacacttgccagcgccctagcgcccgctcctttcgctttctt 6885

```
cccttcctttctcgccacgttcgccggctttccccgtcaagctct    6930
aaatcgggggctccctttagggttccgatttagtgctttacggca    6975
cctcgaccccaaaaaacttgattagggtgatggttcacgtagtgg    7020
gccatcgccctgatagacggtttttcgccctttgacgttggagtc    7065
cacgttctttaatagtggactcttgttccaaactggaacaacact    7110
caaccctatctcggtctattcttttgatttataagggattttgcc    7155
gatttcggcctattggttaaaaaatgagctgatttaacaaaaatt    7200
taacgcgaattttaacaaaatattaacgcttacaatttcctgatg    7245
cggtattttctccttacgcatctgtgcggtatttcacaccgcata    7290
gggtaataactgatataattaaattgaagctctaatttgtgagtt    7335
tagtatacatgcatttacttataatacagttttttagttttgctg    7380
gccgcatcttctcaaatatgcttcccagcctgcttttctgtaacg    7425
ttcaccctctaccttagcatcccttccctttgcaaatagtcctct    7470
tccaacaataataatgtcagatcctgtagagaccacatcatccac    7515
ggttctatactgttgacccaatgcgtctcccttgtcatctaaacc    7560
cacaccgggtgtcataatcaaccaatcgtaaccttcatctcttcc    7605
acccatgtctctttgagcaataaagccgataacaaaatctttgtc    7650
gctcttcgcaatgtcaacagtacccttagtatattctccagtaga    7695
tagggagcccttgcatgacaattctgctaacatcaaaaggcctct    7740
aggttcctttgttacttcttctgccgcctgcttcaaaccgctaac    7785
aatacctgggcccaccacaccgtgtgcattcgtaatgtctgccca    7830
ttctgctattctgtatacacccgcagagtactgcaatttgactgt    7875
attaccaatgtcagcaaattttctgtcttcgaagagtaaaaaatt    7920
gtacttggcggataatgcctttagcggcttaactgtgccctccat    7965
ggaaaaatcagtcaagatatccacatgtgtttttagtaaacaaat    8010
tttgggacctaatgcttcaactaactccagtaattccttggtggt    8055
acgaacatccaatgaagcacacaagtttgtttgcttttcgtgcat    8100
gatattaaatagcttggcagcaacaggactaggatgagtagcagc    8145
acgttccttatatgtagctttcgacatgatttatcttcgtttcct    8190
gcaggtttttgttctgtgcagttgggttaagaatactgggcaatt    8235
tcatgtttcttcaacactacatatgcgtatatataccaatctaag    8280
tctgtgctccttccttcgttcttccttctgttcggagattaccga    8325
atcaaaaaaatttcaaggaaaccgaaatcaaaaaaaagaataaaa    8370
aaaaaatgatgaattgaaaaggtggtatggtgcactctcagtaca    8415
atctgctctgatgccgcatagttaagccagccccgacacccgcca    8460
acacccgctgacgcgccctgacgggcttgtctgctcccggcatcc    8505
gcttacagacaagctgtgaccgtctccgggagctgcatgtgtcag    8550
aggttttcaccgtcatcaccgaaacgcgcgagacgaaagggcctc    8595
gtgatacgcctatttttataggttaatgtcatgataataatggtt    8640
tcttagcaggttagactaacatgcaaaaagtttatattcattttt    8685
gtagttattcttgttttcattaggatgattacacactgtggatgt    8730
ggattaatcgatcattgttaatttagtgtagctactaccatttca    8775
aacaaaacgtaaattcttgcagtcgtactggatctgtgaatctat    8820
tagtatatatgaattaaagtagcttgtacatattattctgttgaa    8865
tcatatcgcagagcattggttgaaatccaaaatataaaaatgtaa    8910
tatcacaaaaaataatactaattctaacattaatgggtcagattt    8955
ttagtgaatacttaaatttataatatcgtctatttaagctagcaa    9000
atggaacaacatttaaagtaagaacatcatatctacatgaaaatg    9045
tatatttcaatctgactaataacgcagagcacatctttcagtgat    9090
gtctgtcacatgatcaaaaagaattgtatttaaataattcataat    9135
aaaagcttaaaaaattacaataatgaaaataaagtaaataatgac    9180
```

```
atgggtaagagttccgaaatagaatcttagtgtacaaaacaaaaa    9225
ttgcatcattagagatccccacattcaaatatgtatccgctcatg    9270
agacaataaccctgataaatgcttcaataatattgaaaaaggaag    9315
agtatgagtattcaacatttccgtgtcgcccttattcccttttt     9360
gcggcattttgccttcctgtttttgctcacccagaaacgctggtg    9405
aaagtaaaagatgctgaagatcagttgggtgcacgagtgggttac    9450
atcgaactggatctcaacagcggtaagatccttgagagttttcgc    9495
cccgaagaacgttttccaatgatgagcacttttaaagttctgcta    9540
tgtggcgcggtattatcccgtattgacgccgggcaagagcaactc    9585
ggtcgccgcatacactattctcagaatgacttggttgagtactca    9630
ccagtcacagaaaagcatcttacggatggcatgacagtaagagaa    9675
ttatgcagtgctgccataaccatgagtgataacactgcggccaac    9720
ttacttctgacaacgatcggaggaccgaaggagctaaccgctttt    9765
ttgcacaacatgggggatcatgtaactcgccttgatcgttgggaa    9810
ccggagctgaatgaagccataccaaacgacgagcgtgacaccacg    9855
atgcctgtagcaatggcaacaacgttgcgcaaactattaactggc    9900
gaactacttactctagcttcccggcaacaattaatagactggatg    9945
gaggcggataaagttgcaggaccacttctgcgctcggcccttccg    9990
gctggctggtttattgctgataaatctggagccggtgagcgtggg    10,035
tctcgcggtatcattgcagcactggggccagatggtaagccctcc    10,080
cgtatcgtagttatctacacgacggggagtcaggcaactatggat    10,125
gaacgaaatagacagatcgctgagataggtgcctcactgattaag    10,170
cattggtaactgtcagaccaagtttactcatatatactttagatt    10,215
gatttaaaacttcattttaatttaaaaggatctaggtgaagatc    10,260
ctttttgataatctcatgaccaaaatcccttaacgtgagttttcg    10,305
ttccactgagcgtcagaccccgtagaaaagatcaaaggatcttc •••  10,349
```

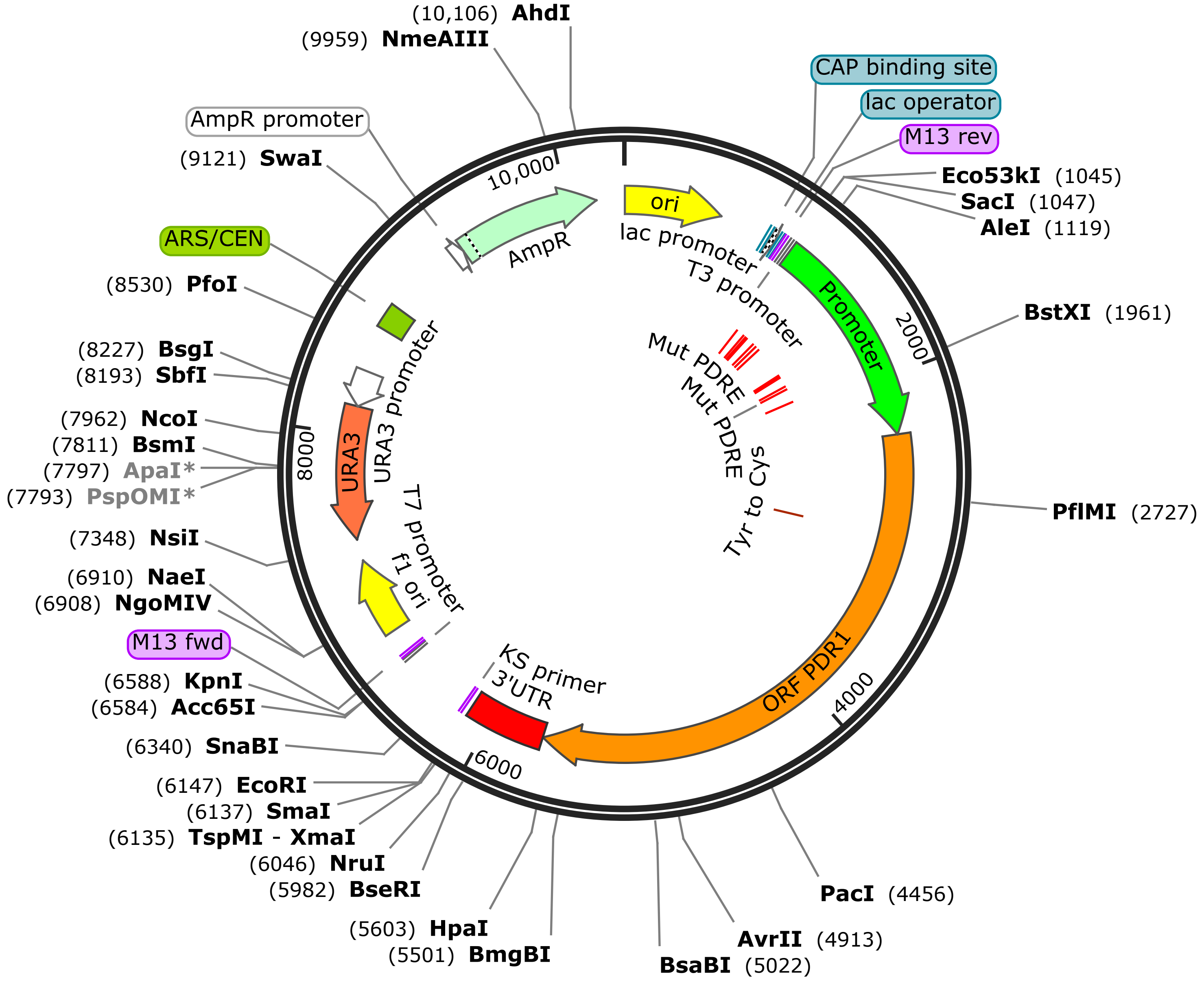


**pPDR1-Y208C-mPDRE**
10,349 bp

...
ttgagatcctttttttctgcgcgtaatctgctgcttgcaaacaaa 45
aaaaccaccgctaccagcggtggtttgtttgccggatcaagagct 90
accaactctttttccgaaggtaactggcttcagcagagcgcagat 135
accaaatactgttcttctagtgtagccgtagttaggccaccactt 180
caagaactctgtagcaccgcctacatacctcgctctgctaatcct 225
gttaccagtggctgctgccagtggcgataagtcgtgtcttaccgg 270
gttggactcaagacgatagttaccggataaggcgcagcggtcggg 315
ctgaacggggggttcgtgcacacagcccagcttggagcgaacgac 360
ctacaccgaactgagatacctacagcgtgagctatgagaaagcgc 405
cacgcttcccgaagggagaaaggcggacaggtatccggtaagcgg 450
cagggtcggaacaggagagcgcacgagggagcttccagggggaaa 495
cgcctggtatctttatagtcctgtcgggtttcgccacctctgact 540
tgagcgtcgatttttgtgatgctcgtcaggggggcggagcctatg 585
gaaaaacgccagcaacgcggcctttttacggttcctggccttttg 630
ctggccttttgctcacatgttctttcctgcgttatcccctgattc 675
tgtggataaccgtattaccgcctttgagtgagctgataccgctcg 720
ccgcagccgaacgaccgagcgcagcgagtcagtgagcgaggaagc 765
ggaagagcgcccaatacgcaaaccgcctctccccgcgcgttggcc 810
gattcattaatgcagctggcacgacaggtttcccgactggaaagc 855
gggcagtgagcgcaacgcaattaatgtgagttagctcactcatta 900
ggcaccccaggctttacactttatgcttccggctcgtatgttgtg 945
tggaattgtgagcggataacaatttcacacaggaaacagctatga 990
ccatgattacgccaagcgcgcaattaaccctcactaaagggaaca 1035
aaagctggagctcGCATTATCTAGGTGGCTGGGTGCCGGTAATCT 1080
GAGTTTGGATATATATATATGCGTTTGATGAAGGCACAATGGTGG 1125
GGGTCCGATTTTAAAGATATTTTCCTTTTTTAGTTTTGTTGTTAT 1170
TTTTTGATATAAGCCGAATGATCATCTGAATCTAAATTTGAAATG 1215
AAAGACTTCGTTAGTTCTTTTCCTTTTTCAAGTGGGTATCTTTAC 1260
ACTACTGAGATGAAAGCAATAACTGTTGATTGTACCCATACAGAA 1305
GAAAACTTAGAAAAATGAAATACTAGTACATTAAGAAAAAAACAG 1350
CAGAAAAAAAAAAGAAAAGCTGAAGCATTATTATACATCGTAACA 1395
AACATTTCCTCATAGATCTAATCTTTTACTTCAGCAATTTCTAGT 1440
CTCTGCAACAAATTACACTAAAAAAATTACAAATCTTGCACGAGT 1485
ATCATAACGATGCAGTGTCAAGGTCTGAGCATACTGAGGTTATTT 1530
TGTTGGCTTGTATTATCAGTTTCTTTTCTTTCTTCCTTTTTTGTT 1575
TTTTTTTTGTATTTCTTTTGGAGTTGTCCGTTTCCCGCAACCAGA 1620
CAAAGACCCCAGTATTTGTAGACTCATTaaaaaaAGCAAATTGTG 1665
CTATATGGATTATTTCTCTGTGCTTCATTTTCTACCTCGTAGATT 1710
AGGTTACGTTCAAATTTTAAGTGGCAAAACCATATCCTGATCGTC 1755
AGTACTAAACCTGAGTTGTCCCGATGAGCCTCCTATTaaaaaaAA 1800
AGATGGCTTTTCCTATTGATCTAAGTGGATAAGCAGGTATTGCGT 1845
TGATCATTATAATTGTGGGTAAAACTAGGTATTAGGTCCTTCTAA 1890
TAGTCATCTTTATTGTTGTAAAAATTGTTTCGATGGCGTATTCAT 1935
AGAATCCGAACATATATACCAACCAAATGGATTTTAAAGGAAAAA 1980
GCAAATACATTAAGGTCTGTTGTAGGAAAAGACCATATTGATTTA 2025
TCTATTTAGTTGAGTAACAGAAATCACATATAGTTTTAAATTTTT 2070
GGTAATCAAAGACACTTGAAACTATAAAACATTTGGTTGAAGAAG 2115
AAGTATTAACTTTGCTTGGAGAAAAGCGTGCCCACACTTGCTTGG 2160
TTTTCAGGATAATAACAAGCATAGAGGCGCTGTTATTACTTAAAC 2205
AATTTTTAAGTAACACATTCAAACTTCCATTACTTCGTACCCCAT 2250
ATCGTATTGCCATTGTGATATGGAATTAGTGTTTTATTCTGCCTT 2295

TTTTTTTAGAATATATTGGTAAAGTCATTCTTTAGCTACGTTATT 2340
GAGAGAATATGCAAACATTAGAAACTACATCAAAATCAAATCCAG 2385
GGGAAGTCAAAGCACAGAAGCCTAGTACAAGAAGAACAAAAGTTG 2430
GAAAAGCTTGTGATAGCTGTAGAAGGAGGAAAATAAAATGTAATG 2475
GGCTAAAACCTTGTCCATCTTGTACAATCTATGGCTGTGAATGTA 2520
CATATACTGATGCAAAATCGACAAAAAATCTCAAATCAAATGATG 2565
CAGGTAAATCAAAACCAACAGGGAGAGTATCAAAGAATAAAGAAA 2610
CTACTAGAGTCGACAAAGATATTAGGAAATTAGAGCAGCAGTATG 2655
TCCCTATTAATGCTAATATTCATGTTGGTCCCAGGTTCCCCTCCG 2700
AGAATATATTGAATGGATATCCACAATGTGGAGCACCACAGAACA 2745
ATGTTGTGGGTAATCCACTAGCGGTTAATACTCAATGCCATAGAG 2790
GTCTTTCTGAAACTCCTATGTCCTCAACATTCAAAGAATCTAACT 2835
TAAGAGATGATCGGCTACTACAGTCATCAGATACAGATGATATGA 2880
GGAATGGTGACTCGGAAGAAAGGGACTTGAAAGGGAGTGACAGCG 2925
AGAATGTCAAAAGTAAAGACAATAAAAGTGATCCTTTGATTATAT 2970
GCAAAGATGATACACATATTGAAAGCACGGTTAATAAACTAACAC 3015
AGGCAGTTAATGAACTCAAATCACTTCAAAATGCACCCAGTTCGA 3060
TAAAATCATCCATTGACGCCATTGAGTTACAACTTAGAAACATTT 3105
TAGACAACTGGAAACCAGAGGTAGATTTCGAGAAAGCAAAGATTA 3150
ATGAAAGTGCCACCACTAAGTCACTTGAAACAAACTTGCTGAGGA 3195
ATAAATACACTAATCACGTTCATTTAACAAGATTTAGGATATGGA 3240
TAGATTATAAAAATGCGAACAAAAACAATCATTTTATGGGAGAGT 3285
GTGGATTTAGTCTTGCAGAATCTTTTTTTGCTTCTAATCAGCCAT 3330
TGGTCGATGAATTGTTTGGGTTGTATTCCCAGGTAGAGGCCTTTT 3375
CTTTGCAAGGTCTTGGTTACTGTGTTCACCTTTATGAGCCATATA 3420
TGAAAACTGAGGAAGCGATAAAACTGATGAAAGAGACCTTATATA 3465
TTATACTACGGTTTATTGATATATGTGTTCACCATATCAATGAAG 3510
AGTCGATATCGATTGCCAACCCGTTAGAAACATATTTACGAAAAA 3555
AACATCTAATGCCTATGACTCCTACACCAAGGTCGTCCTATGGAA 3600
GTCCACAAAGTGCTAGTACAAAGAGCTTGGTAAGTAAGATAATAG 3645
AGAGAATACCGCAACCGTTTATTGAGAGTGTAACTAATGTGTCGA 3690
GTCTTCAACTATTAGATCTTCGAGATGACGAGTCAAAAATGTTTG 3735
GAACATTGCTGAACATGTGTAAGTCTATAAGGCGAAAATTTGACT 3780
CTGTTATGAGCGATTACGATTCCATTGTCACAGAAAAATCCGAAG 3825
GCGAACAAAATGATGGTAAAGTAACTGTAGCTGAGTTCACATCTT 3870
TGTGTGAAGCGGAAGAAATGCTCTTAGCATTATGCTATAACTATT 3915
ATAATCTGACGTTATACAGTTTCTTTGAATTTGGGACTAATATTG 3960
AATACATGGAACATCTGTTGCTTCTTCTTGAAGAACAGCTTGCTC 4005
TCGACGAATACTATGGTTTTGAAAAGGTCTTGAATGTAGCTGTTG 4050
CAAATGCTAAAAAAATGGGTTTCCACCGTTGGGAGTTTTACGTCG 4095
GCTATGAAGAGTCGACTGCTGAAAAGAGGCGGCTACTATGGTGGA 4140
AGTTATACAATTATGAAAAAGCCAGTACTATGAAGAAGGGTTTTT 4185
TTTCTGTGATTGATGATGCTACTGTCAACTGTTTATTACCTAAGA 4230
TTTTTAGAAACTTTGGCTATCTGGATAGGGTGGAGTTTCTAGAAA 4275
ATATTCAAAAGCCAATGGATCTTAGTGTGTTTTCCGATGTTCCAA 4320
TTTCTGTCCTTTGTAAATACGGTGAGTTGGCCCTTACAATAGTTA 4365
CCAGTGAGTTTCATGAAAAATTTTTATATGCTGATAGATACACTT 4410
CTATTCGAAATTCCGCGAAACCGCCGACATTAAAAACCAATTAA 4455
TTAAGGAAATTGTGGATGGTATAGCTTATACAGAGACATCATATG 4500
AGGCAATCAGAAAGCAAACTGCAAAACTATGGGATATTGCATTAG 4545
GTAAGGTGACCAAAGATAAAATCAATAAAGAAGATACAGCAGCAG 4590

```
CTAGCAAATTTACTTTGAGTTATGAATATCACAGATTCAGGCTAA 4635
TCAATATGGCAGACAATTTAATTGCTAGACTAATGGTGAAACCAA 4680
AATCAGATTGGCTAATATCAGTCATGAAGGGGCATCTTAACAGAC 4725
TATATGAGCACTGGAAAGTAATGAATGAAATTATCCTAAGTATGG 4770
ACAACGATTATTCAATTGCAACAACGTTCGAATATTATGCACCAT 4815
CATGTCTGTGTTTAGCTACGCAGACTTTCCTTATTGTGAGGAATA 4860
TGGAAATGGATGATGTCAAGATGATGGTTGCAGTATATAAAAGAT 4905
TTCTTAACCTAGGAATGTTTTTGCAGAGTGCCAAAGTATGCAGCC 4950
TTGCCGATAGTCATACATTCAGAGATTTTTCTAGATCTTTTTCCT 4995
TTATTACGATAATTTCAAGATTGATGATAATCGAATTTATGCAAA 5040
TTAAAGAATTGACGAAGGTAGAGTTTATTGAGAAGTTTTCTGAAG 5085
TATGCCCTGACCTTGCAGATCTACCTCCGATGCTTCTAGATCCAA 5130
ACTCTTGCTTATATTTTTCATTGTTACAGCAGATTAAGAAATCTG 5175
GTTTTACGTTGTCATTCAAAAAAATTCTTGAAGACGCTAGAATGA 5220
TGGACTTCAATTACGACCGCAATTTGGACTCAGAGGCCATTAAAA 5265
AGTGCAATGGTGAATTTAGCAAGTCAATGCCTTCCTGTACCAATG 5310
TCTCAGATACCACCACCGCTGTTTCTGACAATAGTGCTAAGAAGA 5355
AAGCTTCAATGGGGTCGGCGAGGGTAAATTCAACTGATACACTAA 5400
CTGCATCTCCCTTATCGGGCTTAAGGAATCAAACGCAGTTGGATT 5445
CTAAAGACAGTGTTCCATCTCTCGAGGCTTATACACCAATTGATT 5490
CTGTCTCTGACGTGCCCACTGGGGAGATCAACGTTCCATTCCCTC 5535
CTGTTTATAATCAAAATGGATTGGATCAGCAAACCACTTATAATT 5580
TGGGAACTTTAGATGAGTTTGTTAACAAGGGAGATTTGAATGAAC 5625
TCTATAATAGCCTATGGGGTGACCTATTTTCTGATGTTTACTTGT 5670
GATGCTATTAATTTAGACAGTGTGCATAGCCTGTATAATTTTATA 5715
TTTAATCTGAAAAATCATTTTTTAGTGGTTATGTTGCATAATTTT 5760
ACATGAGACTTAGACTACCTCAAATTAGCGATAACACTACAATAT 5805
CTCTCATTAAAAAATTGTAGTTTAATATATGATGATTATTAAAAC 5850
GTATTGTACAAATTATAACTTATAAAAGAAAAACCCTTCGATTCT 5895
GAATGAAGATTCTAACAGCAAAACTGTTTGGGTTTATTCAATATT 5940
AAGGATACCCATTATCCTTTTTCCGTGAGGAGTGTTTCTAATATT 5985
ACCAACTAGTAGTGGTGTAACCAAGTCAACCAATTTCTTATACAT 6030
ATAAGGATTTGTGTCGCGACTAATATCCAATGCAGTCTGTAGCAC 6075
ATAGTTACCGAAGCTATCGTTTAACAGAGCTTGAACCTCAGCACT 6120
ACCTTCGTTGATCCCCCGGGCTGCAGgaattcgatatcaagctta 6165
tcgataccgtcgacctcgagaacacaacccacagctaccaccatc 6210
aacaatatttatatatataacgtacacatagaaatcacacaaaca 6255
gagtatttattcttaactacatgaactaccatcagaccgtctggg 6300
ccactatataatgtgccattcataaacgtgatcactttacgtagc 6345
aggcaaccccaggtgaaaatttttcagcgagctgccagattgtca 6390
ggtgaaaaactgaaaaaaacttctgggcgatgagcttgtggcggg 6435
aaattaagtatataaagcagttagtttcttctgcttcttgtggtt 6480
ctgggtattcttgtaatatttggaagaatagaaaatcaagactac 6525
tgcctttcttttcatattatagaggaagaaatacgcacgaacacg 6570
atatagaggtaaaggtacccaattcgccctatagtgagtcgtatt 6615
acgcgcgctcactggccgtcgttttacaacgtcgtgactgggaaa 6660
accctggcgttacccaacttaatcgccttgcagcacatccccctt 6705
tcgccagctggcgtaatagcgaagaggcccgcaccgatcgccctt 6750
cccaacagttgcgcagcctgaatggcgaatggacgcgccctgtag 6795
cggcgcattaagcgcggcgggtgtggtggttacgcgcagcgtgac 6840
cgctacacttgccagcgccctagcgcccgctcctttcgctttctt 6885
```

ccctttcctttctcgccacgttcgccggctttccccgtcaagctct 6930
aaatcgggggctccctttagggttccgatttagtgctttacggca 6975
cctcgaccccaaaaaacttgattagggtgatggttcacgtagtgg 7020
gccatcgccctgatagacggtttttcgccctttgacgttggagtc 7065
cacgttctttaatagtggactcttgttccaaactggaacaacact 7110
caaccctatctcggtctattcttttgatttataagggattttgcc 7155
gatttcggcctattggttaaaaaatgagctgatttaacaaaaatt 7200
taacgcgaattttaacaaaatattaacgcttacaatttcctgatg 7245
cggtattttctccttacgcatctgtgcggtatttcacaccgcata 7290
gggtaataactgatataattaaattgaagctctaatttgtgagtt 7335
tagtatacatgcatttacttataatacagttttttagttttgctg 7380
gccgcatcttctcaaatatgcttcccagcctgcttttctgtaacg 7425
ttcaccctctaccttagcatcccttccctttgcaaatagtcctct 7470
tccaacaataataatgtcagatcctgtagagaccacatcatccac 7515
ggttctatactgttgacccaatgcgtctcccttgtcatctaaacc 7560
cacaccgggtgtcataatcaaccaatcgtaaccttcatctcttcc 7605
acccatgtctctttgagcaataaagccgataacaaaatctttgtc 7650
gctcttcgcaatgtcaacagtacccttagtatattctccagtaga 7695
tagggagcccttgcatgacaattctgctaacatcaaaaggcctct 7740
aggttcctttgttacttcttctgccgcctgcttcaaaccgctaac 7785
aatacctgggcccaccacaccgtgtgcattcgtaatgtctgccca 7830
ttctgctattctgtatacacccgcagagtactgcaatttgactgt 7875
attaccaatgtcagcaaattttctgtcttcgaagagtaaaaaatt 7920
gtacttggcggataatgcctttagcggcttaactgtgccctccat 7965
ggaaaaatcagtcaagatatccacatgtgtttttagtaaacaaat 8010
tttgggacctaatgcttcaactaactccagtaattccttggtggt 8055
acgaacatccaatgaagcacacaagtttgtttgcttttcgtgcat 8100
gatattaaatagcttggcagcaacaggactaggatgagtagcagc 8145
acgttccttatatgtagctttcgacatgatttatcttcgtttcct 8190
gcaggtttttgttctgtgcagttgggttaagaatactgggcaatt 8235
tcatgtttcttcaacactacatatgcgtatatataccaatctaag 8280
tctgtgctccttccttcgttcttccttctgttcggagattaccga 8325
atcaaaaaaatttcaaggaaaccgaaatcaaaaaaaagaataaaa 8370
aaaaaatgatgaattgaaaaggtggtatggtgcactctcagtaca 8415
atctgctctgatgccgcatagttaagccagccccgacacccgcca 8460
acacccgctgacgcgccctgacgggcttgtctgctcccggcatcc 8505
gcttacagacaagctgtgaccgtctccgggagctgcatgtgtcag 8550
aggttttcaccgtcatcaccgaaacgcgcgagacgaaagggcctc 8595
gtgatacgcctatttttataggttaatgtcatgataataatggtt 8640
tcttagcaggttagactaacatgcaaaagttatattcattttt 8685
gtagttattcttgtttcattaggatgattacacactgtggatgt 8730
ggattaatcgatcattgttaatttagtgtagctactaccatttca 8775
aacaaaacgtaaattcttgcagtcgtactggatctgtgaatctat 8820
tagtatatatgaattaaagtagcttgtacatattattctgttgaa 8865
tcatatcgcagagcattggttgaaatccaaaatataaaaatgtaa 8910
tatcacaaaaaataatactaattctaacattaatgggtcagattt 8955
ttagtgaatacttaaatttataatatcgtctatttaagctagcaa 9000
atggaacaacatttaaagtaagaacatcatatctacatgaaaatg 9045
tatatttcaatctgactaataacgcagagcacatctttcagtgat 9090
gtctgtcacatgatcaaaagaattgtatttaaataattcataat 9135
aaaagcttaaaaaattacaataatgaaaataaagtaaataatgac 9180

```
atgggtaagagttccgaaatagaatcttagtgtacaaaacaaaaa    9225
ttgcatcattagagatccccacattcaaatatgtatccgctcatg    9270
agacaataaccctgataaatgcttcaataatattgaaaaaggaag    9315
agtatgagtattcaacatttccgtgtcgcccttattcccttttt    9360
gcggcattttgccttcctgtttttgctcacccagaaacgctggtg    9405
aaagtaaaagatgctgaagatcagttgggtgcacgagtgggttac    9450
atcgaactggatctcaacagcggtaagatccttgagagttttcgc    9495
cccgaagaacgttttccaatgatgagcacttttaaagttctgcta    9540
tgtggcgcggtattatcccgtattgacgccgggcaagagcaactc    9585
ggtcgccgcatacactattctcagaatgacttggttgagtactca    9630
ccagtcacagaaaagcatcttacggatggcatgacagtaagagaa    9675
ttatgcagtgctgccataaccatgagtgataacactgcggccaac    9720
ttacttctgacaacgatcggaggaccgaaggagctaaccgctttt    9765
ttgcacaacatgggggatcatgtaactcgccttgatcgttgggaa    9810
ccggagctgaatgaagccataccaaacgacgagcgtgacaccacg    9855
atgcctgtagcaatggcaacaacgttgcgcaaactattaactggc    9900
gaactacttactctagcttcccggcaacaattaatagactggatg    9945
gaggcggataaagttgcaggaccacttctgcgctcggcccttccg    9990
gctggctggtttattgctgataaatctggagccggtgagcgtggg    10,035
tctcgcggtatcattgcagcactggggccagatggtaagccctcc    10,080
cgtatcgtagttatctacacgacggggagtcaggcaactatggat    10,125
gaacgaaatagacagatcgctgagataggtgcctcactgattaag    10,170
cattggtaactgtcagaccaagtttactcatatatactttagatt    10,215
gatttaaaacttcatttttaatttaaaaggatctaggtgaagatc    10,260
cctttttgataatctcatgaccaaaatcccttaacgtgagttttcg    10,305
ttccactgagcgtcagaccccgtagaaaagatcaaaggatcttc •••  10,349
```

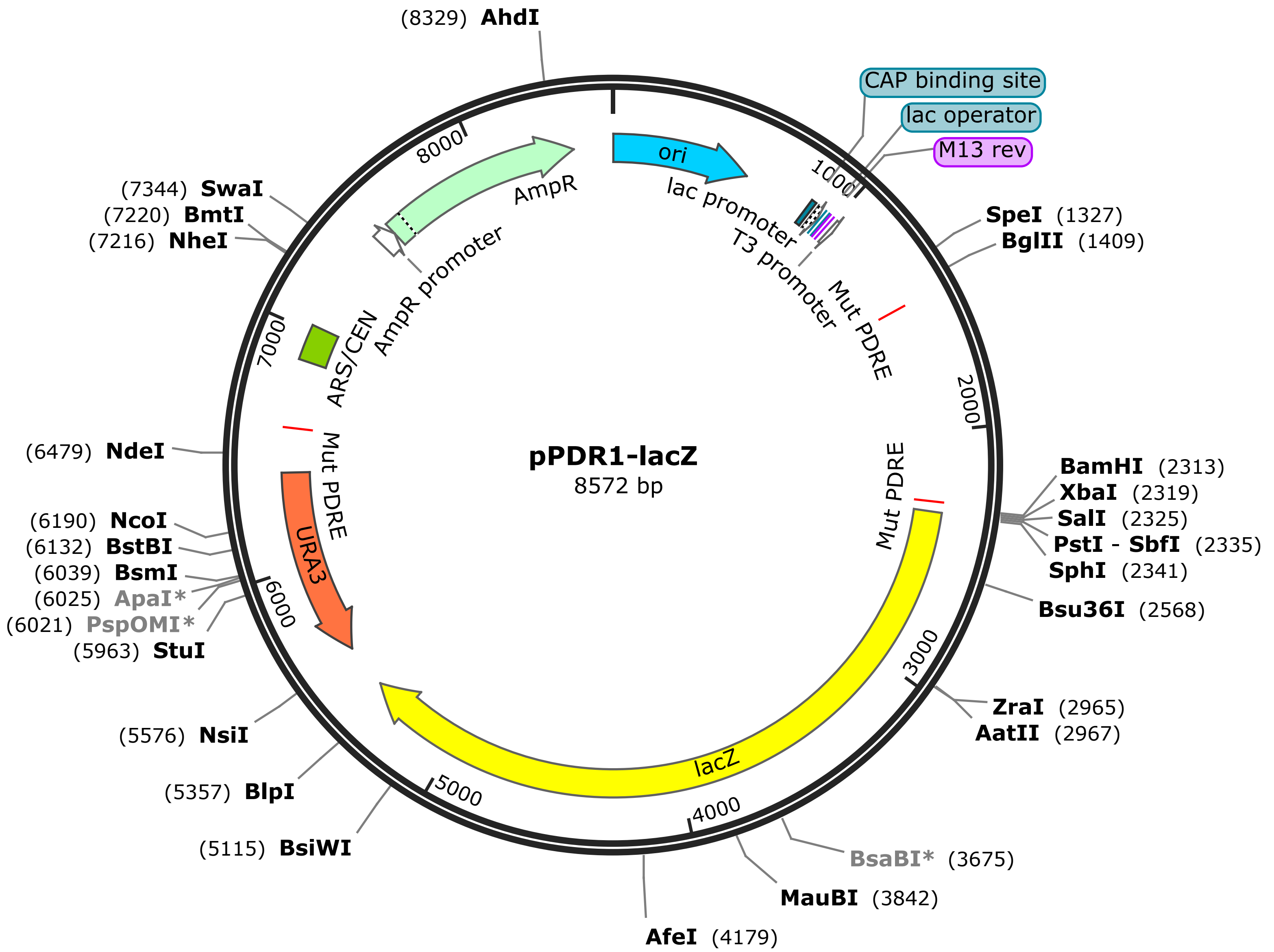
pPDR1-lacZ
8572 bp
(8329) AhdI
CAP binding site
lac operator
M13 rev
SpeI (1327)
BglII (1409)
BamHI (2313)
XbaI (2319)
SalI (2325)
PstI - SbfI (2335)
SphI (2341)
Bsu36I (2568)
ZraI (2965)
AatII (2967)
BsaBI* (3675)
MauBI (3842)
AfeI (4179)
(5115) BsiWI
(5357) BlpI
(5576) NsiI
(5963) StuI
(6021) PspOMI*
(6025) ApaI*
(6039) BsmI
(6132) BstBI
(6190) NcoI
(6479) NdeI
(7216) NheI
(7220) BmtI
(7344) SwaI
ori
lac promoter
T3 promoter
Mut PDRE
Mut PDRE
lacZ
URA3
Mut PDRE
ARS/CEN
AmpR promoter
AmpR
1000
2000
3000
4000
5000
6000
7000
8000

TTGAGATCCTTTTTTTCTGCGCGTAATCTGCTGCTTGCAAACAAAAAAAC 50
CACCGCTACCAGCGGTGGTTTGTTTGCCGGATCAAGAGCTACCAACTCTT 100
TTTCCGAAGGTAACTGGCTTCAGCAGAGCGCAGATACCAAATACTGTTCT 150
TCTAGTGTAGCCGTAGTTAGGCCACCACTTCAAGAACTCTGTAGCACCGC 200
CTACATACCTCGCTCTGCTAATCCTGTTACCAGTGGCTGCTGCCAGTGGC 250
GATAAGTCGTGTCTTACCGGGTTGGACTCAAGACGATAGTTACCGGATAA 300
GGCGCAGCGGTCGGGCTGAACGGGGGGTTCGTGCACACAGCCCAGCTTGG 350
AGCGAACGACCTACACCGAACTGAGATACCTACAGCGTGAGCTATGAGAA 400
AGCGCCACGCTTCCCGAAGGGAGAAAGGCGGACAGGTATCCGGTAAGCGG 450
CAGGGTCGGAACAGGAGAGCGCACGAGGGAGCTTCCAGGGGGAAACGCCT 500
GGTATCTTTATAGTCCTGTCGGGTTTCGCCACCTCTGACTTGAGCGTCGA 550
TTTTTGTGATGCTCGTCAGGGGGGCGGAGCCTATGGAAAAACGCCAGCAA 600
CGCGGCCTTTTTACGGTTCCTGGCCTTTTGCTGGCCTTTTGCTCACATGT 650
TCTTTCCTGCGTTATCCCCTGATTCTGTGGATAACCGTATTACCGCCTTT 700
GAGTGAGCTGATACCGCTCGCCGCAGCCGAACGACCGAGCGCAGCGAGTC 750
AGTGAGCGAGGAAGCGGAAGAGCGCCCAATACGCAAACCGCCTCTCCCCG 800
CGCGTTGGCCGATTCATTAATGCAGCTGGCACGACAGGTTTCCCGACTGG 850
AAAGCGGGCAGTGAGCGCAACGCAATTAATGTGAGTTAGCTCACTCATTA 900
GGCACCCCAGGCTTTACACTTTATGCTTCCGGCTCGTATGTTGTGTGGAA 950
TTGTGAGCGGATAACAATTTCACACAGGAAACAGCTATGACCATGATTAC 1000
GCCAAGCGCGCAATTAACCCTCACTAAAGGGAACAAAAGCTGGAGCTCGC 1050
ATTATCTAGGTGGCTGGGTGCCGGTAATCTGAGTTTGGATATATATATAT 1100
GCGTTTGATGAAGGCACAATGGTGGGGGTCCGATTTTAAAGATATTTTCC 1150
TTTTTTAGTTTTGTTGTTATTTTTTGATATAAGCCGAATGATCATCTGAA 1200
TCTAAATTTGAAATGAAAGACTTCGTTAGTTCTTTTCCTTTTTCAAGTGG 1250
GTATCTTTACACTACTGAGATGAAAGCAATAACTGTTGATTGTACCCATA 1300
CAGAAGAAAACTTAGAAAAATGAAATACTAGTACATTAAGAAAAAAACAG 1350
CAGAAAAAAAAAAGAAAAGCTGAAGCATTATTATACATCGTAACAAACAT 1400
TTCCTCATAGATCTAATCTTTTACTTCAGCAATTTCTAGTCTCTGCAACA 1450
AATTACACTAAAAAAATTACAAATCTTGCACGAGTATCATAACGATGCAG 1500
TGTCAAGGTCTGAGCATACTGAGGTTATTTTGTTGGCTTGTATTATCAGT 1550
TTCTTTTCTTTCTTCCTTTTTTGTTTTTTTTTTGTATTTCTTTTGGAGTT 1600
GTCCGTTTCCCGCAACCAGACAAAGACCCCAGTATTTGTAGACTCATTCC 1650
ACGGAGCAAATTGTGCTATATGGATTATTTCTCTGTGCTTCATTTTCTAC 1700
CTCGTAGATTAGGTTACGTTCAAATTTTAAGTGGCAAAACCATATCCTGA 1750
TCGTCAGTACTAAACCTGAGTTGTCCCGATGAGCCTCCTATTCCGTGGAA 1800
AGATGGCTTTTCCTATTGATCTAAGTGGATAAGCAGGTATTGCGTTGATC 1850
ATTATAATTGTGGGTAAAACTAGGTATTAGGTCCTTCTAATAGTCATCTT 1900
TATTGTTGTAAAAATTGTTTCGATGGCGTATTCATAGAATCCGAACATAT 1950
ATACCAACCAAATGGATTTTAAAGGAAAAAGCAAATACATTAAGGTCTGT 2000
TGTAGGAAAAGACCATATTGATTTATCTATTTAGTTGAGTAACAGAAATC 2050
ACATATAGTTTTAAATTTTTGGTAATCAAAGACACTTGAAACTATAAAAC 2100
ATTTGGTTGAAGAAGAAGTATTAACTTTGCTTGGAGAAAAGCGTGCCCAC 2150
ACTTGCTTGGTTTTCAGGATAATAACAAGCATAGAGGCGCTGTTATTACT 2200
TAAACAATTTTTAAGTAACACATTCAAACTTCCATTACTTCGTACCCCAT 2250
ATCGTATTGCCATTGTGATATGGAATTAGTGTTTTATTCTGCCTTTTTTT 2300
TTAGAATATATTGGATCCTCTAGAGTCGACCTGCAGGCATGCAAGCTTGC 2350
GATCCCGTCGTTTTACAACGTCGTGACTGGGAAAACCCTGGCGTTACCCA 2400
ACTTAATCGCCTTGCAGCACATCCCCCTTTCGCCAGCTGGCGTAATAGCG 2450
AAGAGGCCCGCACCGATCGCCCTTCCCAACAGTTGCGCAGCCTGAATGGC 2500
GAATGGCGCTTTGCCTGGTTTCCGGCACCAGAAGCGGTGCCGGAAAGCTG 2550

GCTGGAGTGCGATCTTCCTGAGGCCGATACTGTCGTCGTCCCCTCAAACT 2600
GGCAGATGCACGGTTACGATGCGCCCATCTACACCAACGTGACCTATCCC 2650
ATTACGGTCAATCCGCCGTTTGTTCCCACGGAGAATCCGACGGGTTGTTA 2700
CTCGCTCACATTTAATGTTGATGAAAGCTGGCTACAGGAAGGCCAGACGC 2750
GAATTATTTTTGATGGCGTTAACTCGGCGTTTCATCTGTGGTGCAACGGG 2800
CGCTGGGTCGGTTACGGCCAGGACAGTCGTTTGCCGTCTGAATTTGACCT 2850
GAGCGCATTTTTACGCGCCGGAGAAAACCGCCTCGCGGTGATGGTGCTGC 2900
GCTGGAGTGACGGCAGTTATCTGGAAGATCAGGATATGTGGCGGATGAGC 2950
GGCATTTTCCGTGACGTCTCGTTGCTGCATAAACCGACTACACAAATCAG 3000
CGATTTCCATGTTGCCACTCGCTTTAATGATGATTTCAGCCGCGCTGTAC 3050
TGGAGGCTGAAGTTCAGATGTGCGGCGAGTTGCGTGACTACCTACGGGTA 3100
ACAGTTTCTTTATGGCAGGGTGAAACGCAGGTCGCCAGCGGCACCGCGCC 3150
TTTCGGCGGTGAAATTATCGATGAGCGTGGTGGTTATGCCGATCGCGTCA 3200
CACTACGTCTGAACGTCGAAAACCCGAAACTGTGGAGCGCCGAAATCCCG 3250
AATCTCTATCGTGCGGTGGTTGAACTGCACACCGCCGACGGCACGCTGAT 3300
TGAAGCAGAAGCCTGCGATGTCGGTTTCCGCGAGGTGCGGATTGAAAATG 3350
GTCTGCTGCTGCTGAACGGCAAGCCGTTGCTGATTCGAGGCGTTAACCGT 3400
CACGAGCATCATCCTCTGCATGGTCAGGTCATGGATGAGCAGACGATGGT 3450
GCAGGATATCCTGCTGATGAAGCAGAACAACTTTAACGCCGTGCGCTGTT 3500
CGCATTATCCGAACCATCCGCTGTGGTACACGCTGTGCGACCGCTACGGC 3550
CTGTATGTGGTGGATGAAGCCAATATTGAAACCCACGGCATGGTGCCAAT 3600
GAATCGTCTGACCGATGATCCGCGCTGGCTACCGGCGATGAGCGAACGCG 3650
TAACGCGAATGGTGCAGCGCGATCGTAATCACCCGAGTGTGATCATCTGG 3700
TCGCTGGGGAATGAATCAGGCCACGGCGCTAATCACGACGCGCTGTATCG 3750
CTGGATCAAATCTGTCGATCCTTCCCGCCCGGTGCAGTATGAAGGCGGCG 3800
GAGCCGACACCACGGCCACCGATATTATTTGCCCGATGTACGCGCGCGTG 3850
GATGAAGACCAGCCCTTCCCGGCTGTGCCGAAATGGTCCATCAAAAAATG 3900
GCTTTCGCTACCTGGAGAGACGCGCCCGCTGATCCTTTGCGAATACGCCC 3950
ACGCGATGGGTAACAGTCTTGGCGGTTTCGCTAAATACTGGCAGGCGTTT 4000
CGTCAGTATCCCCGTTTACAGGGCGGCTTCGTCTGGGACTGGGTGGATCA 4050
GTCGCTGATTAAATATGATGAAAACGGCAACCCGTGGTCGGCTTACGGCG 4100
GTGATTTTGGCGATACGCCGAACGATCGCCAGTTCTGTATGAACGGTCTG 4150
GTCTTTGCCGACCGCACGCCGCATCCAGCGCTGACGGAAGCAAAACACCA 4200
GCAGCAGTTTTTCCAGTTCCGTTTATCCGGGCAAACCATCGAAGTGACCA 4250
GCGAATACCTGTTCCGTCATAGCGATAACGAGCTCCTGCACTGGATGGTG 4300
GCGCTGGATGGTAAGCCGCTGGCAAGCGGTGAAGTGCCTCTGGATGTCGC 4350
TCCACAAGGTAAACAGTTGATTGAACTGCCTGAACTACCGCAGCCGGAGA 4400
GCGCCGGGCAACTCTGGCTCACAGTACGCGTAGTGCAACCGAACGCGACC 4450
GCATGGTCAGAAGCCGGGCACATCAGCGCCTGGCAGCAGTGGCGTCTGGC 4500
GGAAAACCTCAGTGTGACGCTCCCCGCCGCGTCCCACGCCATCCCGCATC 4550
TGACCACCAGCGAAATGGATTTTTGCATCGAGCTGGGTAATAAGCGTTGG 4600
CAATTTAACCGCCAGTCAGGCTTTCTTTCACAGATGTGGATTGGCGATAA 4650
AAAACAACTGCTGACGCCGCTGCGCGATCAGTTCACCCGTGCACCGCTGG 4700
ATAACGACATTGGCGTAAGTGAAGCGACCCGCATTGACCCTAACGCCTGG 4750
GTCGAACGCTGGAAGGCGGCGGGCCATTACCAGGCCGAAGCAGCGTTGTT 4800
GCAGTGCACGGCAGATACACTTGCTGATGCGGTGCTGATTACGACCGCTC 4850
ACGCGTGGCAGCATCAGGGGAAAACCTTATTTATCAGCCGGAAAACCTAC 4900
CGGATTGATGGTAGTGGTCAAATGGCGATTACCGTTGATGTTGAAGTGGC 4950
GAGCGATACACCGCATCCGGCGCGGATTGGCCTGAACTGCCAGCTGGCGC 5000
AGGTAGCAGAGCGGGTAAACTGGCTCGGATTAGGGCCGCAAGAAAACTAT 5050
CCCGACCGCCTTACTGCCGCCTGTTTTGACCGCTGGGATCTGCCATTGTC 5100

AGACATGTATATCCCGTACGTCTTCCCGAGCGAAAACGGTCTGCGCTGCG 5150
GGACGCGCGAATTGAATTATGGCCCACACCAGTGGCGCGGCGACTTCCAG 5200
TTCAACATCAGCCGCTACAGTCAACAGCAACTGATGGAAACCAGCCATCG 5250
CCATCTGCTGCACGCGGAAGAAGGCACATGGCTGAATATCGACGGTTTCC 5300
ACATGGGGATTGGTGGCGACGACTCCTGGAGCCCGTCAGTATCGGCGGAA 5350
TTACAGCTGAGCGCCGGTCGCTACCATTACCAGTTGGTCTGGTGTCAAAA 5400
ATAATAATAACCGGGCAGGCCATGTCTGCCCGTATTTCGCGTAAGGAAAT 5450
CCATTATGTACTATTTCCTGATGCGGTATTTTCTCCTTACGCATCTGTGC 5500
GGTATTTCACACCGCATAGGGTAATAACTGATATAATTAAATTGAAGCTC 5550
TAATTTGTGAGTTTAGTATACATGCATTTACTTATAATACAGTTTTTTAG 5600
TTTTGCTGGCCGCATCTTCTCAAATATGCTTCCCAGCCTGCTTTTCTGTA 5650
ACGTTCACCCTCTACCTTAGCATCCCTTCCCTTTGCAAATAGTCCTCTTC 5700
CAACAATAATAATGTCAGATCCTGTAGAGACCACATCATCCACGGTTCTA 5750
TACTGTTGACCCAATGCGTCTCCCTTGTCATCTAAACCCACACCGGGTGT 5800
CATAATCAACCAATCGTAACCTTCATCTCTTCCACCCATGTCTCTTTGAG 5850
CAATAAAGCCGATAACAAAATCTTTGTCGCTCTTCGCAATGTCAACAGTA 5900
CCCTTAGTATATTCTCCAGTAGATAGGGAGCCCTTGCATGACAATTCTGC 5950
TAACATCAAAAGGCCTCTAGGTTCCTTTGTTACTTCTTCTGCCGCCTGCT 6000
TCAAACCGCTAACAATACCTGGGCCCACCACACCGTGTGCATTCGTAATG 6050
TCTGCCCATTCTGCTATTCTGTATACACCCGCAGAGTACTGCAATTTGAC 6100
TGTATTACCAATGTCAGCAAATTTTCTGTCTTCGAAGAGTAAAAAATTGT 6150
ACTTGGCGGATAATGCCTTTAGCGGCTTAACTGTGCCCTCCATGGAAAAA 6200
TCAGTCAAGATATCCACATGTGTTTTTAGTAAACAAATTTTGGGACCTAA 6250
TGCTTCAACTAACTCCAGTAATTCCTTGGTGGTACGAACATCCAATGAAG 6300
CACACAAGTTTGTTTGCTTTTCGTGCATGATATTAAATAGCTTGGCAGCA 6350
ACAGGACTAGGATGAGTAGCAGCACGTTCCTTATATGTAGCTTTCGACAT 6400
GATTTATCTTCGTTTCGGTTTTTGTTCTGTGCAGTTGGGTTAAGAATACT 6450
GGGCAATTTCATGTTTCTTCAACACTACATATGCGTATATATACCAATCT 6500
AAGTCTGTGCTCCTTCCTTCGTTCTTCCTTCTGTTCGGAGATTACCGAAT 6550
CAAAAAAATTTCAAGGAAACCGAAATCAAAAAAAAGAATAAAAAAAAAAT 6600
GATGAATTGAAAAGGTGGTATGGTGCACTCTCAGTACAATCTGCTCTGAT 6650
GCCGCATAGTTAAGCCAGCCCCGACACCCGCCAACACCCGCTGACGCGCC 6700
CTGACGGGCTTGTCTGCTCCCGGCATCCGCTTACAGACAAGCTGTGACCG 6750
TCTCCGGGAGCTGCATGTGTCAGAGGTTTTCACCGTCATCACCGAAACGC 6800
GCGAGACGAAAGGGCCTCGTGATACGCCTATTTTTATAGGTTAATGTCAT 6850
GATAATAATGGTTTCTTAGCAGGTTAGACTAACATGCAAAAAGTTTATAT 6900
TCATTTTTGTAGTTATTCTTGTTTTCATTAGGATGATTACACACTGTGGA 6950
TGTGGATTAATCGATCATTGTTAATTTAGTGTAGCTACTACCATTTCAAA 7000
CAAAACGTAAATTCTTGCAGTCGTACTGGATCTGTGAATCTATTAGTATA 7050
TATGAATTAAAGTAGCTTGTACATATTATTCTGTTGAATCATATCGCAGA 7100
GCATTGGTTGAAATCCAAAATATAAAAATGTAATATCACAAAAAATAATA 7150
CTAATTCTAACATTAATGGGTCAGATTTTTAGTGAATACTTAAATTTATA 7200
ATATCGTCTATTTAAGCTAGCAAATGGAACAACATTTAAAGTAAGAACAT 7250
CATATCTACATGAAAATGTATATTTCAATCTGACTAATAACGCAGAGCAC 7300
ATCTTTCAGTGATGTCTGTCACATGATCAAAAAGAATTGTATTTAAATAA 7350
TTCATAATAAAAGCTTAAAAAATTACAATAATGAAAATAAAGTAAATAAT 7400
GACATGGGTAAGAGTTCCGAAATAGAATCTTAGTGTACAAAACAAAAATT 7450
GCATCATTAGAGATCCCCACATTCAAATATGTATCCGCTCATGAGACAAT 7500
AACCCTGATAAATGCTTCAATAATATTGAAAAAGGAAGAGTATGAGTATT 7550
CAACATTTCCGTGTCGCCCTTATTCCCTTTTTTGCGGCATTTTGCCTTCC 7600
TGTTTTTGCTCACCCAGAAACGCTGGTGAAAGTAAAAGATGCTGAAGATC 7650

```
AGTTGGGTGCACGAGTGGGTTACATCGAACTGGATCTCAACAGCGGTAAG 7700
ATCCTTGAGAGTTTTCGCCCCGAAGAACGTTTTCCAATGATGAGCACTTT 7750
TAAAGTTCTGCTATGTGGCGCGGTATTATCCCGTATTGACGCCGGGCAAG 7800
AGCAACTCGGTCGCCGCATACACTATTCTCAGAATGACTTGGTTGAGTAC 7850
TCACCAGTCACAGAAAAGCATCTTACGGATGGCATGACAGTAAGAGAATT 7900
ATGCAGTGCTGCCATAACCATGAGTGATAACACTGCGGCCAACTTACTTC 7950
TGACAACGATCGGAGGACCGAAGGAGCTAACCGCTTTTTTGCACAACATG 8000
GGGGATCATGTAACTCGCCTTGATCGTTGGGAACCGGAGCTGAATGAAGC 8050
CATACCAAACGACGAGCGTGACACCACGATGCCTGTAGCAATGGCAACAA 8100
CGTTGCGCAAACTATTAACTGGCGAACTACTTACTCTAGCTTCCCGGCAA 8150
CAATTAATAGACTGGATGGAGGCGGATAAAGTTGCAGGACCACTTCTGCG 8200
CTCGGCCCTTCCGGCTGGCTGGTTTATTGCTGATAAATCTGGAGCCGGTG 8250
AGCGTGGGTCTCGCGGTATCATTGCAGCACTGGGGCCAGATGGTAAGCCC 8300
TCCCGTATCGTAGTTATCTACACGACGGGGAGTCAGGCAACTATGGATGA 8350
ACGAAATAGACAGATCGCTGAGATAGGTGCCTCACTGATTAAGCATTGGT 8400
AACTGTCAGACCAAGTTTACTCATATATACTTTAGATTGATTTAAAACTT 8450
CATTTTTAATTTAAAAGGATCTAGGTGAAGATCCTTTTTGATAATCTCAT 8500
GACCAAAATCCCTTAACGTGAGTTTTCGTTCCACTGAGCGTCAGACCCCG 8550
TAGAAAAGATCAAAGGATCTTC ••• 8572
```

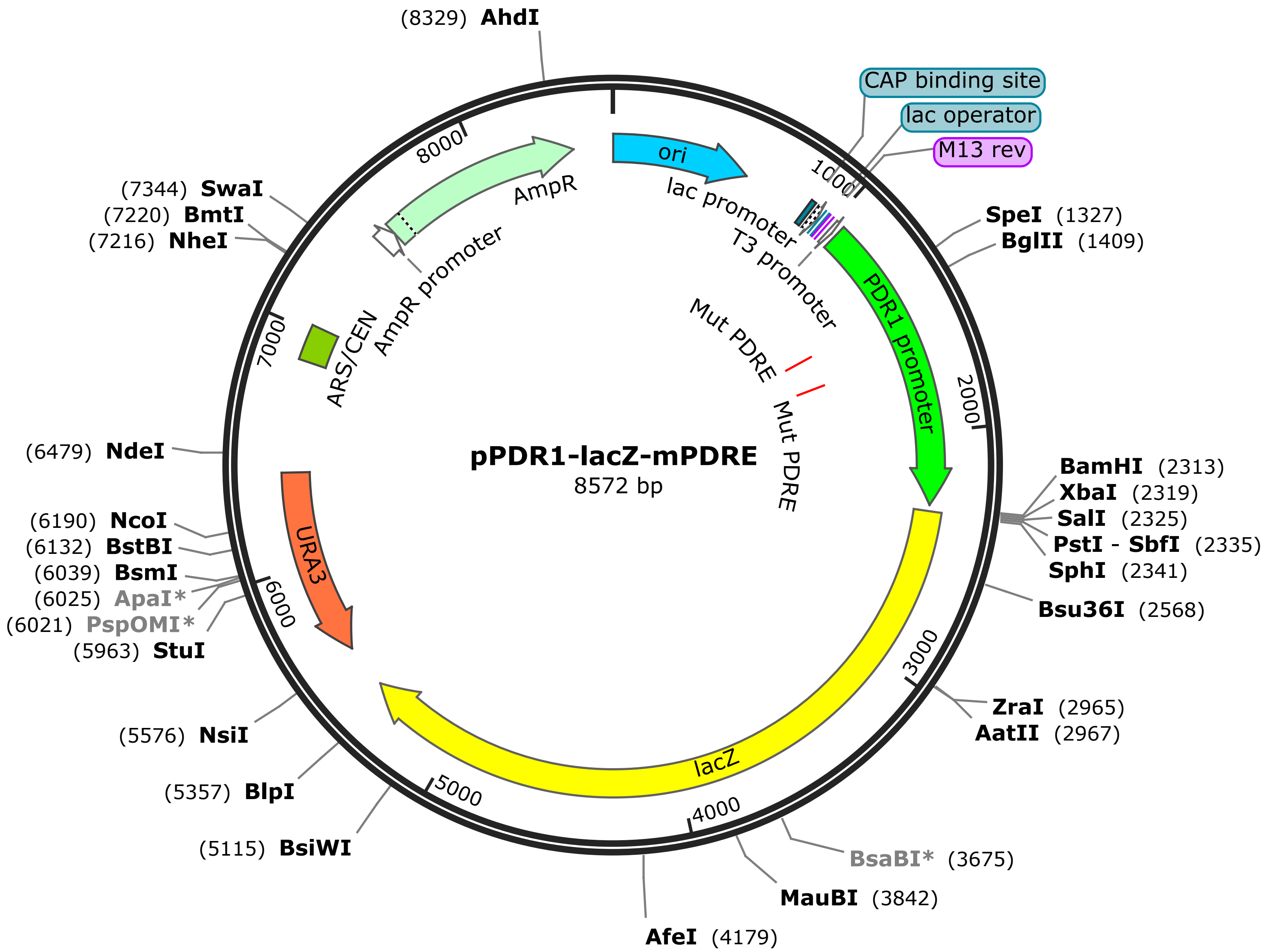
pPDR1-lacZ-mPDRE
8572 bp
(8329) AhdI
(7344) SwaI
(7220) BmtI
(7216) NheI
(6479) NdeI
(6190) NcoI
(6132) BstBI
(6039) BsmI
(6025) ApaI*
(6021) PspOMI*
(5963) StuI
(5576) NsiI
(5357) BlpI
(5115) BsiWI
AfeI (4179)
MauBI (3842)
BsaBI* (3675)
ZraI (2965)
AatII (2967)
Bsu36I (2568)
BamHI (2313)
XbaI (2319)
SalI (2325)
PstI - SbfI (2335)
SphI (2341)
SpeI (1327)
BglII (1409)
CAP binding site
lac operator
M13 rev
ori
lac promoter
T3 promoter
PDR1 promoter
Mut PDRE
Mut PDRE
lacZ
URA3
AmpR
AmpR promoter
ARS/CEN
1000
2000
3000
4000
5000
6000
7000
8000

•••
TTGAGATCCTTTTTTTCTGCGCGTAATCTGCTGCTTGCAAACAAAAAAAC 50
CACCGCTACCAGCGGTGGTTTGTTTGCCGGATCAAGAGCTACCAACTCTT 100
TTTCCGAAGGTAACTGGCTTCAGCAGAGCGCAGATACCAAATACTGTTCT 150
TCTAGTGTAGCCGTAGTTAGGCCACCACTTCAAGAACTCTGTAGCACCGC 200
CTACATACCTCGCTCTGCTAATCCTGTTACCAGTGGCTGCTGCCAGTGGC 250
GATAAGTCGTGTCTTACCGGGTTGGACTCAAGACGATAGTTACCGGATAA 300
GGCGCAGCGGTCGGGCTGAACGGGGGGTTCGTGCACACAGCCCAGCTTGG 350
AGCGAACGACCTACACCGAACTGAGATACCTACAGCGTGAGCTATGAGAA 400
AGCGCCACGCTTCCCGAAGGGAGAAAGGCGGACAGGTATCCGGTAAGCGG 450
CAGGGTCGGAACAGGAGAGCGCACGAGGGAGCTTCCAGGGGGAAACGCCT 500
GGTATCTTTATAGTCCTGTCGGGTTTCGCCACCTCTGACTTGAGCGTCGA 550
TTTTTGTGATGCTCGTCAGGGGGGCGGAGCCTATGGAAAAACGCCAGCAA 600
CGCGGCCTTTTTACGGTTCCTGGCCTTTTGCTGGCCTTTTGCTCACATGT 650
TCTTTCCTGCGTTATCCCCTGATTCTGTGGATAACCGTATTACCGCCTTT 700
GAGTGAGCTGATACCGCTCGCCGCAGCCGAACGACCGAGCGCAGCGAGTC 750
AGTGAGCGAGGAAGCGGAAGAGCGCCCAATACGCAAACCGCCTCTCCCCG 800
CGCGTTGGCCGATTCATTAATGCAGCTGGCACGACAGGTTTCCCGACTGG 850
AAAGCGGGCAGTGAGCGCAACGCAATTAATGTGAGTTAGCTCACTCATTA 900
GGCACCCCAGGCTTTACACTTTATGCTTCCGGCTCGTATGTTGTGTGGAA 950
TTGTGAGCGGATAACAATTTCACACAGGAAACAGCTATGACCATGATTAC 1000
GCCAAGCGCGCAATTAACCCTCACTAAAGGGAACAAAAGCTGGAGCTCGC 1050
ATTATCTAGGTGGCTGGGTGCCGGTAATCTGAGTTTGGATATATATATAT 1100
GCGTTTGATGAAGGCACAATGGTGGGGGTCCGATTTTAAAGATATTTTCC 1150
TTTTTTAGTTTTGTTGTTATTTTTTGATATAAGCCGAATGATCATCTGAA 1200
TCTAAATTTGAAATGAAAGACTTCGTTAGTTCTTTTCCTTTTTCAAGTGG 1250
GTATCTTTACACTACTGAGATGAAAGCAATAACTGTTGATTGTACCCATA 1300
CAGAAGAAAACTTAGAAAAATGAAATACTAGTACATTAAGAAAAAAACAG 1350
CAGAAAAAAAAAAGAAAAGCTGAAGCATTATTATACATCGTAACAAACAT 1400
TTCCTCATAGATCTAATCTTTTACTTCAGCAATTTCTAGTCTCTGCAACA 1450
AATTACACTAAAAAAATTACAAATCTTGCACGAGTATCATAACGATGCAG 1500
TGTCAAGGTCTGAGCATACTGAGGTTATTTTGTTGGCTTGTATTATCAGT 1550
TTCTTTTCTTTCTTCCTTTTTTGTTTTTTTTTTGTATTTCTTTTGGAGTT 1600
GTCCGTTTCCCGCAACCAGACAAAGACCCCAGTATTTGTAGACTCATTAA 1650
AAAAAGCAAATTGTGCTATATGGATTATTTCTCTGTGCTTCATTTTCTAC 1700
CTCGTAGATTAGGTTACGTTCAAATTTTAAGTGGCAAAACCATATCCTGA 1750
TCGTCAGTACTAAACCTGAGTTGTCCCGATGAGCCTCCTATTTTTTTTAA 1800
AGATGGCTTTTCCTATTGATCTAAGTGGATAAGCAGGTATTGCGTTGATC 1850
ATTATAATTGTGGGTAAAACTAGGTATTAGGTCCTTCTAATAGTCATCTT 1900
TATTGTTGTAAAAATTGTTTCGATGGCGTATTCATAGAATCCGAACATAT 1950
ATACCAACCAAATGGATTTTAAAGGAAAAAGCAAATACATTAAGGTCTGT 2000
TGTAGGAAAAGACCATATTGATTTATCTATTTAGTTGAGTAACAGAAATC 2050
ACATATAGTTTTAAATTTTTGGTAATCAAAGACACTTGAAACTATAAAAC 2100
ATTTGGTTGAAGAAGAAGTATTAACTTTGCTTGGAGAAAAGCGTGCCCAC 2150
ACTTGCTTGGTTTTCAGGATAATAACAAGCATAGAGGCGCTGTTATTACT 2200
TAAACAATTTTTAAGTAACACATTCAAACTTCCATTACTTCGTACCCCAT 2250
ATCGTATTGCCATTGTGATATGGAATTAGTGTTTTATTCTGCCTTTTTTT 2300
TTAGAATATATTGGATCCTCTAGAGTCGACCTGCAGGCATGCAAGCTTGC 2350
GATCCCGTCGTTTTACAACGTCGTGACTGGGAAAACCCTGGCGTTACCCA 2400
ACTTAATCGCCTTGCAGCACATCCCCCTTTCGCCAGCTGGCGTAATAGCG 2450
AAGAGGCCCGCACCGATCGCCCTTCCCAACAGTTGCGCAGCCTGAATGGC 2500
GAATGGCGCTTTGCCTGGTTTCCGGCACCAGAAGCGGTGCCGGAAAGCTG 2550

GCTGGAGTGCGATCTTCCTGAGGCCGATACTGTCGTCGTCCCCTCAAACT 2600
GGCAGATGCACGGTTACGATGCGCCCATCTACACCAACGTGACCTATCCC 2650
ATTACGGTCAATCCGCCGTTTGTTCCCACGGAGAATCCGACGGGTTGTTA 2700
CTCGCTCACATTTAATGTTGATGAAAGCTGGCTACAGGAAGGCCAGACGC 2750
GAATTATTTTTGATGGCGTTAACTCGGCGTTTCATCTGTGGTGCAACGGG 2800
CGCTGGGTCGGTTACGGCCAGGACAGTCGTTTGCCGTCTGAATTTGACCT 2850
GAGCGCATTTTTACGCGCCGGAGAAAACCGCCTCGCGGTGATGGTGCTGC 2900
GCTGGAGTGACGGCAGTTATCTGGAAGATCAGGATATGTGGCGGATGAGC 2950
GGCATTTTCCGTGACGTCTCGTTGCTGCATAAACCGACTACACAAATCAG 3000
CGATTTCCATGTTGCCACTCGCTTTAATGATGATTTCAGCCGCGCTGTAC 3050
TGGAGGCTGAAGTTCAGATGTGCGGCGAGTTGCGTGACTACCTACGGGTA 3100
ACAGTTTCTTTATGGCAGGGTGAAACGCAGGTCGCCAGCGGCACCGCGCC 3150
TTTCGGCGGTGAAATTATCGATGAGCGTGGTGGTTATGCCGATCGCGTCA 3200
CACTACGTCTGAACGTCGAAAACCCGAAACTGTGGAGCGCCGAAATCCCG 3250
AATCTCTATCGTGCGGTGGTTGAACTGCACACCGCCGACGGCACGCTGAT 3300
TGAAGCAGAAGCCTGCGATGTCGGTTTCCGCGAGGTGCGGATTGAAAATG 3350
GTCTGCTGCTGCTGAACGGCAAGCCGTTGCTGATTCGAGGCGTTAACCGT 3400
CACGAGCATCATCCTCTGCATGGTCAGGTCATGGATGAGCAGACGATGGT 3450
GCAGGATATCCTGCTGATGAAGCAGAACAACTTTAACGCCGTGCGCTGTT 3500
CGCATTATCCGAACCATCCGCTGTGGTACACGCTGTGCGACCGCTACGGC 3550
CTGTATGTGGTGGATGAAGCCAATATTGAAACCCACGGCATGGTGCCAAT 3600
GAATCGTCTGACCGATGATCCGCGCTGGCTACCGGCGATGAGCGAACGCG 3650
TAACGCGAATGGTGCAGCGCGATCGTAATCACCCGAGTGTGATCATCTGG 3700
TCGCTGGGGAATGAATCAGGCCACGGCGCTAATCACGACGCGCTGTATCG 3750
CTGGATCAAATCTGTCGATCCTTCCCGCCCGGTGCAGTATGAAGGCGGCG 3800
GAGCCGACACCACGGCCACCGATATTATTTGCCCGATGTACGCGCGCGTG 3850
GATGAAGACCAGCCCTTCCCGGCTGTGCCGAAATGGTCCATCAAAAAATG 3900
GCTTTCGCTACCTGGAGAGACGCGCCCGCTGATCCTTTGCGAATACGCCC 3950
ACGCGATGGGTAACAGTCTTGGCGGTTTCGCTAAATACTGGCAGGCGTTT 4000
CGTCAGTATCCCCGTTTACAGGGCGGCTTCGTCTGGGACTGGGTGGATCA 4050
GTCGCTGATTAAATATGATGAAAACGGCAACCCGTGGTCGGCTTACGGCG 4100
GTGATTTTGGCGATACGCCGAACGATCGCCAGTTCTGTATGAACGGTCTG 4150
GTCTTTGCCGACCGCACGCCGCATCCAGCGCTGACGGAAGCAAAACACCA 4200
GCAGCAGTTTTTCCAGTTCCGTTTATCCGGGCAAACCATCGAAGTGACCA 4250
GCGAATACCTGTTCCGTCATAGCGATAACGAGCTCCTGCACTGGATGGTG 4300
GCGCTGGATGGTAAGCCGCTGGCAAGCGGTGAAGTGCCTCTGGATGTCGC 4350
TCCACAAGGTAAACAGTTGATTGAACTGCCTGAACTACCGCAGCCGGAGA 4400
GCGCCGGGCAACTCTGGCTCACAGTACGCGTAGTGCAACCGAACGCGACC 4450
GCATGGTCAGAAGCCGGGCACATCAGCGCCTGGCAGCAGTGGCGTCTGGC 4500
GGAAAACCTCAGTGTGACGCTCCCCGCCGCGTCCCACGCCATCCCGCATC 4550
TGACCACCAGCGAAATGGATTTTTGCATCGAGCTGGGTAATAAGCGTTGG 4600
CAATTTAACCGCCAGTCAGGCTTTCTTTCACAGATGTGGATTGGCGATAA 4650
AAAACAACTGCTGACGCCGCTGCGCGATCAGTTCACCCGTGCACCGCTGG 4700
ATAACGACATTGGCGTAAGTGAAGCGACCCGCATTGACCCTAACGCCTGG 4750
GTCGAACGCTGGAAGGCGGCGGGCCATTACCAGGCCGAAGCAGCGTTGTT 4800
GCAGTGCACGGCAGATACACTTGCTGATGCGGTGCTGATTACGACCGCTC 4850
ACGCGTGGCAGCATCAGGGGAAAACCTTATTTATCAGCCGGAAAACCTAC 4900
CGGATTGATGGTAGTGGTCAAATGGCGATTACCGTTGATGTTGAAGTGGC 4950
GAGCGATACACCGCATCCGGCGCGGATTGGCCTGAACTGCCAGCTGGCGC 5000
AGGTAGCAGAGCGGGTAAACTGGCTCGGATTAGGGCCGCAAGAAAACTAT 5050
CCCGACCGCCTTACTGCCGCCTGTTTTGACCGCTGGGATCTGCCATTGTC 5100

AGACATGTATATCCCGTACGTCTTCCCGAGCGAAAACGGTCTGCGCTGCG 5150
GGACGCGCGAATTGAATTATGGCCCACACCAGTGGCGCGGCGACTTCCAG 5200
TTCAACATCAGCCGCTACAGTCAACAGCAACTGATGGAAACCAGCCATCG 5250
CCATCTGCTGCACGCGGAAGAAGGCACATGGCTGAATATCGACGGTTTCC 5300
ACATGGGGATTGGTGGCGACGACTCCTGGAGCCCGTCAGTATCGGCGGAA 5350
TTACAGCTGAGCGCCGGTCGCTACCATTACCAGTTGGTCTGGTGTCAAAA 5400
ATAATAATAACCGGGCAGGCCATGTCTGCCCGTATTTCGCGTAAGGAAAT 5450
CCATTATGTACTATTTCCTGATGCGGTATTTTCTCCTTACGCATCTGTGC 5500
GGTATTTCACACCGCATAGGGTAATAACTGATATAATTAAATTGAAGCTC 5550
TAATTTGTGAGTTTAGTATACATGCATTTACTTATAATACAGTTTTTTAG 5600
TTTTGCTGGCCGCATCTTCTCAAATATGCTTCCCAGCCTGCTTTTCTGTA 5650
ACGTTCACCCTCTACCTTAGCATCCCTTCCCTTTGCAAATAGTCCTCTTC 5700
CAACAATAATAATGTCAGATCCTGTAGAGACCACATCATCCACGGTTCTA 5750
TACTGTTGACCCAATGCGTCTCCCTTGTCATCTAAACCCACACCGGGTGT 5800
CATAATCAACCAATCGTAACCTTCATCTCTTCCACCCATGTCTCTTTGAG 5850
CAATAAAGCCGATAACAAAATCTTTGTCGCTCTTCGCAATGTCAACAGTA 5900
CCCTTAGTATATTCTCCAGTAGATAGGGAGCCCTTGCATGACAATTCTGC 5950
TAACATCAAAAGGCCTCTAGGTTCCTTTGTTACTTCTTCTGCCGCCTGCT 6000
TCAAACCGCTAACAATACCTGGGCCCACCACACCGTGTGCATTCGTAATG 6050
TCTGCCCATTCTGCTATTCTGTATACACCCGCAGAGTACTGCAATTTGAC 6100
TGTATTACCAATGTCAGCAAATTTTCTGTCTTCGAAGAGTAAAAAATTGT 6150
ACTTGGCGGATAATGCCTTTAGCGGCTTAACTGTGCCCTCCATGGAAAAA 6200
TCAGTCAAGATATCCACATGTGTTTTTAGTAAACAAATTTTGGGACCTAA 6250
TGCTTCAACTAACTCCAGTAATTCCTTGGTGGTACGAACATCCAATGAAG 6300
CACACAAGTTTGTTTGCTTTTCGTGCATGATATTAAATAGCTTGGCAGCA 6350
ACAGGACTAGGATGAGTAGCAGCACGTTCCTTATATGTAGCTTTCGACAT 6400
GATTTATCTTCGTTTCGGTTTTTGTTCTGTGCAGTTGGGTTAAGAATACT 6450
GGGCAATTTCATGTTTCTTCAACACTACATATGCGTATATATACCAATCT 6500
AAGTCTGTGCTCCTTCCTTCGTTCTTCCTTCTGTTCGGAGATTACCGAAT 6550
CAAAAAAATTTCAAGGAAACCGAAATCAAAAAAAAGAATAAAAAAAAAAT 6600
GATGAATTGAAAAGGTGGTATGGTGCACTCTCAGTACAATCTGCTCTGAT 6650
GCCGCATAGTTAAGCCAGCCCCGACACCCGCCAACACCCGCTGACGCGCC 6700
CTGACGGGCTTGTCTGCTCCCGGCATCCGCTTACAGACAAGCTGTGACCG 6750
TCTCCGGGAGCTGCATGTGTCAGAGGTTTTCACCGTCATCACCGAAACGC 6800
GCGAGACGAAAGGGCCTCGTGATACGCCTATTTTTATAGGTTAATGTCAT 6850
GATAATAATGGTTTCTTAGCAGGTTAGACTAACATGCAAAAAGTTTATAT 6900
TCATTTTTGTAGTTATTCTTGTTTTCATTAGGATGATTACACACTGTGGA 6950
TGTGGATTAATCGATCATTGTTAATTTAGTGTAGCTACTACCATTTCAAA 7000
CAAAACGTAAATTCTTGCAGTCGTACTGGATCTGTGAATCTATTAGTATA 7050
TATGAATTAAAGTAGCTTGTACATATTATTCTGTTGAATCATATCGCAGA 7100
GCATTGGTTGAAATCCAAAATATAAAAATGTAATATCACAAAAAATAATA 7150
CTAATTCTAACATTAATGGGTCAGATTTTTAGTGAATACTTAAATTTATA 7200
ATATCGTCTATTTAAGCTAGCAAATGGAACAACATTTAAAGTAAGAACAT 7250
CATATCTACATGAAAATGTATATTTCAATCTGACTAATAACGCAGAGCAC 7300
ATCTTTCAGTGATGTCTGTCACATGATCAAAAAGAATTGTATTTAAATAA 7350
TTCATAATAAAAGCTTAAAAAATTACAATAATGAAAATAAAGTAAATAAT 7400
GACATGGGTAAGAGTTCCGAAATAGAATCTTAGTGTACAAAACAAAAATT 7450
GCATCATTAGAGATCCCCACATTCAAATATGTATCCGCTCATGAGACAAT 7500
AACCCTGATAAATGCTTCAATAATATTGAAAAAGGAAGAGTATGAGTATT 7550
CAACATTTCCGTGTCGCCCTTATTCCCTTTTTTGCGGCATTTTGCCTTCC 7600
TGTTTTTGCTCACCCAGAAACGCTGGTGAAAGTAAAAGATGCTGAAGATC 7650

```
AGTTGGGTGCACGAGTGGGTTACATCGAACTGGATCTCAACAGCGGTAAG    7700
ATCCTTGAGAGTTTTCGCCCCGAAGAACGTTTTCCAATGATGAGCACTTT    7750
TAAAGTTCTGCTATGTGGCGCGGTATTATCCCGTATTGACGCCGGGCAAG    7800
AGCAACTCGGTCGCCGCATACACTATTCTCAGAATGACTTGGTTGAGTAC    7850
TCACCAGTCACAGAAAAGCATCTTACGGATGGCATGACAGTAAGAGAATT    7900
ATGCAGTGCTGCCATAACCATGAGTGATAACACTGCGGCCAACTTACTTC    7950
TGACAACGATCGGAGGACCGAAGGAGCTAACCGCTTTTTTGCACAACATG    8000
GGGGATCATGTAACTCGCCTTGATCGTTGGGAACCGGAGCTGAATGAAGC    8050
CATACCAAACGACGAGCGTGACACCACGATGCCTGTAGCAATGGCAACAA    8100
CGTTGCGCAAACTATTAACTGGCGAACTACTTACTCTAGCTTCCCGGCAA    8150
CAATTAATAGACTGGATGGAGGCGGATAAAGTTGCAGGACCACTTCTGCG    8200
CTCGGCCCTTCCGGCTGGCTGGTTTATTGCTGATAAATCTGGAGCCGGTG    8250
AGCGTGGGTCTCGCGGTATCATTGCAGCACTGGGGCCAGATGGTAAGCCC    8300
TCCCGTATCGTAGTTATCTACACGACGGGGAGTCAGGCAACTATGGATGA    8350
ACGAAATAGACAGATCGCTGAGATAGGTGCCTCACTGATTAAGCATTGGT    8400
AACTGTCAGACCAAGTTTACTCATATATACTTTAGATTGATTTAAAACTT    8450
CATTTTTAATTTAAAAGGATCTAGGTGAAGATCCTTTTTGATAATCTCAT    8500
GACCAAAATCCCTTAACGTGAGTTTTCGTTCCACTGAGCGTCAGACCCCG    8550
TAGAAAAGATCAAAGGATCTTC ••• 8572
```
